# Treg- and CD8-Specific PD-1 Deficiency Reveals That CD8⁺ T Cell-Intrinsic Signaling Uniquely Restrains Pressure Overload Inflammation and Heart Failure

**DOI:** 10.64898/2026.11.13.766421

**Authors:** Xiaochen He, Xinyu Weng, Lihong Pan, Ziru Niu, Umesh Bhattarai, Rui Xu, Dongzhi Wang, Heng Zeng, Jian-Xiong Chen, John S. Clemmer, Ajay Singh, Babbette LaMarca, Bruce R Blazar, Michael Hall, Yingjie Chen

## Abstract

**Aims:** PD-1 signaling limits cardiac remodeling and heart failure (HF). Whether this protection depends on PD-1 signaling intrinsic to CD8⁺ T cells or on regulation by other immune populations, including regulatory T cells (Tregs), remains unclear.

**Methods and results:** Using transverse aortic constriction (TAC), we studied global PD-1-knockout (KO) mice with or without CD8⁺ T cell depletion, CD8⁺ T cell-specific (PD1KO^CD8^), and Treg-specific PD-1-KO mice. We assessed left ventricular (LV) remodeling, genomic profiles, immune infiltration, and the therapeutic impact of C-X-C motif chemokine receptor 3 (CXCR3) blockade. CD8⁺ T cell depletion abolished TAC-induced left ventricular (LV) hypertrophy, fibrosis, and dysfunction of global PD-1-KO mice. PD1KO^CD8^ mice had normal baseline cardiac function but, after TAC, showed reduced survival, worse LV dysfunction and fibrosis, and greater immune cell infiltration. CD8⁺ T cells from PD1KO^CD8^ mice showed increased activation and proliferation, with increased production of cytotoxic molecules, such as IFN-γ⁺ and granzyme A, both at baseline and after TAC. Post-TAC transcriptomics revealed enhanced inflammatory pathways and suppressed metabolic pathways in PD1KO^CD8^ hearts. TAC also increased LV CXCR3^+^CD8^+^ T cell accumulation and CXCL9/CXCL10 expression in PD1KO^CD8^ mice. CXCR3 neutralizing antibody treatment reduced LV CD8^+^ T cell accumulation and attenuated cardiac dysfunction. In contrast, Treg-specific PD-1 deficiency did not alter baseline or TAC-induced cardiac remodeling.

**Conclusions:** CD8⁺ T cell-intrinsic PD-1 restrains TAC-induced HF, whereas Treg-intrinsic PD-1 is dispensable. CXCR3-dependent recruitment drives pathological CD8⁺ T cell accumulation, identifying PD-1/CXCR3 axis as a key regulator of CD8⁺ T cell-mediated cardiac injury.

## 1. Introduction

Immune checkpoint inhibitors (ICIs) targeting the programmed cell death protein 1 (PD-1) pathway have revolutionized the treatment of multiple malignancies and substantially improved survival across a broad range of cancers ^1^. However, the widespread clinical success of these therapies has unveiled significant cardiovascular liabilities, primarily manifesting as immune-related adverse events such as fulminant myocarditis and the accelerated progression of heart failure (HF) ^2, 3^. Crucially, ICI-induced cardiac injury and dysfunction often occur in people with preexisting cardiovascular diseases, such as coronary artery disease, valvular disease, and hypertension ^4^. Hypertension and other conditions that increase left ventricular (LV) afterload impose chronic pressure overload on the myocardium, a pathological stimulus that can be modeled experimentally using transverse aortic constriction (TAC). Within this pathological context, CD8^+^ T cells are increasingly recognized as an important immune cell subset in promoting ICI-induced cardiac injury and HF ^5^. Moreover, CD8^+^ T cells play an important role in driving systolic overload-induced cardiac inflammation and HF, even in the absence of ICI therapies ^6, 7^. The blockade of PD-1 disrupts fundamental immune tolerance checkpoints, unleashing hyperactive, cytotoxic CD8+ T cell responses that, while highly effective against tumor antigens, can inadvertently target the myocardium and drive severe, organ-threatening cardiac inflammation ^8^.

Despite the clinical importance of PD-1-directed therapies, the endogenous role of the PD-1 signaling in regulating immune responses during pressure overload remains incompletely understood. While clinical observations suggest a link between PD-1 disruption and cardiac dysfunction, the specific immunological mechanisms that govern tolerance during systolic overload remain poorly defined. In particular, it is unclear whether the protective effects of PD-1 depend primarily on inhibitory signaling intrinsic to CD8⁺ T cells or on PD-1-dependent regulation by other immune populations. Regulatory T cells (Tregs) are of particular interest because they suppress excessive immune activation and protect against pressure overload-induced cardiac inflammation and HF development ^9^. Resolving this ambiguity is essential for understanding how immune networks govern adverse cardiac remodeling and for developing safer, more targeted immunomodulatory strategies.

To address these fundamental knowledge gaps, we hypothesized that PD-1 signaling intrinsic to CD8^+^ T cells, rather than extrinsic regulation by Tregs, is required to restrict pathological CD8⁺ T-cell activation and myocardial accumulation during chronic systolic overload. To test this hypothesis and definitively isolate the source of immune regulation, the current study utilized a comparative genetic approach utilizing global PD-1 knockout (KO) mice, together with Treg-specific PD-1 KO (PD1KO^FoxP3^) and CD8^+^ T cell-specific PD-1 KO (PD1KO^CD8^) mice subjected to TAC. These complementary genetic models were combined with CD8⁺ T-cell depletion and CXCR3 blockade to determine the contribution of CD8^+^ T cell-intrinsic PD-1 signaling to pressure overload-induced cardiac remodeling and to identify mechanisms governing CD8⁺ T-cell tolerance and pathological CD8⁺ T-cell recruitment to the myocardium.

## 2. Methods

### 2.1 Data Availability

The data that support the findings of this study are available from the corresponding author upon reasonable request. Detailed experimental methods are available in the Supplementary Materials.

### 2.2 Study Approval

All animal protocols were approved by the Institutional Animal Care and Use Committee (IACUC) of the University of Mississippi Medical Center (Protocol ID: 2022-1254) and were consistent with the National Institutes of Health Guide for the Care and Use of Laboratory Animals (8^th^ edition, Revised 2011). All animals were fed with laboratory standard chow and water ad libitum and housed in individually ventilated cages in the Center for Comparative Research at the University of Mississippi Medical Center.

### 2.3 Mice

Wild-type (WT, C57BL/6J) and global PD1KO mice (Strain #:028276) were purchased from the Jackson Laboratories. PD1KO^CD8^ mice were generated using the Cre-LoxP system and PD1^flox/flox^ mice as depicted in **Supplementary Figure S3**. Exons 2 and 3 of the PD-1 gene are flanked by LoxP sites, priming them for subsequent deletion by Cre recombinase. C57BL/6-Tg(Cd8a-cre)1Itan/J transgenic mice (Strain #: 008766) were purchased from the Jackson Laboratories. PD1^flox/flox^ mice were crossbred with CD8-Cre transgenic mice. The resulting PD1^flox/−^CD8-Cre^+/-^ heterozygous mutants were then mated with PD1^flox/flox^ mice to obtain CD8-specific PD1 KO mice (PD1^flox/flox^CD8-Cre^+/-^, PD1KO^CD8^). WT littermates (PD1^flox/flox^CD8-Cre^-/-^, PD1^f/f^ wt) were used as controls. 10-to 12-week-old male and female mice were used in the experiments as indicated. Genotyping was performed by tail DNA PCR analysis. Primer sequences used for genotyping floxed PD-1 allele were listed in the **Supplementary Table S11**. PCR products were analyzed on 1.5% tris-acetate-EDTA (TAE) agarose gels (**Supplementary Figure S10**). FoxP3^Cre^ mice (Jackson Laboratories, Strain #: 016959) were used to generate PD1KO^FoxP3^ mice. Primer sequences used for genotyping were listed in the **Supplementary Table S11**. PCR products were analyzed on 1.5% tris-acetate-EDTA (TAE) agarose gels.

### 2.4 Transvers Aortic Constriction Surgery Procedure

10-12 weeks old male and female mice were subjected to TAC or sham surgery. LV pressure overload was induced by TAC surgery for an indicated period as previously described ^10^. Briefly, the mice were anesthetized with a single intraperitoneal injection of ketamine (100 mg/kg) and xylazine (10 mg/kg). A longitudinal midline cervical incision was made to expose the proximal portion of the sternum, followed by a partial thoracotomy. The transverse aortic arch was exposed after gentle separation of the thymus and fat tissue. A small piece of 6-0 silk suture was placed between the innominate and the left common carotid arteries. The degree of aortic constriction induced by ligation was controlled by a blunt 27-gauge needle. The sham procedure was identical, except that there was no ligation of the aorta. Mice were given Buprenorphine Base in Polymer (1mg/kg) via subcutaneous injection shortly before waking up. At the end of the study, cardiac function was assessed by echocardiography, and the animals were euthanized by being given an overdose of ketamine (300 mg/kg) and xylazine (30 mg/kg) intraperitoneally, followed by exsanguination, cardiac perfusion with ice-cold phosphate-buffered saline, tissue collection, and subsequent analysis, to ensure death. All procedures were performed in strict accordance with the AVMA Guidelines for the Euthanasia of Animals and approved by the IACUC of the University of Mississippi Medical Center (Protocol ID: 2022-1254).

### 2.5 Statistical analysis

Data are presented as mean ± SEM. The assumptions of normality in both comparison groups were determined by the Shapiro-Wilk test. For a normal distribution, two-group comparisons were performed with an unpaired Student’s *t* test, while comparisons across >2 groups were assessed using one-way ANOVA followed by Tukey’s multiple comparisons test. For non-normally distributed data, two-group comparisons were conducted using the Mann– Whitney U test, and multiple-group comparisons were analyzed with the Kruskal–Wallis test followed by Dunn’s multiple comparisons test. When analyzing two pairs of groups, a two-way ANOVA with Tukey’s post hoc multiple comparisons test was applied. Statistical differences between survival rates of Kaplan-Meier curves were compared with the log-rank test. All statistical analyses were performed using GraphPad Prism 10 (GraphPad Software, Boston, MA). The applied tests are indicated in the figure legends, and p < 0.05 was considered statistically significant. *p < 0.05, **p < 0.01, and ***p < 0.001.

## 3. Results

### 3.1 Targeted Depletion of CD8⁺ T Cells Rescues Cardiac Dysfunction and Pathological Remodeling in PD-1 Deficient Mice

To investigate whether the exacerbated pathological remodeling observed in PD-1-deficient mice is directly mediated by CD8⁺ T cells, we performed targeted *in vivo* depletion of this specific immune population. Wild-type (WT) and global PD-1 KO mice were subjected to sham or TAC surgery and were treated with either an anti-CD8 depleting antibody or an IgG isotype control over a 28-day period (**Figure 1A**).

**Figure 1.**
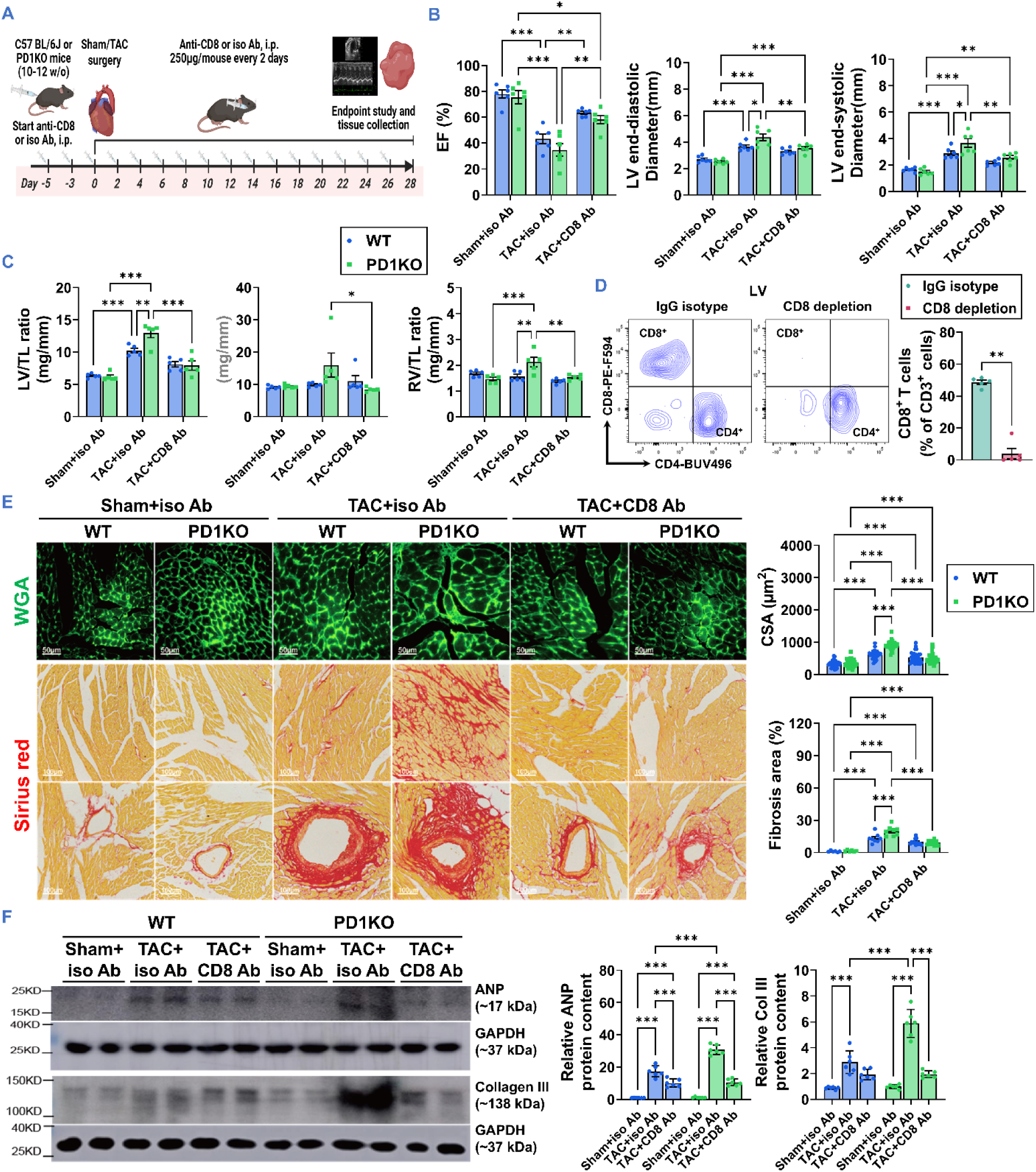
CD8^+^ T cell depletion attenuates cardiac dysfunction and pathological remodeling in PD-1 KO mice following TAC. **(A)** Experimental schematic. C57BL/6J (WT) and PD-1 KO mice underwent Sham or TAC surgery at day 0. Mice received intraperitoneal (i.p.) injections of either an anti-CD8 depleting antibody or an isotype control (Iso Ab) starting at day-5 and continuing every 2 days until day 28 at endpoint. **(B)** Quantitative analysis of echocardiographic parameters at day 28 post-TAC, including ejection fraction (EF%) and left ventricular (LV) internal dimension at systole and diastole. **(C)** Assessment of cardiac hypertrophy via heart weight-to-tibial length ratio (LV/TL), pulmonary congestion via lung weight-to-tibial length ratio (Lung/TL), and right ventricular weight-to-tibial length ratio (RV/TL). **(D)** Representative flow cytometry contour plots and analysis identifying CD8^+^ and CD4^+^ T cell populations in hearts on day 7 after TAC with anti-CD8 treatment, showing that CD8^+^ T cells were nearly completely depleted. **(E)** Representative histological images and quantitative analysis of LV sections. Top row: Wheat Germ Agglutinin (WGA) staining (green) to delineate cardiomyocyte membranes. Scale bar = 50 μm. Middle and bottom rows: Sirius red staining to visualize interstitial and perivascular collagen deposition, respectively. Scale bar = 100 μm. **(F)** Representative Western blots and densitometric quantitative analysis of the protein expression of profibrotic markers Fibronectin and Collagen I in cardiac tissue lysates. GAPDH served as the loading control. Data are presented as mean ± SEM. Statistical significance was determined by a two-way ANOVA with Tukey’s multiple comparisons test for panels A-C and E-F. Mann-Whitney test was used for panel (D). *p < 0.05, **p < 0.01, ***p < 0.001. n = 5-6 mice per group.

Echocardiographic assessment at the study endpoint revealed that while PD-1 KO mice treated with the isotype control exhibited severe LV systolic dysfunction following TAC, the systemic depletion of CD8⁺ T cells markedly preserved cardiac function. Specifically, anti-CD8 treatment in the KO cohort significantly restored LV ejection fraction (LVEF%) and fractional shortening (LVFS%) and robustly attenuated LV chamber dilation at end-systole and end-diastole compared with isotype-treated controls (**Figure 1B**). Consistent with this functional preservation, gravimetric analyses demonstrated that CD8⁺ T cell depletion significantly blunted the exaggerated cardiac hypertrophy (LV/TL and RV/TL ratios) and pulmonary congestion (Lung/TL ratio) typically induced by pressure overload in the absence of PD-1 (**Figure 1C**). As anticipated, an effective depletion of CD8^+^ T cells *in vivo* by anti-CD8 antibody in LV (**Figure 1D**), spleen (**Supplementary Figure S1**), and lymph node (**Supplementary S2D**) was confirmed by flow cytometry.

The structural and phenotypic benefits of CD8⁺ T cell depletion were further confirmed at the cellular and molecular levels. Histological evaluation via Wheat Germ Agglutinin (WGA) staining revealed a significant reduction in TAC-induced cardiomyocyte hypertrophy in anti-CD8 antibody-treated PD1 KO mice compared to their isotype-treated counterparts (**Figure 1E**, top). These changes were corroborated with the immunoblot analysis of atrial natriuretic peptide (ANP), a molecular marker of cardiac stress and hypertrophy (**Figure 1F**). Moreover, Sirius red staining revealed profound attenuation of both interstitial and perivascular collagen deposition, indicating suppression of extracellular matrix expansion (**Figure 1E**, middle and bottom). This reduction in pathological fibrosis was accompanied by Western blot analysis of Collagen III, a profibrotic marker, which showed that the elevated expression of Collagen III in the heart following TAC was significantly reduced following CD8⁺ T cell depletion (**Figure 1F**).

Because TAC-induced LV immune cell accumulation typically peaks at 7 to 10 days post-surgery, we evaluated a separate cohort of global PD-1 KO mice at 7 days post-TAC to capture the early inflammatory phase (**Supplementary Figure S2**). Consistent with the 4-week endpoint, CD8⁺ T cell depletion significantly preserved early systolic function (EF and FS) compared to IgG-treated controls (**Supplementary Figure S2A**). LV dimensions at end-systole and end-diastole were also significantly reduced in CD8⁺ T cell-depleted mice compared to IgG-treated controls (**Supplementary Figure S2B**). Furthermore, early LV hypertrophy and pulmonary congestion (Lung/TL) were significantly attenuated by CD8⁺ T cell depletion, though right ventricular remodeling (RV/TL) had not yet significantly diverged between the groups at this early stage (**Supplementary Figure S2C**). Crucially, immunofluorescence staining confirmed that early anti-CD8 treatment not only abolished CD8⁺ T cell presence within the myocardium (**Supplementary Figure S2E**) and lung (**Supplementary Figure S2F**), but also drove a marked secondary reduction in overall CD45⁺ leukocyte infiltration in the LV (**Supplementary Figure S2E**).

Collectively, these findings establish that CD8⁺ T cells are the primary cellular drivers of the exacerbated HF phenotype observed in PD-1 deficiency. Depleting this specific immune compartment is sufficient to rescue cardiac systolic function, limit pathological cardiomyocyte hypertrophy, and suppress adverse fibrotic remodeling during chronic systolic overload.

### 3.2 Cell-intrinsic PD-1 signaling in CD8⁺ T cells has no detectable effect on cardiopulmonary structure and function in both male and female mice under control conditions

To definitively determine whether the cardioprotective role of the PD-1 axis is governed by a requirement for cell-intrinsic self-tolerance within the CD8⁺ T cell compartment, we generated and evaluated our novel CD8⁺ T cell-specific PD-1 KO (PD1KO^CD8^) mouse strain alongside wild-type (PD1^f/f^ wt) littermates (**Supplementary Figure S3**). PD-1 deletion in CD8+ T cells was confirmed in both male and female PD1KO^CD8^ mice (**Supplementary Figure S4**). Under control conditions, there were no functional, structural, or pathological differences between PD1KO^CD8^ mice and WT littermates in either sex (**Supplemental Figures S5 and S6**). Both male and female PD1KO^CD8^ mice exhibited systolic function comparable to that of age-and sex-matched PD1^f/f^ wt littermates, as evidenced by similar ejection fraction and fractional shortening (**Supplemental Figures S5A and S5C**). In addition, LV/TL, Lung/TL, and RV/TL ratios are not different from the PD1^f/f^ wt control littermates (**Supplemental Figures S5B and S5D**). However, there was a small but significant increase in the number of infiltrating CD45^+^ leukocytes and CD8^+^ cytotoxic T cells in the LV section (**Supplemental Figure S7A and S7B**), but no difference in the RV sections (**Supplemental Figure S7C and S7D**). There were no functional, structural, or pathological differences in the pulmonary tissue between PD1KO^CD8^ mice and PD1^f/f^ wt littermates, although there was a small but significant increase in the number of infiltrating CD8^+^ cytotoxic T cells (**Supplemental Figure S8**).

### 3.3 Cell-Intrinsic PD-1 Restrains CD8⁺ T Cell Proliferation, Memory Differentiation, and Cytotoxic Molecule Productions at Baseline

To determine the impact of CD8-specific PD-1 deletion on systemic and tissue-resident immune profiles, we first assessed baseline leukocyte populations in peripheral blood mononuclear cells (PBMCs), hearts, and lungs. Flow cytometric profiling of cardiac CD45⁺ cells revealed no significant baseline differences in major immune subsets—including total T cells, macrophages, natural killer (NK) cells, B cells, neutrophils, and dendritic cells—between PD-1^f/f^ wt and PD1KO^CD8^ mice of either sex (**Supplementary Figure S9A, S9C, S9E, and S10A**).

Similarly, the relative percentages of CD8⁺ and CD4⁺ T cells within the CD3⁺ compartment remained unchanged in both PBMCs and cardiac tissues (**Supplementary Figure S9B, S9D, and Supplementary Figure S10B**). In the lungs, however, PD1KO^CD8^ mice exhibited a significant increase in the proportion of CD8⁺ T cells and a reciprocal decrease in CD4⁺ T cells (**Supplementary Figure S9F**).

Given the differences in cardiac remodeling and immune infiltration observed previously, we next sought to characterize the baseline activation status of these T cell populations. Phenotypic analysis of the CD8⁺ T cell compartment revealed a significant shift toward an effector memory phenotype in PD1KO^CD8^ mice. Compared to PD-1^f/f^ wt controls, PD1KO^CD8^ mice demonstrated a marked increase in the fraction of effector memory CD8⁺ T cells (**Figure 2A, 2B, 2C**) and a corresponding reduction in naïve CD8⁺ T cells across PBMCs, cardiac, and pulmonary tissues (**Supplementary Figure S11A, S12A, S12C, and S14A**). The proportion of central memory CD8⁺ T cells remained unchanged. In contrast, the distribution of naïve, effector memory, and central memory subsets within the CD4⁺ T cell compartment was completely unaffected by CD8-specific PD-1 deletion across all evaluated tissues (**Supplementary Figure S11D, S12B, S12D, S14B, and S14C**).

**Figure 2.**
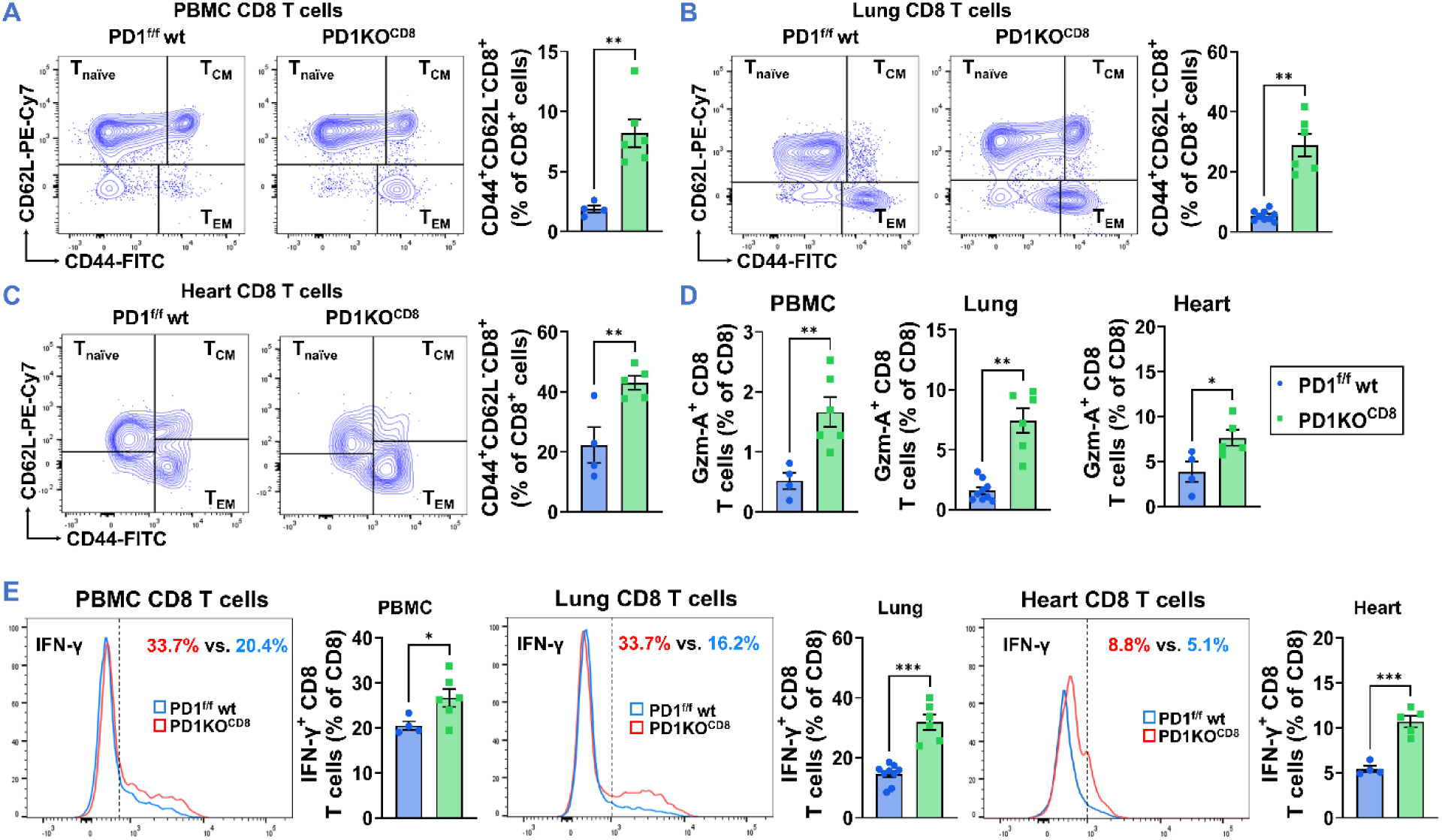
CD8-specific PD-1 deficiency significantly increases systemic CD8^+^ T cell activation and enhanced intracellular IFN-γ and Granzyme A production. (A–C) Representative flow cytometry contour plots and quantitative analysis of CD8^+^ T cell subsets isolated from PBMC (blood), lungs, and hearts (female) of PD1^f/f^ wt (blue) and PD1KO^CD8^ (green) mice. Subsets are phenotypically defined by surface marker expression as naïve (T_naïve_), central memory (T_CM_), and effector memory (T_EM_). The data demonstrate a significant phenotypic shift toward the activated T_EM_ compartment in PD1KO^CD8^ mice across all evaluated tissues. **(D)** Summarized quantification of the cytotoxic effector molecule Granzyme A expression within the CD8^+^ T cell populations from the aforementioned tissues (PBMC, lungs, and hearts), indicating heightened cytotoxic potential following PD-1 ablation. **(E)** Representative flow cytometry histograms and corresponding summarized data depicting the intracellular production of the pro-inflammatory cytokine IFN-γ in CD8^+^ T cells from PBMC, lungs, and hearts (female). The percentages of IFN-γ^+^ cells are markedly elevated in the PD-1-deficient cohort. Data are presented as mean ± SEM. Statistical significance was determined by unpaired two-tailed Student’s t-test, except that Mann-Whitney test was used for panel (A), and Welch’s t-test was used for Panel (B) and Granzyme A expression within the CD8^+^ T cells of lung tissue in panel (D). *p < 0.05, **p < 0.01, ***p < 0.001. n = 4-6 mice per group.

To evaluate functional capacity, PBMCs, isolated lung cells, and purified cardiac CD45⁺ leukocytes from control PD-1^f/f^ wt and PD1KO^CD8^ mice were stimulated *ex vivo* with PMA and ionomycin, followed by intracellular staining for Granzyme A, IFN-γ, and the proliferation marker Ki-67. Interestingly, the percentages of LV IFN-γ⁺ and Granzyme A⁺ cells within the cardiac CD8⁺ T cell population were significantly elevated compared to PD-1^f/f^ wt controls (**Figure 2D, 2E, and Supplementary Figures S13A-S13B**). Systemically, this enhanced effector profile was consistently observed, as the proportions of IFN-γ⁺ and Granzyme A⁺CD8⁺ T cells were significantly increased in both the PBMCs and lungs of PD1KO^CD8^ mice (**Figures 2D, 2E, and Supplementary Figures S11B and S15A-S15B**). The proliferation marker, Ki-67, did not change in CD8⁺ T cells from PBMC or heart (**Supplementary Figures S11C, and S13A-S13B**), but increased significantly in CD8⁺ T cells from the lungs of PD1KO^CD8^ mice (**Supplementary Figures S15A and S15B**). In contrast, intracellular IFN-γ and Ki-67 in CD4⁺ T cells from PBMC, lung, and heart were largely unaffected (**Supplementary Figures S11E-S11F, S13C-S13D, and S15C-S15D**). Taken together, these findings indicate that cell-intrinsic PD-1 signaling plays a critical role in restraining CD8⁺ T cell baseline activation, proliferation, and production of cytotoxic molecules, while the CD4⁺ T cell compartment remains functionally unaffected.

### 3.4 Cell-intrinsic PD-1 signaling in CD8⁺ T Cells Is Essential to Mitigate Pressure Overload-Induced Mortality, LV Hypertrophy, Dysfunction, and Inflammation in Mice of Both Sexes

To further evaluate the effect of PD-1 deletion on intrinsic CD8⁺ T cell tolerance under stressed conditions, PD1^f/f^ wt and PD1KO^CD8^ mice were subjected to TAC-induced pressure overload. The data show a profoundly exacerbated HF phenotype. Echocardiographic analysis revealed that both male and female CD8-specific PD-1 KO mice developed significantly worse LV systolic dysfunction, as evidenced by a marked reduction in LVEF, accompanied by more severe ventricular dilation (increased LVESD and LVEDD) compared to their sex-matched PD1^f/f^ wt controls (**Figure 3A, 3D, and Supplementary Figure S16A and S16D**). Consistent with this accelerated functional decline, Kaplan-Meier survival analysis demonstrated increased mortality in both male and female KO mice subjected to TAC, with statistical significance in female mice (**Figure 3B, 3E**). In addition, TAC-induced LV hypertrophy (LV/TL), left atrial dilation (LA/TL), pulmonary congestion (lung/TL), and right ventricular enlargement (RV/TL) were also significantly increased in the PD1KO^CD8^ mice after TAC when compared to the sex-matched PD1^f/f^ wt littermates (**Figure 3C, 3F**). Taken together, these data provide definitive evidence that cell-intrinsic PD-1 signaling in CD8⁺ T cells is a critical, indispensable regulatory checkpoint that limits adverse cardiac remodeling, preserves ventricular function, and promotes survival during chronic systolic overload.

**Figure 3.**
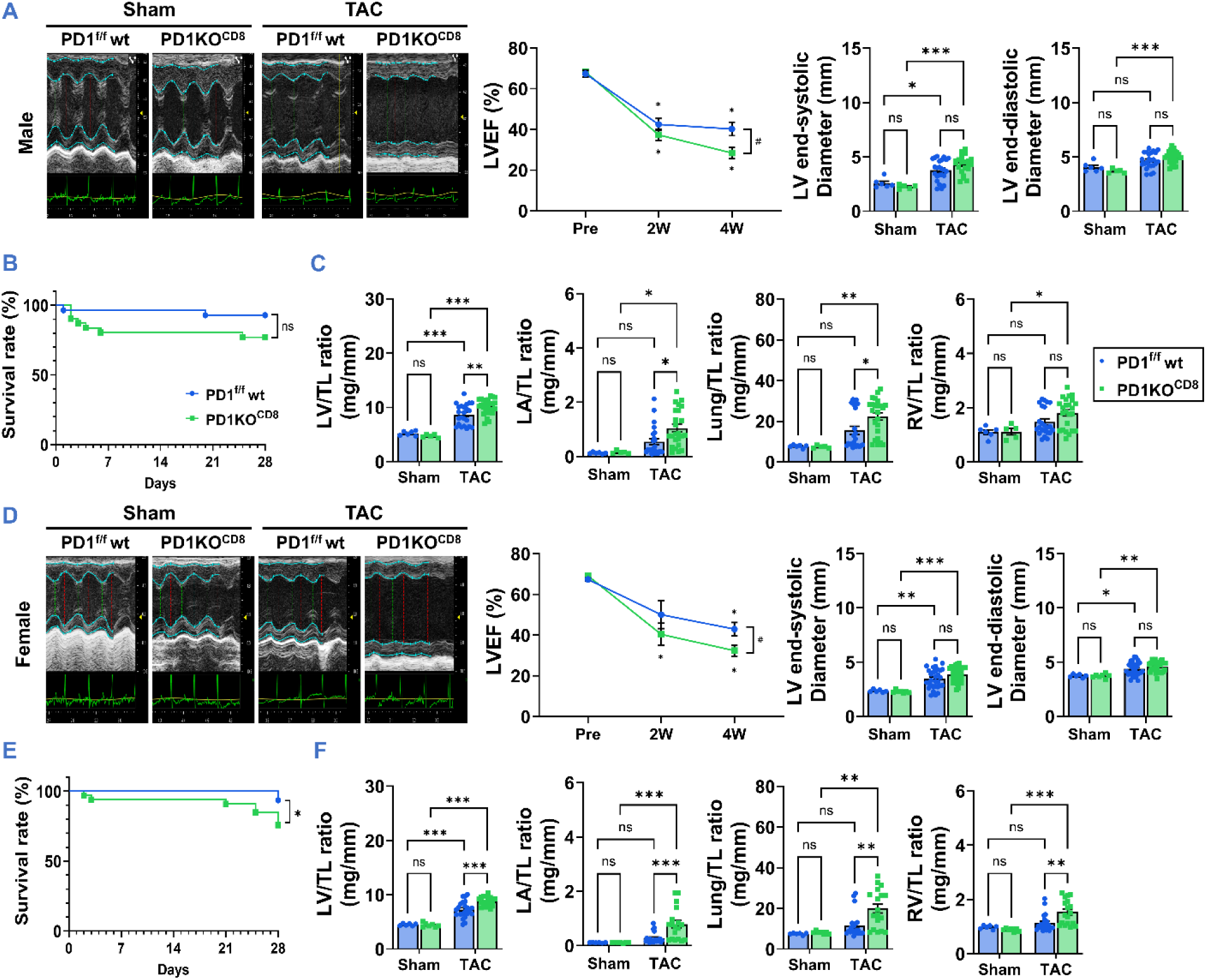
Cell-intrinsic PD-1 ablation in CD8⁺ T cells severely exacerbates cardiac dysfunction, adverse remodeling, and mortality following pressure overload in both male and female mice. **(A)** Representative M-mode echocardiography images (left) and quantitative echocardiographic analysis (right) of LVEF%, LV Internal Dimension at end-systole (LVESD), and end-diastole (LVEDD) in male wild-type (PD1^f/f^ wt, blue) and CD8⁺ T cell-specific PD-1 knockout (PD1KO^CD8^, green) mice following sham or transverse aortic constriction (TAC) surgery. **(B)** Kaplan-Meier survival curves for male PD1^f/f^ wt and PD1KO^CD8^ mice subjected to TAC. **(C)** Gravimetric analysis of male mice at the study endpoint, displaying Left Ventricle-to-Tibia Length (LV/TL), Right Ventricle-to-Tibia Length (RV/TL), Lung-to-Tibia Length (Lung/TL), and Left Atria-to-Tibia Length (LA/TL) ratios. **(D)** Representative M-mode echocardiograms and corresponding quantitative analysis of LVEF%, LVESD, and LVEDD in female PD1^f/f^ wt and PD1KO^CD8^ mice subjected to sham or TAC surgery. **(E)** Kaplan-Meier survival curves for female PD1^f/f^ wt and PD1KO^CD8^ mice following TAC. **(F)** Gravimetric analysis of female mice at the study endpoint (LV/TL, RV/TL, Lung/TL, and Atria/TL ratios). Data are presented as mean ± SEM. Statistical significance was determined by two-way ANOVA with Tukey’s multiple comparisons test. ns = not significant, *p < 0.05, **p < 0.01, ***p < 0.001. n = 5-31 mice per group.

In addition, following TAC, targeted deletion of PD-1 in CD8^+^ T cells resulted in a markedly exacerbated inflammatory and hypertrophic response in the myocardium. Immunostaining of LV sections with CD45 or CD8 antibodies reveals a profound increase in the number of infiltrating CD45^+^ leukocytes and CD8^+^ cytotoxic T cells in the LV of both male and female PD1KO^CD8^ mice, when compared to the sex-matched PD1^f/f^ wt mice (**Figure 4A and 4B**). WGA staining further revealed significantly greater TAC-induced cardiomyocyte hypertrophy in PD1KO^CD8^ mice than in their PD1^f/f^ wt counterparts (**Figure 4A and 4B**), which is consistent with the tissue weight data. Flow cytometry analysis of cardiac tissue showed that the relative proportions of several major immune cell subsets were generally comparable within the population of cardiac CD45⁺ cells between the PD1KO^CD8^ mice and PD1^f/f^ wt littermates in both sexes (**Supplementary Figure S16B and S16E**). However, female PD1KO^CD8^ mice exhibited a significant but mild increase in B cell subsets and a trend toward increased neutrophils, along with reduced NK, following TAC (**Supplementary Figure S16E**). Further analysis of CD8⁺ and CD4⁺ T cell subsets within the cardiac CD3⁺ T cell population revealed a modest but significant increase in the CD8⁺ fraction and a corresponding decrease in the CD4⁺ fraction in both male and female PD1KO^CD8^ mice relative to their PD1^f/f^ wt littermates after TAC (**Figure 4C, 4D, and Supplementary Figure S16C and S16F**). Moreover, cardiac CD8⁺ T cell activation and their intracellular production of IFNγ were significantly higher in PD1KO^CD8^ mice as compared to their PD1^f/f^ wt littermates in both male and female mice, whereas cardiac CD4⁺ T cell activation and their intracellular production of IFNγ were unaffected (**Figure 4C and 4D and Supplementary Figures S17-S19**). Collectively, these findings demonstrate that CD8⁺ T cell-specific PD-1 deficiency is sufficient to exacerbate TAC-induced mortality (female), cardiac dysfunction, hypertrophic remodeling, fibrosis, and immune cell infiltration in both male and female mice, with some sex-specific differences in the composition of the infiltrated cardiac immune cells.

**Figure 4.**
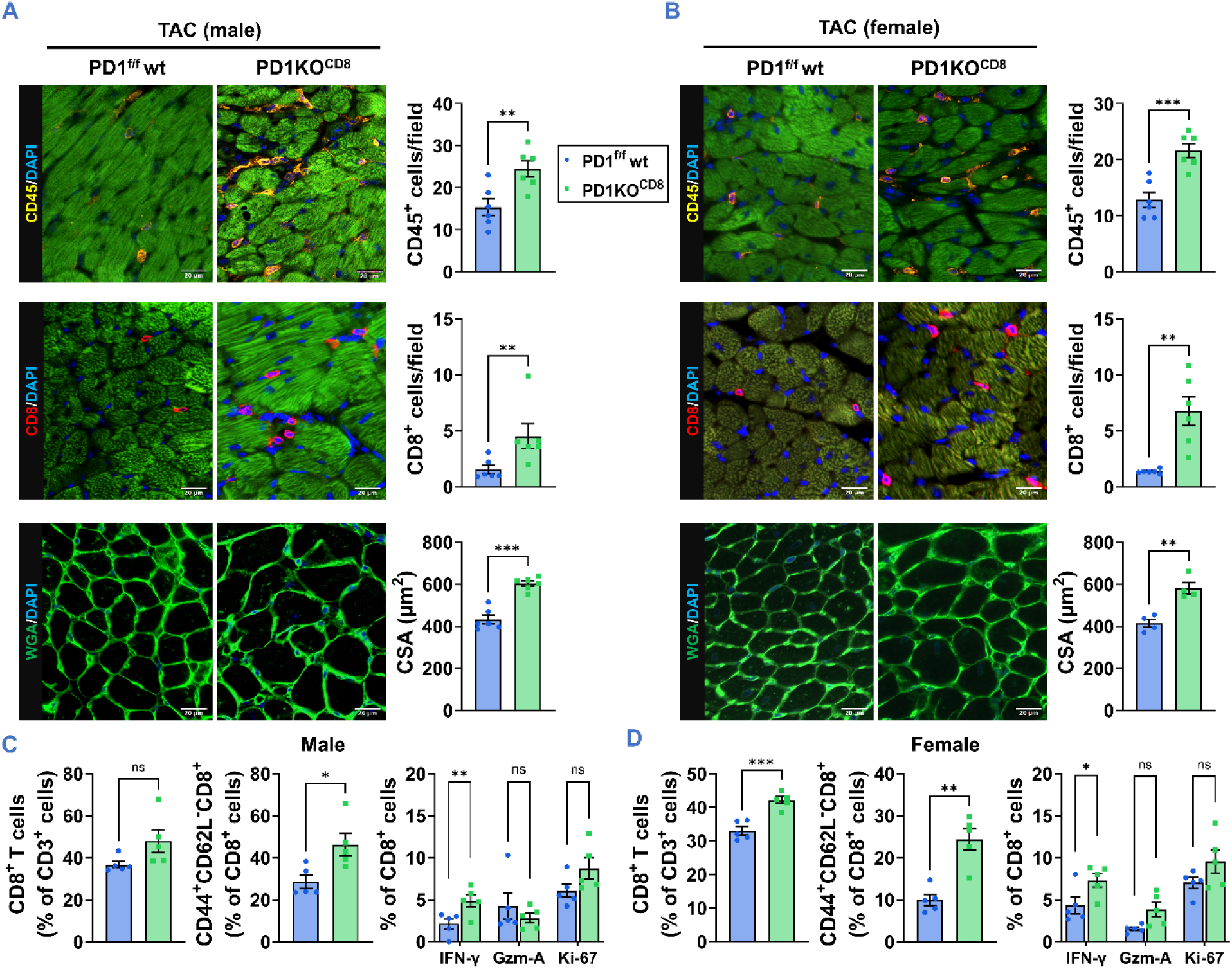
CD8-specific PD-1 KO exacerbates cardiac inflammation and hypertrophy following TAC. (A-B) Representative immunofluorescence images and corresponding quantification of LV sections from male **(A)** and female **(B)** PD1 wt (blue) and PD1KO (green) mice post-TAC. Tissue sections were stained for CD45 (top row) to identify total leukocytes, CD8 (middle row) to identify cytotoxic T cells, and WGA (bottom row) to measure cardiomyocyte cross-sectional area. Nuclei were counterstained with DAPI (blue). Scale bar = 20 μm. **(C-D)** Flow cytometric analysis quantifying infiltrating immune cell populations in LV tissues from male **(C)** and female **(D)** mice following TAC. Data are presented as mean ± SEM. Statistical significance was determined by unpaired two-tailed Student’s t-test, except that Mann-Whitney test was used for infiltrating CD8 in panel **(A)** and % of granzyme A CD8 T cells in panel **(C)**, and that Welch’s t-test was used for infiltrating CD8 in panel **(A)**, % of CD8 T cells in panel **(C)**, and % of granzyme A CD8 T cells in panel **(D)**. ns = not significant, *p < 0.05, **p < 0.01, ***p < 0.001. n = 5-6 mice per group.

### 3.5 CD8-Specific PD-1 Deletion Promotes the Cardiac Transcriptome Associated with Inflammation and Fibrosis Following TAC

To determine the mechanisms by which CD8-specific PD-1 deletion alters TAC-induced LV remodeling, we performed bulk RNA sequencing on LV tissues (**Figure 5**). Principal component analysis (PCA) revealed distinct clustering of LV gene profiles between the PD1KO^CD8^ TAC and PD1^f/f^ wt TAC groups, whereas the sham groups clustered closely together (**Figure 5A**). Evaluation of differentially expressed genes (DEGs) demonstrated that transcriptional changes associated with PD-1 deficiency are predominantly stress-dependent, with 2,384 DEGs detected post-TAC compared to only two in sham conditions (**Supplementary Figure S20A**). Heatmap clustering and volcano plots confirmed that CD8-specific PD-1 deficiency dramatically amplifies these TAC-induced transcriptomic alterations (**Figure 5B and Supplementary Figure S20B**). Importantly, the specific downregulated genes within this profile include *Scn4b*, *Rpl3l*, *Mylk4*, and *Pdp2*, highlighting a transcriptomic signature indicative of suppressed LV fibrosis in the PD1KO^CD8^ cohort, which was accompanied by a significant upregulation of canonical stress and remodeling markers like *Nppb*, *Nppa*, *Myh7*, and *Tgfb2* (**Figure 5B**). Further pathway analysis identified several major molecular networks driving these changes (**Supplemental Figure S20C**). While TAC induced pathways in PD1^f/f^ wt mice as expected, PD1KO^CD8^ TAC mice exhibited distinct Kyoto Encyclopedia of Genes and Genomes (KEGG) enrichment in propanoate metabolism, focal adhesion, and PI3K-Akt signaling (**Supplemental Figure S20C**).

**Figure 5.**
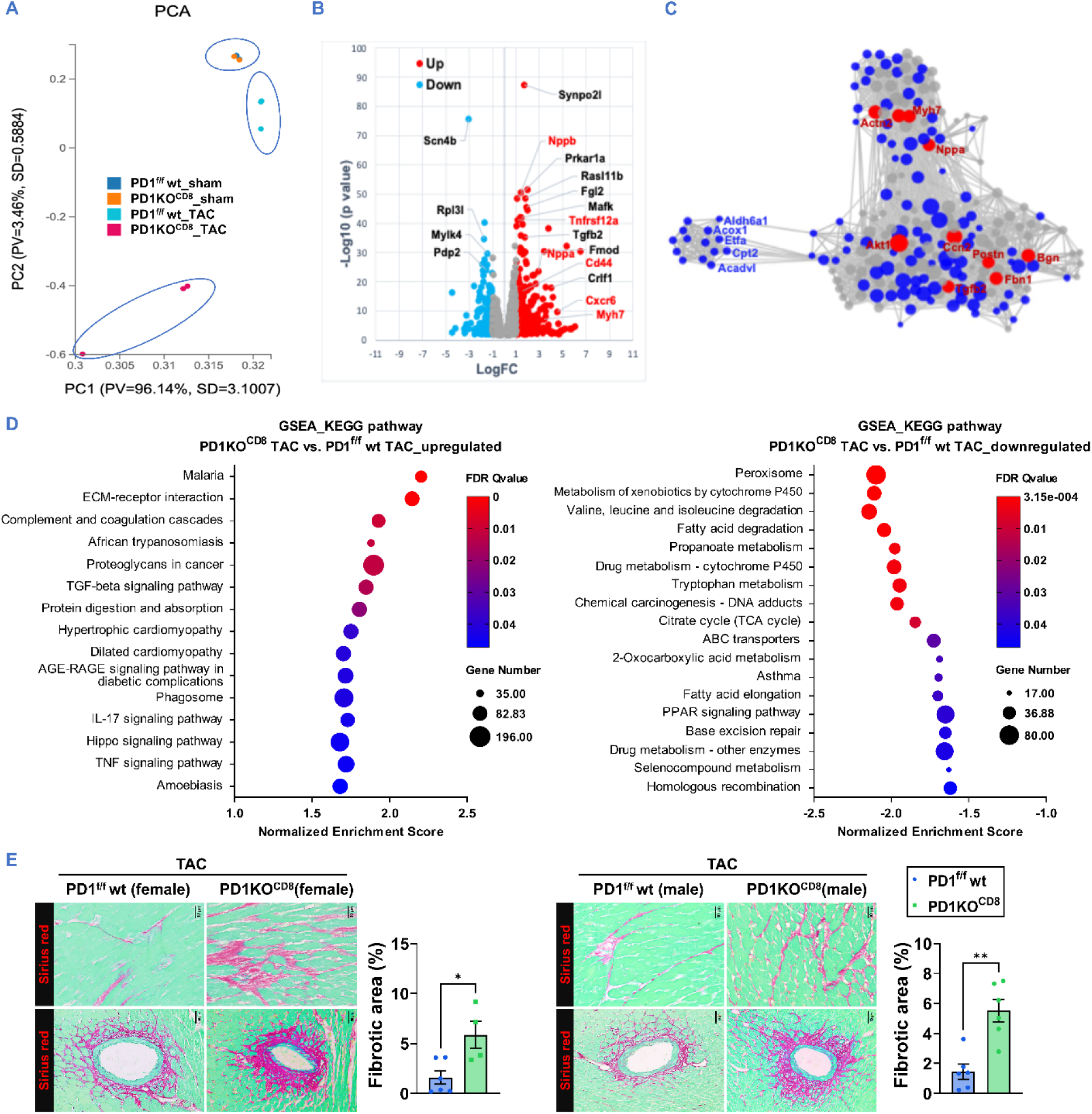
CD8-specific PD-1 KO alters the cardiac transcriptome and suppresses LV fibrosis following TAC. **(A)** PCA plot of bulk RNA-seq data from male LV tissue, clustering the experimental sham and TAC groups. **(B)** Volcano plot of differentially expressed genes post-TAC. Red indicates significant upregulation (e.g., Nppb, Myh7, Tgfb2), while blue highlights downregulated genes (e.g., Scn4b, Rpl3l, Mylk4, Pdp2) as a transcriptomic signature indicative of suppressed LV fibrosis in the KO cohort. **(C)** Key Driver Analysis (KDA) network revealing three major regulatory clusters: an upregulated fibrosis and ECM remodeling cluster (e.g., Postn, Fbn1, Bgn, Akt1), a cardiac stress and hypertrophy cluster (e.g., Myh7, Nppa, Actn2), and a downregulated mitochondrial and metabolic cluster (e.g., Cpt2, Acadvl, Aldh6a1). Red nodes designate key driver genes; blue nodes represent selected significantly changed genes. **(D)** GSEA pathway enrichment dot plots of the up-and downregulated pathways, highlighting that ECM or fibrosis-related pathways are drastically upregulated (left), while mitochondrial or metabolic-related pathways are downregulated (right). **(E)** Representative Sirius red-stained sections and quantification of LV interstitial (Scale bar = 30 μm) and perivascular fibrosis (Scale bar = 50 μm). Histological analysis reconfirms that LV fibrosis is significantly reduced in CD8-specific PD-1 KO mice of both sexes following TAC compared to the PD1 wt controls. Blue bars represent PD1 wt+TAC mice; green bars represent PD1KO +TAC mice. Data are presented as mean ± SEM. Statistical significance was determined by unpaired two-tailed Student’s t-test. *p < 0.05, **p < 0.01. n = 6 mice per group.

Key Driver Analysis (KDA) further delineated this transcriptomic reprogramming by revealing three major regulatory clusters within the network of significantly changed genes (**Figure 5C**). The first cluster is strongly associated with fibrosis and extracellular matrix (ECM) remodeling, featuring key upregulated driver genes such as Postn, Fbn1, Bgn, Tgfb2, Ccn2, and Akt1. The second major cluster is characterized by classical cardiac stress and hypertrophy markers, highlighted by upregulated nodes including Myh7, Nppa, and Actn2. The third distinct cluster predominantly comprises mitochondrial and metabolic genes, physically separated in the network and highlighted by central downregulated nodes including Cpt2, Acadvl, Aldh6a1, Acox1, and Etfa. Consistent with this network architecture, Gene Set Enrichment Analysis (GSEA) revealed a profound functional shift, confirming that metabolism-related pathways are significantly downregulated, while ECM-and fibrosis-related pathways are markedly upregulated (**Figure 5D**).

GSEA using ReactomeGSA further characterized this functional divergence. Compared to the PD1^f/f^ wt TAC controls, PD1KO^CD8^ mice showed a robustly upregulated pro-inflammatory and fibrotic signature, including malaria, ECM-receptor interaction, complement and coagulation cascades, African trypanosomiasis, proteoglycans in cancer, and TGF-β signaling, and protein digestion and absorption, as well as hypertrophic cardiomyopathy, dilated cardiomyopathy, IL-17 and TNF signaling pathways, indicating that under TAC stress, loss of PD-1 in CD8⁺ T cells is associated with a strong inflammatory, immune–matrix remodeling, and profibrotic transcriptional program (**Figure 5D**, left panel). These findings were consistent with heightened LV immune cell infiltration, as shown in **Figure 4**. Conversely, there was a profound global suppression of metabolic and mitochondrial programs, including peroxisome, metabolism of xenobiotics by cytochrome P450, valine/leucine/isoleucine degradation, fatty acid degradation, propanoate metabolism, and TAC cycle, suggesting a global suppression of metabolic and mitochondrial programs after TAC, with a particularly strong impact on lipid and amino-acid metabolism and detoxification capacity (**Figure 5D**, right panel). Together, GSEA using ReactomeGSA and KEGG pathways showed that PD1KO^CD8^ KO significantly upregulated LV pathways associated with inflammation, immune–matrix remodeling, and a profibrotic transcriptional program.

Finally, functional and structural validation of these molecular signatures was performed via histological assessment. Sirius red staining in both male and female LV tissues reconfirms that LV interstitial and perivascular fibrosis are significantly enhanced in CD8-specific PD-1 KO mice of both sexes following TAC (**Figure 5E**). Together, these data demonstrate that the loss of PD-1 on CD8⁺ T cells fundamentally alters both metabolic and fibrotic transcriptional networks, ultimately converging to attenuate pathological collagen deposition.

### 3.6 CXCR3 Blockade Attenuates Exacerbated Cardiac Dysfunction and Remodeling in CD8-Specific PD-1 KO Mice Following TAC

T cell activation induces the expression of tissue-homing receptors, such as CXCR3, which facilitate targeted trafficking to peripheral tissues. To determine whether the exacerbated cardiac phenotype in CD8-specific PD-1 KO mice is mediated by the CXCR3 signaling axis, we administered a CXCR3-neutralizing antibody or an IgG isotype control for 7 days following TAC (**Figure 6A**). Echocardiographic analysis on day 7 revealed that isotype treated PD1KO^CD8^ mice exhibited severe cardiac dysfunction, characterized by significantly reduced LVEF% and LVFS%, along with pronounced LV dilation, when compared to the IgG-treated PD1^f/f^ wt mice (**Figure 6B and 6C**). Whereas, CXCR3 neutralizing antibody significantly preserved systolic function and attenuated maladaptive structural remodeling (**Figure 6B, 6C**).

**Figure 6.**
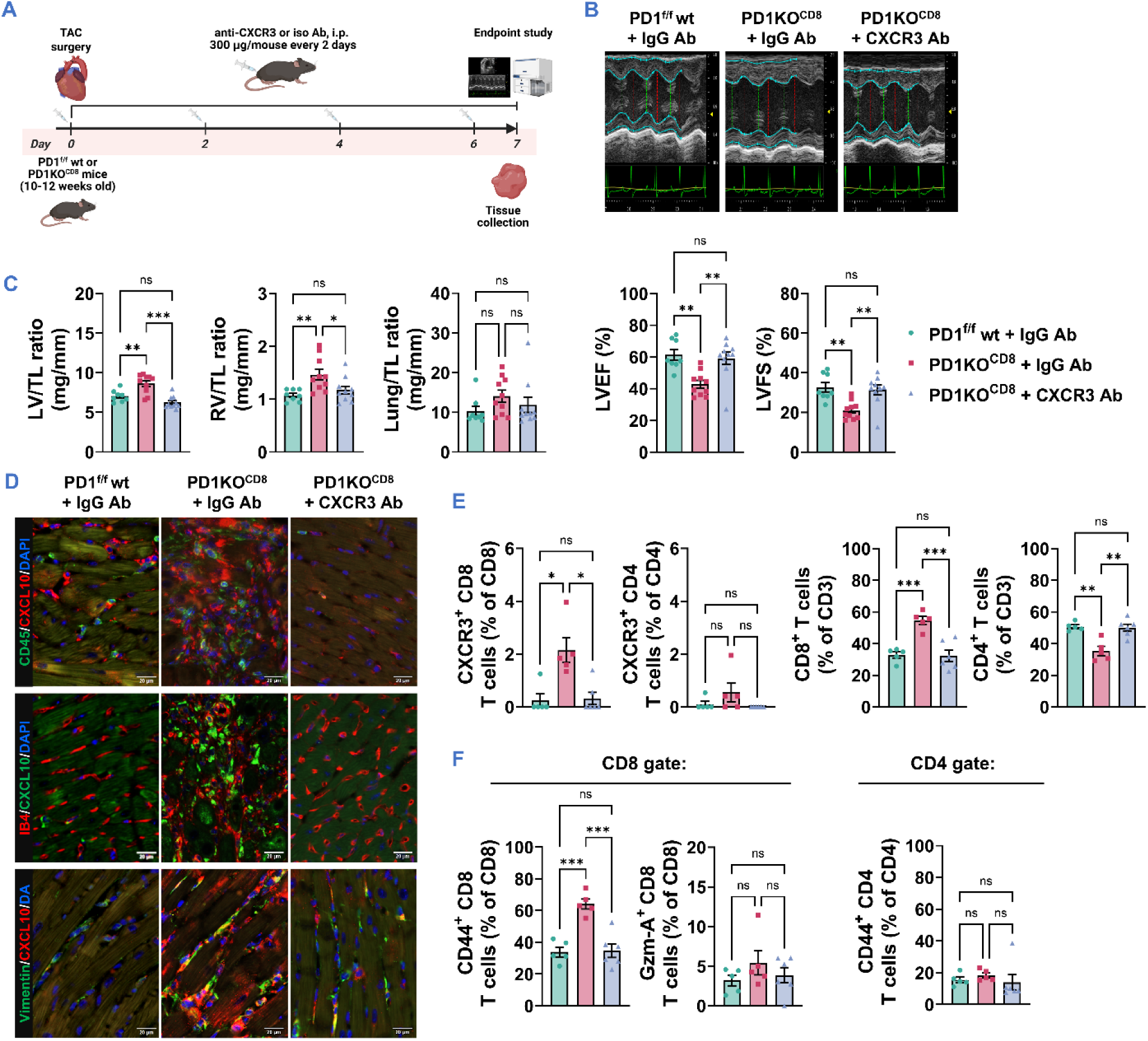
CXCR3 blockade attenuates cardiac dysfunction and pathological remodeling in CD8-specific PD-1 KO mice following TAC. **(A)** Experimental schematic. PD1 wt and PD1KO mice were subjected to TAC surgery at day 0 and received either an anti-CXCR3 neutralizing antibody (anti-CXCR3) or an isotype control (iso Ab) every 2 days until day 7. **(B)** Representative M-mode echocardiographic images of the LV at day 7 post-TAC (top) and quantitative analysis of echocardiographic parameters: LVEF%, and LVFS% (bottom) in PD1^f/f^ wt+TAC+iso Ab (green), PD1KO^CD8^+TAC+iso Ab (pink), and PD1KO +TAC+anti-CXCR3 Ab (light blue). **(C)** Assessment of cardiac hypertrophy via LV/TL ratio, pulmonary congestion by Lung/TL ratio, and right RV/TL ratio. **(D)** Representative confocal images of LV sections co-stained for CXCL10 and specific cell markers: CD45 (leukocytes), IB4 (endothelial cells), or Vimentin (fibroblasts). Nuclei were counterstained with DAPI (blue). Scale bar = 20 μm. **(E-F)** Flow cytometric analysis quantifying infiltrating immune cell subsets of CD8 and CD4 T cells in LV tissues. Data are presented as mean ± SEM. Statistical significance was determined by one-way ANOVA with Tukey’s multiple comparisons test, except that Kruskal-Wallis test with Dunn’s multiple comparisons test was used for LVEF% **(B)**, Lung/TL ratio **(C)**, % of CXCR3 CD8 T cells **(E)**, % of CXCR3 CD4 T cells **(E)**, and % of CD44 CD4 T cells **(F)**. ns = not significant, *p < 0.05, **p < 0.01, ***p < 0.001. n = 5-6 mice per group.

Given the critical role of chemokines in driving pathological remodeling, we investigated the local expression of CXCL10, a primary ligand of CXCR3. Immunofluorescence co-staining demonstrated that CXCL10 is prominently expressed across multiple cell types in the LV myocardium, including CD45⁺ leukocytes, IB4⁺ endothelial cells, Vimentin⁺ fibroblasts, and cardiomyocytes (**Figure 6D**). Further characterization of the local immune profile via flow cytometry confirmed that the robust expansion, activation, and recruitment of infiltrating CD8⁺ T cells observed in PD1KO^CD8^ mice were significantly suppressed by systemic CXCR3 neutralization (**Figure 6E, 6F**).

Taken together, these findings demonstrate that genetic ablation of PD-1 in CD8⁺ T cells exacerbates TAC-induced HF, at least in part through a CXCR3-dependent mechanism. Disrupting this chemokine signaling axis effectively mitigates maladaptive immune infiltration and cellular responses, highlighting CXCR3 blockade as a potent intervention to rescue the severe structural and functional deterioration observed in this model.

### 3.7 CD8-Specific PD-1 Deletion Exacerbates Pulmonary Inflammation, Remodeling, and T Cell Effector Function Following TAC

Our previous studies demonstrated that LV failure causes secondary pulmonary remodeling and profound inflammation. To determine the effect of CD8-specific PD-1 deletion on these processes, we assessed lung tissues following TAC. Immunofluorescence staining revealed that TAC significantly increased the infiltration of CD45^+^ leukocytes and CD8^+^ T cells, an effect that was profoundly exacerbated in PD1KO^CD8^ mice of both sexes (**Supplemental Figures S21A-S21B and S24A-S24B**). Similarly, PD-1 deletion in CD8⁺ T cells significantly exacerbated TAC-induced pulmonary fibrosis (**Supplemental Figures S21C and S24C**). Flow cytometric profiling of pulmonary CD45⁺ leukocytes confirmed broad shifts in the immune composition of PD1KO^CD8^ mice post-TAC, with notable alterations in NK cells, dendritic cells, and neutrophils or macrophages, depending on sex (**Supplemental Figures S21D and S24D**). Within the CD3⁺ compartment, PD1KO^CD8^ mice consistently displayed a significantly higher percentage of CD8⁺ T cells and a reciprocal reduction in CD4⁺ T cells under TAC conditions (**Supplemental Figures S21E-S21F and S24E-S24F**).

Previously, flow cytometry analysis of pulmonary T cell activation states at baseline revealed a striking phenotype: PD1KOCD8 mice exhibited a dramatic shift from naïve (CD44⁻CD62L⁺, T_naïve_) to effector memory (CD44⁺CD62L⁻, T_EM_) CD8⁺ T cell phenotype (**Figure 2B and Supplemental Figure S14**). This naïve-to-TEM transition was further exacerbated following TAC in both sexes (**Supplementary Figure S22A and S25A**). Central memory CD8⁺ T cells were modestly altered. Pulmonary CD4⁺ T cells showed mild and not significant changes in both male and female PD1KO^CD8^ mice post-TAC (**Supplementary Figure S22B and S25B**).

Consistent with this enhanced memory differentiation, *ex vivo* stimulation of isolated pulmonary leukocytes demonstrated a functional hyperactivation following TAC that was further amplified in both male and female PD1KO^CD8^ mice, exhibiting significantly greater proportions of IFN-γ⁺, Granzyme A⁺, and proliferating (Ki-67⁺) CD8⁺ T cells in PD1KO^CD8^ mice (**Supplementary Figure S23A and S26A**). In contrast, functional alterations in pulmonary CD4⁺ T cells were minimal or entirely absent in both sexes (**Supplementary Figure S23B and S26B**). Collectively, these findings demonstrate that cell-intrinsic PD-1 deficiency drives progressive pulmonary CD8⁺ T cell accumulation, effector memory differentiation, and proinflammatory cytokine production. This targeted T cell dysregulation is sufficient to induce more severe pulmonary remodeling and inflammation during TAC-induced HF progression.

Given that sustained pulmonary remodeling can subsequently drive RV hypertrophy, we next examined RV histology and immune cell infiltration in male and female PD1^f/f^ wt and PD1KO^CD8^ mice. WGA staining showed that TAC induced a significant increase in RV cardiomyocyte size in both male and female PD1KO^CD8^ mice when compared to the PD1^f/f^ wt mice (**Supplementary Figure S27A and S27E**). Sirius red staining revealed no significant differences in RV fibrotic area among the groups (**Supplementary Figure S27B and S27F**).

Immunofluorescence staining showed no difference in CD45⁺ leukocyte infiltration in the RV between the PD1KO^CD8^ and PD1^f/f^ wt mice after TAC (**Supplementary Figure S27C and S27G**), but a significant increase in the infiltration of CD8⁺ T cells was observed in the female PD1KO^CD8^ mice’s RV tissue (**Supplementary Figure S27D and S27H**), when compared to the PD1^f/f^ wt mice after TAC. These findings demonstrate that CD8⁺ T cell-specific PD-1 deficiency exacerbates RV hypertrophy following TAC in both sexes, independent of significant fibrotic remodeling, and is accompanied by modestly increased immune cell infiltration, specifically in females.

### 3.8 Regulatory T Cell-Specific PD-1 Deficiency Does Not Alter Baseline or TAC-Induced Cardiac Remodeling

Regulatory T cells (Tregs) constitutively express high levels of PD-1 and play a well-documented, critical role in restraining excessive cardiopulmonary inflammation. Given this prominent immunomodulatory function, we tested the hypothesis that disrupting PD-1 signaling specifically within the Treg compartment might alter their suppressive capacity and subsequently influence cardiac remodeling. To investigate this, we generated a male Treg-specific PD-1 KO model by crossing Foxp3^Cre^ mice with the PD1^f/f^ strain (**Supplementary Figure S28**).

Surprisingly, comprehensive echocardiographic and structural evaluation revealed that Treg-specific PD-1 deletion had no detectable impact on the cardiac phenotype. Under baseline control conditions, LV, lung, and RV weight to TL ratios remained completely unaffected (**Supplementary Figure S29A**). Furthermore, following TAC-induced pressure overload, these mice exhibited a hypertrophic and functional decline that was virtually indistinguishable from their wild-type counterparts, with no significant exacerbation of LV dilation or functional impairment between the PD1KO^FoxP3^ and PD1^f/f^ wt mice (**Supplemental Figure S29B-S29C**).

Also, no difference in the infiltration of CD45^+^ leukocytes was detected between the PD1KO^FoxP3^ and PD1^f/f^ wt mice after TAC (**Supplemental Figure S29D)**. These unexpected findings highlight that the exacerbated HF phenotype observed in our study is uniquely driven by the CD8⁺ effector T cell compartment, rather than by a secondary loss of Treg-mediated immunosuppression.

## 4. Discussion

The present study identifies CD8⁺ T cells as critical mediators of cardiac and pulmonary pathology in a mouse model of pressure overload-induced HF, and establishes that cell-intrinsic loss of PD-1 signaling within CD8⁺ T cells is sufficient to recapitulate and exacerbate this phenotype independent of PD-1 deficiency in other cell types. PD-1 in CD8 cells plays a more critical role in restraining CD8 T cell cytotoxic molecule production and TAC-induced cardiac inflammation and HF, whereas PD-1 deletion in FoxP3^+^ T regulatory cells has no detectable effect. Using both antibody-mediated CD8⁺ T cell depletion in global PD1KO mice and a CD8⁺ T cell-specific PD-1 KO model (PD1KO^CD8^), we demonstrate that CD8⁺ T cells drive systolic dysfunction, cardiac hypertrophy, interstitial fibrosis, and pulmonary inflammation following TAC, and that these effects are substantially amplified when PD-1, a central inhibitory checkpoint, is selectively deleted from this cell population.

One key finding of this work is that CD8⁺ T cell-specific PD-1 deletion alone was sufficient to reduce survival, impair LV systolic function, and exacerbate cardiac and pulmonary remodeling following TAC. This observation extends our prior work using CD8⁺ T cell Egln1-deficient mice ^6^, and provides direct genetic evidence that PD-1 signaling acts intrinsically within CD8⁺ T cells to restrain pathological cardiac inflammation during chronic pressure overload.

These findings are also consistent with clinical and experimental evidence implicating disruption of PD-1 signaling in ICI-associated myocarditis and may help explain the greater susceptibility to ICI-related cardiotoxicity reported in patients with preexisting cardiovascular disease, in whom underlying cardiac inflammation or preexisting CD8⁺ T-cell activation may amplify the consequences of PD-1 blockade ^11–15^. Our finding is also consistent with results published in *Circulation* by an independent research laboratory ^16^. Both studies reach a similar conclusion using different mouse strains, highlighting the robustness of the data, addressing the critical priority of reproducibility in medical research, and providing additional support for a protective role of CD8⁺ T cell-intrinsic PD-1 signaling in limiting cardiac injury. Importantly, our study extends these observations by demonstrating baseline CD8⁺ T-cell hyperactivation in the blood, heart, and lung of both sexes, supporting a model in which pre-existing systemic and tissue-level immune priming sensitizes the heart to subsequent pressure overload. We further investigate the contribution of regulatory T-cell-intrinsic PD-1 signaling by deleting PD1 specifically in Treg cells.

Another important finding of our study is that transcriptomic profiling provided further mechanistic insight into how CD8⁺ T cell-specific PD-1 loss reshapes the cardiac metabolic and inflammatory profiles. PD1KO^CD8^ hearts subjected to TAC showed selective enrichment of IL-17, TNF, TGF-β, and ECM-receptor interaction signaling pathways, alongside pathways directly linked to hypertrophic and dilated cardiomyopathy, while pathways governing oxidative and lipid/amino acid metabolism (peroxisomal function, fatty acid degradation, the TCA cycle) were coordinately suppressed. This is consistent with the report by Patsoukis et al. that PD1 signaling promotes fatty acid β-oxidation and metabolic reprogramming in activated T cells ^17^. The GO biological pathways enrichment analysis revealed that the major enriched biological pathways in PD1KO^CD8^ mice after TAC were related to collagen fibril organization, extracellular matrix organization, cell adhesion, and mitochondrial function. KEGG pathway analysis of DEGs revealed that the major enriched pathways were related to protein digestion and absorption, various metabolic pathways, hypertrophic/dilated cardiomyopathy, and the HIF-1 signaling pathway. The finding that deletion of PD-1 in CD8^+^ T cells alone significantly enhanced the LV gene expression profiles, particularly those associated with extracellular collagen formation, cellular metabolism, and inflammation, align with the histological and functional deterioration observed in these mice. These findings demonstrate that PD-1 in CD8^+^ T cells exerts important and broad roles in promoting systolic overload-induced LV inflammation, fibrosis, and dysfunction. Our results also suggest a significant crosstalk between CD8^+^ T cells and the surrounding environment during TAC-induced LV inflammation and dysfunction.

The chemokine receptor CXCR3 and its ligands, CXCL9/CXCL10/CXCL11, have been implicated in chronic inflammation, immune dysfunction, cancer, and ICI myocarditis ^18–20^. Ngwenyama et al. previously demonstrated that CXCR3⁺ T cells infiltrate the stressed heart and CXCR3 deletion attenuates T cell recruitment, fibrosis, and dysfunction following TAC, with cardiac fibroblasts and myeloid cells serving as a local source of the CXCL9 or CXCL10 driving this recruitment ^21^. Consistent with this study, we found that CXCL10 expression was elevated in the lesion area and predominantly co-localized with Vimentin⁺ fibroblasts, with only minimal overlap in CD45⁺ leukocytes and IB4⁺ endothelial cells. More recent study by Souza-Neto et al. identified CXCL10 specifically as a chemokine mediator of the CXCR3 pathway, with CXCL10 deficiency similarly reducing T cell recruitment to the heart and mitigating pressure overload-induced remodeling ^22^. These studies, however, implicated CD4⁺ rather than CD8⁺ T cells as the principal CXCR3-responsive population. Huang et al. showed that blockade or depletion of CXCR3 reduced CXCR3^high^ CD8^+^ T cells in the heart and reduced the myocarditis phenotype in a pharmacological mouse model of ICI, linking PD-1 loss to CXCR3-driven CD8⁺ T cell cardiotoxicity in both genetic and pharmacological ICI-myocarditis models ^18^. Our finding that CXCR3 blockade similarly reverses CD8⁺ T cell cardiac accumulation and rescues systolic function in CD8⁺ T cell-specific PD1KO mice under pressure overload closely parallels this ICI myocarditis model. We found that blocking CXCR3 signaling decreased the percentage of CD8^+^ T cells in the CD3^+^ T cell population. It also decreased the percentage of CD44^+^ or Ki-67^+^ CD8 T cells, suggesting that the treatment effectively attenuated the activation and proliferation of CD8^+^ T cells. However, these effects were not seen in CD4^+^ T cells. These data indicate that PD-1 deletion affects CXCR3 signaling in CD8^+^ T cells in our model.

Axelrod et al. reported that depletion of CD4^+^ T cells did not rescue survival ^5^, which partially supports the idea that PD-1 on CD4^+^ T cells may not play a major role in the development of TAC-induced HF. However, they did not investigate the role of CD4^+^ T cells in the initiation of myocarditis. A subpopulation of CD4^+^ T cells, known as CD4^+^CD25^+^Foxp3^+^ Tregs, expresses a high level of PD-1 and modulates immune tolerance and suppresses inflammatory responses in various clinical diseases ^23^. The role of PD-1 on the Treg cells in autoimmunity and cancer therapy remains controversial. Some studies suggest that PD-1 blockade increases Treg proliferation but can either enhance or reduce its suppressive function ^24, 25^. Another study suggested that PD-1 blockade or Treg-specific PD-1 deletion reduces the tumor-infiltrating Treg cells ^26^. Our data show that deletion of PD-1 in Foxp3^+^ Treg cells, a subset of CD4^+^ T cells, did not rescue TAC-induced cardiac dysfunction nor LV hypertrophy, suggesting that PD-1 in Treg cells is not a major player during the development of cardiac inflammation and HF.

Finally, our data demonstrate that CD8⁺ T cell-specific PD-1 deficiency promotes pulmonary inflammation at baseline and HF progression following TAC. Notably, even sham-operated PD1KO^CD8^ mice exhibited increased pulmonary CD8⁺ T cell accumulation, effector memory differentiation, and functional activation, likely reflecting the lung’s constant exposure to environmental antigens, which may unmask the effects of PD-1 deletion even in the absence of TAC-induced pressure overload. PD1KO^CD8^ mice following TAC exhibited even greater pulmonary CD8⁺ T cell accumulation, effector memory differentiation, and functional activation, along with exacerbated right ventricular hypertrophy in both sexes. This suggests that CD8⁺ T cell-mediated inflammation may propagate along the cardiopulmonary axis. These findings are consistent with our previous studies demonstrating that inflammation not only contributes to the development of HF but also to its progression ^27–29^, and that pulmonary inflammation can drive lung remodeling and RV hypertrophy independently of LV function in mice with established HF^29^.

This study has a few limitations. First, while a concurrent publication by another laboratory in *Circulation* ^16^ regarding CD8⁺ T cell-specific PD-1 affects the standalone novelty of our work, the simultaneous identification of this pathway by two independent groups underscores its fundamental biological importance. Furthermore, reproducibility is a critical priority in biomedical research, and our study expands upon this shared finding by comprehensively investigating the impact of CD8⁺ T cell-specific PD-1 on cytotoxic molecule production, activation status, and global cardiac genomic profiles under both baseline and TAC conditions. Second, while we demonstrate that deleting PD-1 in CD8⁺ T cells exacerbates TAC-induced cardiac inflammation, cardiomyocyte hypertrophy, and fibrosis, we did not examine the effects of PD-1 deletion in total CD4⁺ T cells, representing another major lymphocyte population. However, we did establish that deleting PD-1 in CD4⁺FoxP3⁺ regulatory T cells, a subset of CD4⁺ T cells, has no detectable effect on TAC-induced cardiac remodeling and failure. Third, our *in vivo* CXCR3 blockade using a neutralizing antibody lacked cell-type specificity. Because CXCR3 is also moderately expressed across multiple immune subsets, including natural killer cells, macrophages, neutrophils, and B cells, we were unable to definitively isolate the specific contribution of CXCR3 signaling strictly within the CD8⁺ T cell compartment.

In conclusion, our data clearly demonstrate that PD-1 in CD8^+^ T cells plays a critical role in restraining autoreactive CD8^+^ T cell function and TAC-induced cardiac inflammation and HF. Deletion of PD-1 in CD8^+^ T cells alone is sufficient to exacerbate systolic overload-induced cardiac inflammation and HF development, while deletion of PD-1 in FoxP3^+^ Tregs has little effect. These results have direct translational relevance for understanding the increased susceptibility of people with preexisting cardiovascular disease to ICI-induced cardiotoxicity. and nominate CXCR3-dependent CD8⁺ T cell trafficking as a candidate therapeutic target for systolic overload-induced cardiac inflammation and dysfunction, as well as in people with cancer exhibiting immune-related adverse events (iRAEs).

## Translational Perspective

Immune checkpoint inhibitors that block programmed cell death protein 1 raise the risk of heart failure in cancer patients, especially those with hypertension or other cardiovascular disease. Using two cell-specific knockout mouse strains, we show that this protein acts within CD8^+^ T cells, but not within regulatory T cells, to restrain cytotoxic T cell activation, cytotoxic molecule production, and cardiopulmonary inflammation and dysfunction. In addition, CD8^+^ T cell depletion and CXCR3 blockade attenuate pressure overload-induced cardiac inflammation and dysfunction. These findings identify cytotoxic T cells as active drivers of hypertensive cardiopulmonary remodeling and dysfunction.

## Acknowledgments

We would like to thank the histology and flow cytometry core facility for their technical support. We appreciate Dr. Kenneth Weir’s help with manuscript preparation.

## Author contributions

XH, XW, LP, ZN, UB, RX, DW, and YC contributed to data collection and analysis. XH and YC contributed to experimental design and manuscript preparation. HZ, J-XC, JC, AS, BL, BB, MH, and YC edited the manuscript. All authors contributed to the article and approved the submitted version.

## Ethics declarations

The authors declare no competing interests.

