## Supplemental materials for "Treg- and CD8-Specific PD-1 Deficiency Reveals That CD8⁺ T Cell-Intrinsic Signaling Uniquely Restrains Pressure Overload Inflammation and Heart Failure"

Short title: Cell-specific PD1 ablation and heart failure

Conflict of Interest: None.

**#Correspondence**

Yingjie Chen, Ph.D.

Professor

Department of Physiology and Biophysics,

University of Mississippi Medical Center,

2500 North State Street,

Jackson, MS, 39216

**Supplementary Figures S1-S35 and Tables S1-S13**

### Methods

#### *In vivo* depletion of $CD8^+$ T cells or blockade of CXCR3

C57BL/6J, global PD1KO, or $PD1KO^{CD8}$ mice were subjected to TAC surgery for 7 days or 4 weeks and treated with either rat anti-CD8α (clone YTS 169.4, BP0117, 250 µg/mouse/dose) or rat IgG2b isotype (clone LTF-2, BP0090), diluted in 150 µL phosphate-buffered saline (PBS) through intraperitoneal injection every two days, starting immediately after surgery. Additionally, different groups of $PD1^{f/f}$ wt and $PD1KO^{CD8}$ mice were subjected to TAC surgery for 7 days and treated with either hamster anti-CXCR3 blocking antibody (clone CXCR3-173, BE0249, 300 µg/mouse/dose) or Armenian hamster polyclonal IgG isotype (BE0091), diluted in 150 µL phosphate-buffered saline (PBS) through intraperitoneal injection every two days, starting immediately after surgery. All blocking antibodies and their respective isotype control antibodies were purchased from BioXcell (Lebanon, NH). The doses of the CD8-blocking antibodies and CXCR3-blocking antibodies were based on previous reports, with minor modifications [1, 2]. At the end of the study, cardiac function was assessed by echocardiography, and the animals were euthanized for tissue collection and flow cytometry analysis.

#### Echocardiography

Transthoracic echocardiograms were performed by using a Vevo 3100 Preclinical Imaging Platform equipped with an MX400 transducer (FUJIFILM Visual Sonics Inc., Canada). The mice were anesthetized by inhalation of 1–2% isoflurane mixed with 100% medical oxygen. M-mode cine loops were recorded and analyzed by Vevo LAB software (FUJIFILM Visual Sonics Inc., Canada) to assess LV ejection fraction (EF), fractional shortening (FS), end-systolic diameter (LVESD), LV end-diastolic diameter (LVEDD), LV end-systolic volume (LVESV), LV end-diastolic volume (LVEDV), LV anterior wall and posterior wall thickness at end-systole or end-diastole, stroke volume, and cardiac output [3-8]

**Tissue collection and flow cytometry analysis**

Mice were anesthetized with a single intraperitoneal injection of ketamine (100 mg/kg) and xylazine (10 mg/kg). The mice were perfused with cold PBS through the right ventricle (RV) to flush the left ventricle (LV) and lung. The LV was digested in RPMI-1640 medium containing 0.12 mg/mL of Liberase and 60U/mL of DNAse I (Millipore Sigma, MO) and the lung tissue was digested in RPMI-1640 medium containing 1mg/mL of collagenase D and 60U/mL of DNAse I, on the gentle MACS™ Octo dissociator (Miltenyi, CA). The cells were isolated and flow cytometry analysis was performed as previously described with minor modifications [9]. Single-cell suspensions were stained with fixable viability dye (FVS440UV, BD Bioscience, CA) in PBS followed by washing in staining buffer and blocking with anti-mouse CD16/32 (clone 2.4G2, Biolegend, CA) antibodies to prevent non-specific binding of antibodies to FcRγ. The cells were then incubated with the fluorescence-conjugated antibodies listed in Supplementary Table S8, accordingly. For cytokine production analysis, single cell suspensions were stimulated with a cell stimulation cocktail (eBioscience, MA) diluted in RPMI medium containing 10% fetal bovine serum (FBS) for two hours in a cell culture incubator at 37°C with 5% $CO_2$. The cells were then collected and stained with FVS440UV, blocked by anti-mouse CD16/32, and labeled with fluorescence-conjugated antibodies listed in Supplementary Table S8. Samples were run on the BD FACSymphony™ A3 Cell Analyzer (BD Biosciences, CA). Data were analyzed by FlowJo_V10 (FlowJo, OR) software. Gating strategies are shown in the **Supplementary Figures S30-S35** as indicated.

**Bulk RNA Sequencing**

LV tissue samples were collected from WT and PD1KO$^{CD8}$ male mice under sham and 4 weeks after TAC and stored at -80°C until RNA isolation. LV tissue samples were used for RNA extraction and sequencing. HiSeq4000 platform was used for RNA sequencing, and 150 bp paired-end reads were generated. Raw reads were quality-checked and filtered, and clean reads were aligned to the mouse reference genome GCF_000001635.27_GRCm39 using HISAT2. Data analysis was performed using the method as previously reported [10, 11].

**Histological analysis and immunofluorescence staining**

Tissues were excised, fixed in 10% formalin, and then embedded in paraffin. 5-micron sections were stained with Sirius red and Fast green Stain Kit (Chondrex, WA) to evaluate LV, RV, and pulmonary fibrosis. Heart sections were also stained with Alexa Fluor 488-conjugated wheat germ agglutinin (Invitrogen, 5μg/ml) to evaluate cardiomyocyte size in LV and RV. For immunofluorescence staining of $CD45^+$ leukocytes, $CD8^+$ T cells, or CXCL10, antigen retrieval was performed in heated citrate buffer (pH 6.0) and the sections were blocked with 1% bovine serum albumin in PBS for one hour at room temperature, followed by incubation with goat anti-CD45 (R&D systems, 1:100) or rabbit anti-CD8$\alpha$ (Cell Signaling Technology, 1: 100) primary antibodies at 4°C overnight. The tissue sections were then washed with PBS and incubated with Alexa Fluor 555-conjugated secondary donkey anti-goat or goat anti-rabbit (Invitrogen, 1:500), respectively. Additionally, the section was co-stained with goat anti-CD45 and rabbit anti-CXCL10 antibodies (Invitrogen, 1: 100). Endothelial cells were stained by Griffonia Bandeiraea Simplicifolia Isolectin B4 (IB4, 1:50, Invitrogen, OR). The primary antibodies for immunofluorescence staining were listed in the **Supplementary Table S13**. All images were captured using an Olympus BX43 fluorescent microscope, then quantified using the inForm Advanced Image Analysis (Akoya Biosciences, MA) or ImageJ software.

**Immunoblot Analysis**

Protein extractions from LV samples were prepared in RIPA lysis buffer with protease/phosphatase inhibitor cocktail (A32961, ThermoFisher Scientific, NY). Lysates were mixed with Laemmli sample buffer (2X, Bio-Rad, CA) containing 10% mercaptoethanol and boiled at 95°C for 10 minutes. The samples were separated by SDS-PAGE gel and then transferred to a PVDF membrane. Membranes were blocked in 5% milk and washed 3 times with 1X TBST (TBS containing 0.1% Tween-20, Millipore Sigma, MA). The membrane was then cut to separate the desired proteins for probing. Membranes were incubated with primary antibodies against Collagen III (22734-1-AP, Proteintech, IL) or ANP (PA5-29559, ThermoFisher Scientific, NY) and then incubated with an anti-rabbit (31460, 1:10000, ThermoFisher Scientific, NY) secondary antibody conjugated with horseradish peroxidase for 1 hour at room temperature. Secondary antibodies were detected using the ECL Western Blotting Substrate (32106, ThermoFisher Scientific, NY). Samples were analyzed by densitometry using Image Lab software 6.0 (Bio-Rad, CA) and normalized to the loading control.

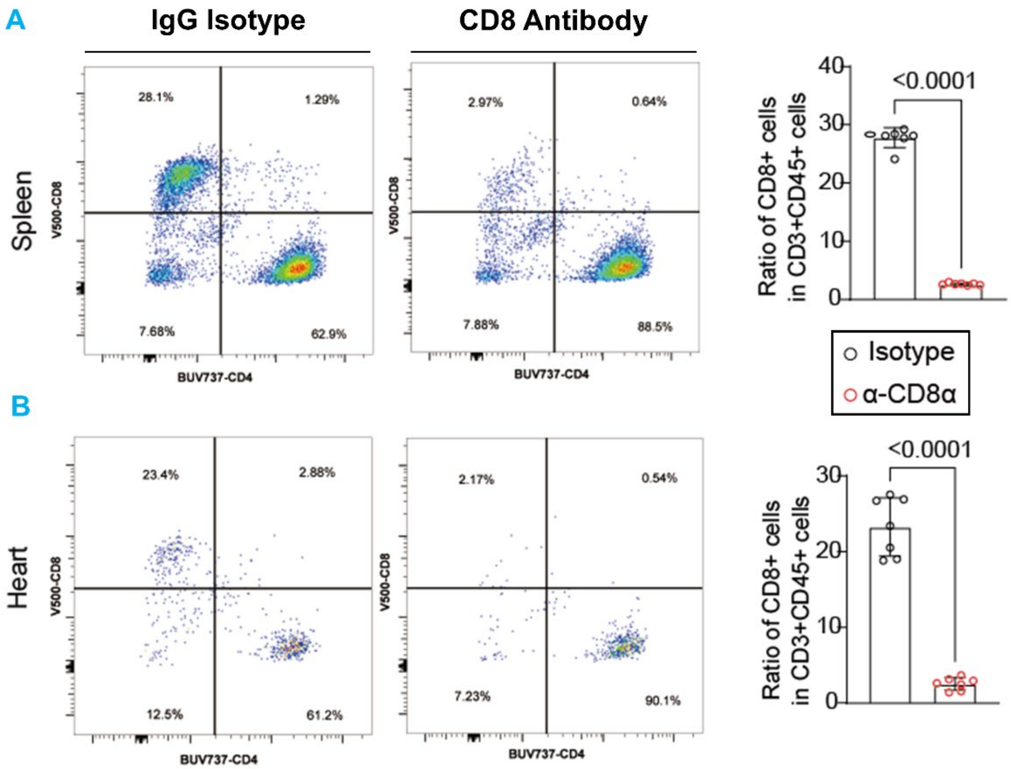


**Supplementary Figure S1. Effect of CD8 antibody on depletion of spleen and cardiac CD8$^+$ T cells.**

**(A)** Representative flow cytometry plots identifying CD4$^+$ and CD8$^+$ T cell populations in the spleen following *in vivo* treatment with either an IgG Isotype control (left, black circles) or a CD8-depleting antibody (right, red circles). The accompanying bar graph (right) quantifies depletion efficacy, showing the relative percentage of CD8$^+$ cells in the total CD3$^+$ T cell population. **(B)** Representative flow cytometry plots and corresponding quantitative analysis demonstrates the successful depletion of CD8$^+$ T cells in cardiac tissue of mice treated with the CD8 antibody compared to the IgG Isotype control. Data are presented as mean ± SEM. Statistical significance was determined by unpaired two-tailed Student's t-test. n = 7 mice per group.

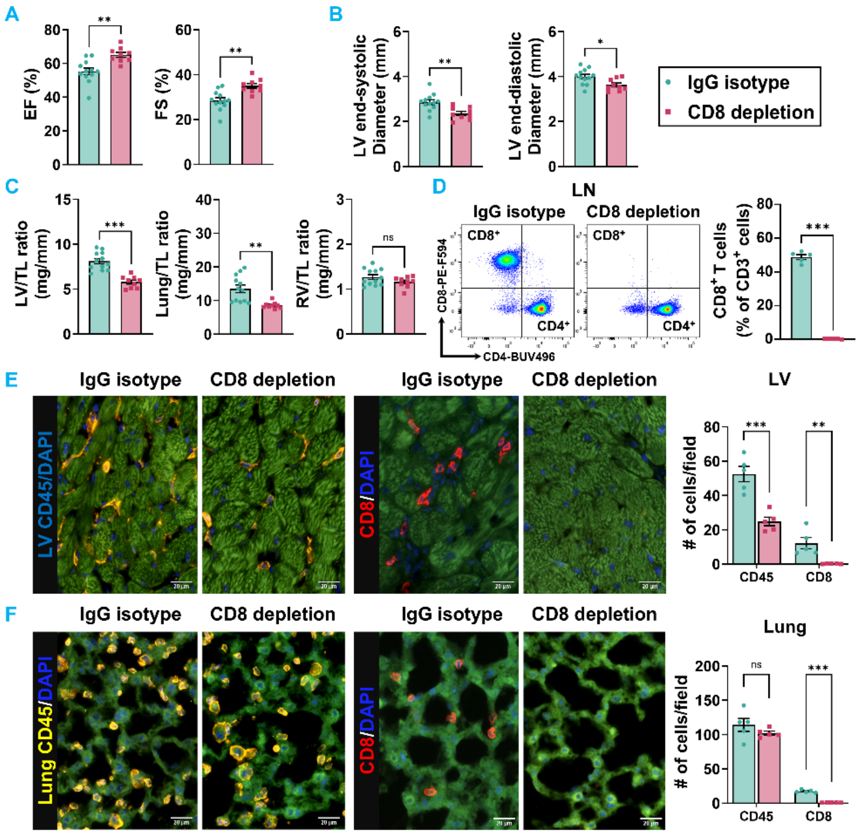


**Supplementary Figure S2. Early CD8$^{+}$ T cell depletion preserves cardiac function and attenuates inflammation at 7 days post-TAC in PD1KO mice.**

**(A)** Quantitative analysis of early systolic function, including left ventricular ejection fraction (LVEF%) and fractional shortening (LVFS%), evaluated at 7 days post-TAC with (PD1KO+TAC+*anti-CD8 Ab*, pink bars) or without depletion of CD8$^{+}$ T cells (PD1KO+TAC+iso Ab, green bars). **(B)** Echocardiographic analysis of LV dimensions at end-systole and end-diastole. **(C)** Assessment of early cardiac hypertrophy and pulmonary congestion via LV weight-to-tibial length (LV/TL), lung weight-to-tibial length (Lung/TL), and right ventricular weight-to-tibial length (RV/TL) ratios. **(D)** Representative flow cytometry plots identifying CD4$^{+}$ and CD8$^{+}$ T cell populations in the lymph node (LN) following *in vivo* treatment with either an IgG Isotype control (left) or a CD8-depleting antibody (right) for 7 days following TAC. The accompanying bar graph (right) quantifies depletion efficacy, showing the relative percentage of CD8$^{+}$ cells in the total CD3$^{+}$ T cell population. **(E)** Representative immunofluorescence images and corresponding quantification of infiltrating CD45$^{+}$ (yellow; left panels) and cytotoxic CD8$^{+}$ (red; right panels) cells per field in LV sections. Nuclei were counterstained with DAPI (blue). The accompanying bar graph quantifies the infiltrating CD45$^{+}$ and CD8$^{+}$ cells per field in the myocardium. Scale bar = 20 µm. **(F)** Representative immunofluorescence images and corresponding quantification of infiltrating CD45$^{+}$ and CD8$^{+}$ cells per field in lung sections. Scale bar = 20 µm. Data are presented as mean ± SEM. Statistical significance was determined by unpaired two-tailed Student's t-test, except that Welch's t-test was used for Lung/TL ratio **(C)**, % of CD8$^{+}$ T cells **(D)**, and lung infiltrating CD8 T cells **(F)**, and Mann-Whitney test was used for LV infiltrating CD8 T cells **(E)**. ns = not significant, $*p < 0.05$, $**p < 0.01$, $***p < 0.001$. n = 5-12 mice per group.

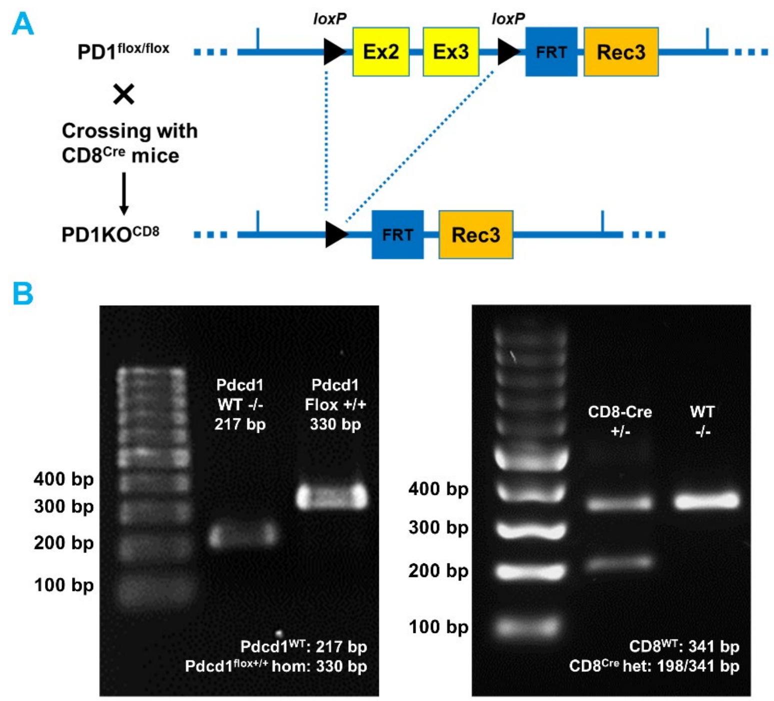


**Supplementary Figure S3. Diagram of CD8 cell specific PD1 KO mice and represented PCR for genotyping on the KO and WT strains.**
**(A)** Schematic diagram illustrating the genetic strategy used to generate CD8-specific PD-1 knockout mice ($PD1KO^{CD8}$). Mice possessing *loxP* sites flanking exons 2 and 3 of the *Pdcd1* gene ($PD1^{flox/flox}$) were crossed with mice expressing Cre recombinase under the control of the CD8 promoter, leading to the targeted excision of the floxed region. **(B)** Representative agarose gel images of PCR genotyping results. The left gel demonstrates genotyping for the *Pdcd1* allele, distinguishing the wild-type (WT) band at 217 bp from the homozygous floxed band ($PD1^{f/f}$) at 330 bp. The right gel demonstrates genotyping for the CD8-Cre transgene, identifying the WT allele by a single 341 bp band and the hemizygous CD8-Cre genotype by the presence of both 198 bp and 341 bp bands.

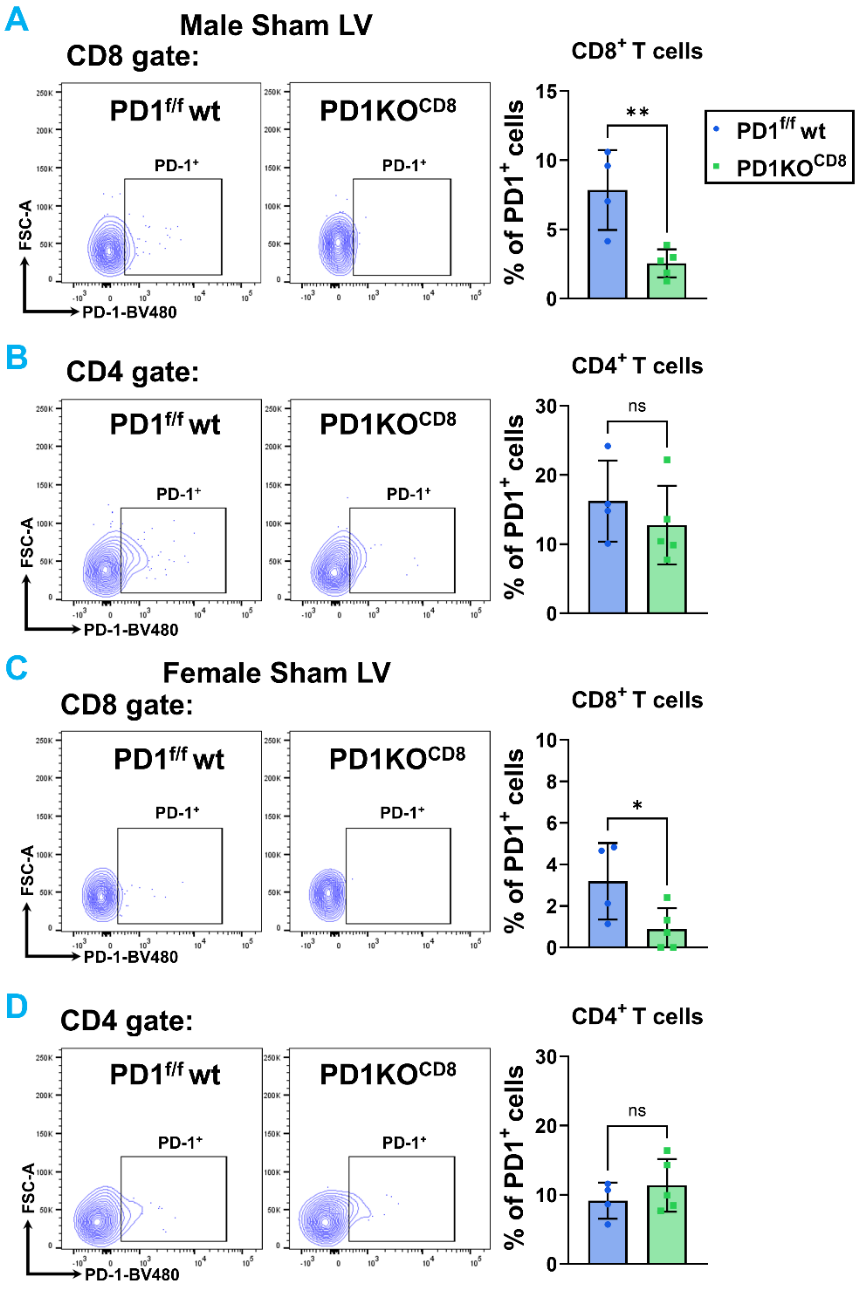


**Supplementary Figure S4. Validation of CD8$^+$ T cell-specific PD-1 deletion in cardiac tissue.**

**(A-B)** Representative flow cytometry plots and quantification of PD-1$^+$ cells among cardiac CD8$^+$ and CD4$^+$ T cells in the LV of male PD1$^{f/f}$ WT (blue) and PD1KO$^{CD8}$ (green) mice. **(C-D)** Representative flow cytometry plots and quantification of PD-1$^+$ cells among cardiac CD8$^+$ and CD4$^+$ T cells in the LV of female PD1$^{f/f}$ WT (blue) and PD1KO$^{CD8}$ (green) mice. Data are presented as mean ± SEM. Statistical significance was determined by unpaired two-tailed Student's t-test. ns = not significant, *p < 0.05, **p < 0.01. n = 4-5 mice per group.

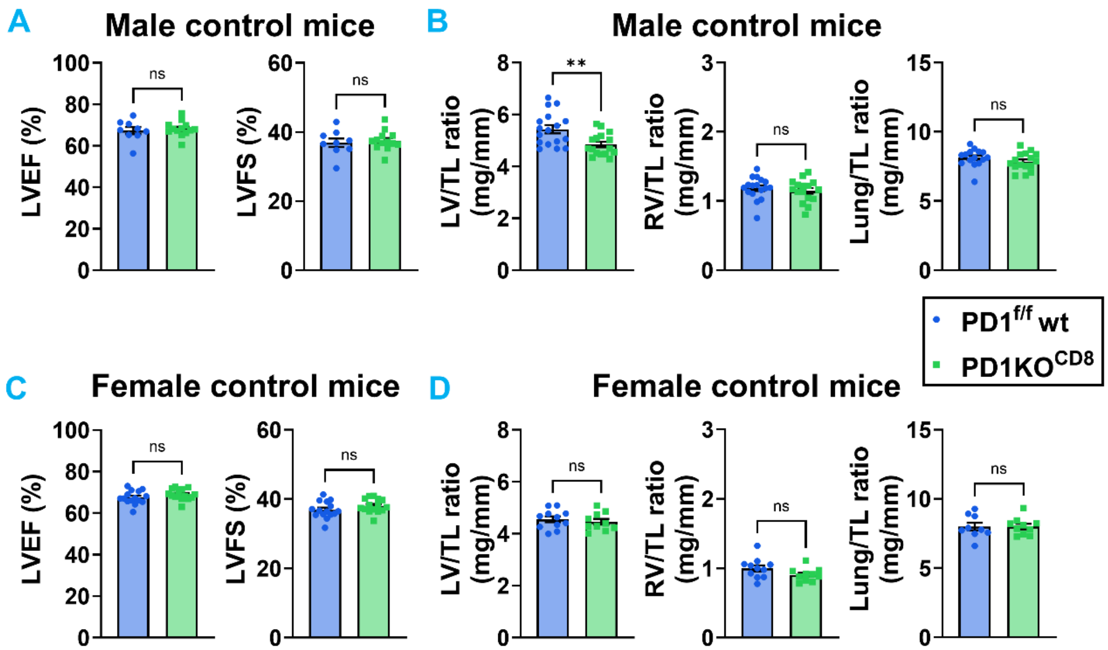


**Supplementary Figure S5. Baseline echocardiographic assessment and tissue weights in male and female mice.**

**(A)** Quantitative analysis of baseline systolic function in male PD1$^{f/f}$ wt (blue) and PD1KO$^{CD8}$ (green) mice, including LV ejection fraction (LVEF%) and fractional shortening (LVFS%). **(B)** Quantitative analysis of baseline tissue weights in male mice to assess baseline organ morphometry. **(C)** Quantitative analysis of baseline systolic function (LVEF% and LVFS%) in female PD1$^{f/f}$ wt (blue) and PD1KO$^{CD8}$ (green) mice. **(F)** Quantitative analysis of baseline tissue weights in female mice. Data are presented as mean ± SEM. Statistical significance was determined by unpaired two-tailed Student's t-test. ns = not significant, **$p < 0.01$. n = 5-17 mice per group.

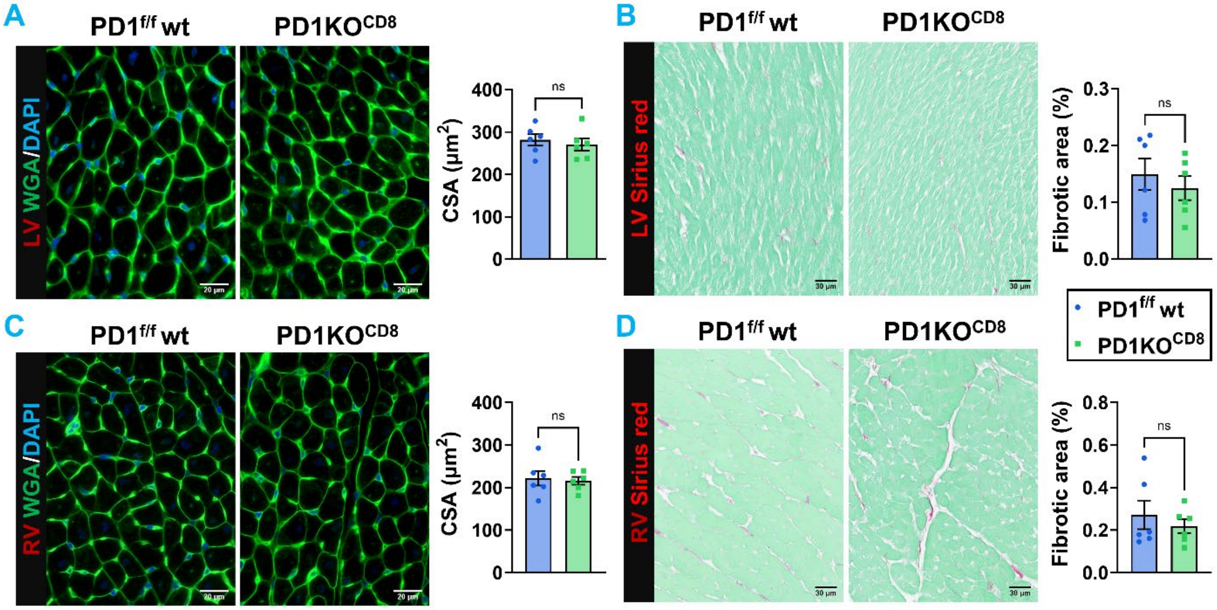


**Supplementary Figure S6. Baseline morphometric and histological analysis of left and right ventricular tissues.**

**(A)** Representative immunofluorescence images of LV sections from baseline PD1$^{f/f}$ wt (blue) and PD1KO$^{CD8}$ (green) mice. Tissues were stained with Wheat Germ Agglutinin (WGA, green) to delineate cardiomyocyte membranes and assess cell size. Nuclei were counterstained with DAPI (blue). The accompanying bar graph quantifies the LV cardiomyocyte cross-sectional area. Scale bar = 20 µm. **(B)** Representative images of LV sections stained with Sirius red to visualize baseline collagen deposition, alongside the corresponding quantification of the fibrotic area. Scale bar = 30 µm. **(C)** Representative WGA/DAPI staining and corresponding quantification of cardiomyocyte cross-sectional area in right ventricular (RV) sections. Scale bar = 20 µm. **(D)** Representative Sirius red staining and corresponding quantification of fibrotic area in RV sections. Scale bar = 30 µm. Data are presented as mean ± SEM. Statistical significance was determined by unpaired two-tailed Student's t-test. ns = not significant. n = 6 mice per group.

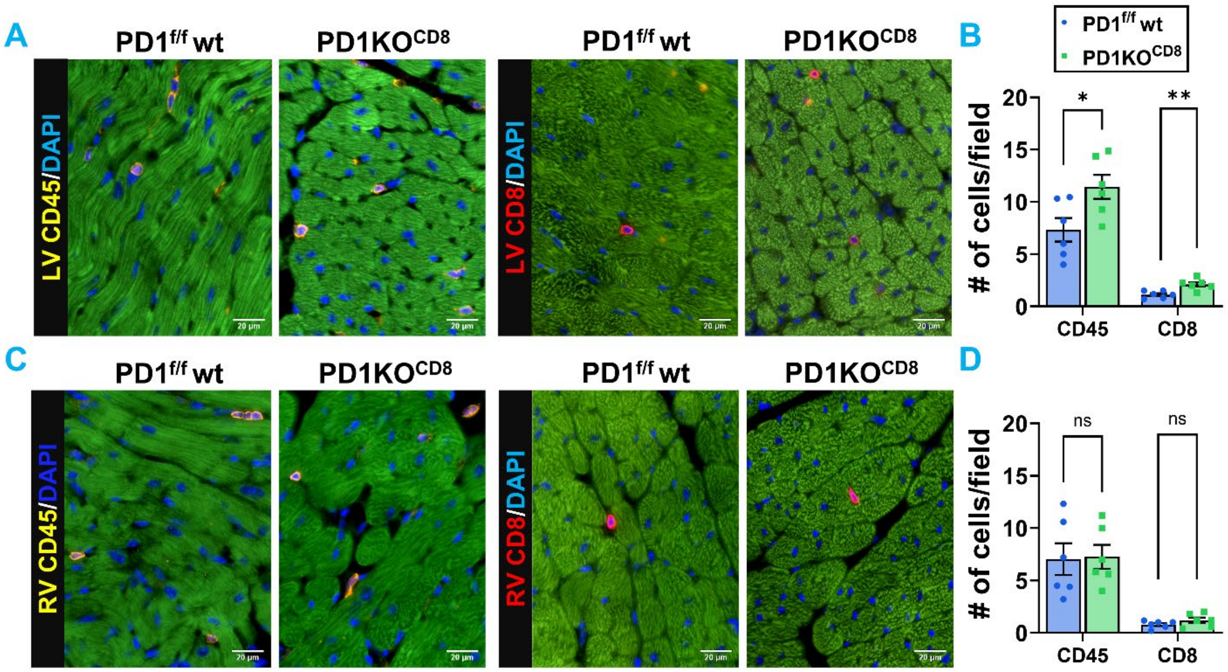


**Supplementary Figure S7. Baseline immune cell infiltration in left and right ventricular tissues.**

**(A)** Representative immunofluorescence images of LV sections from PD1$^{f/f}$ wt (blue) and PD1KO$^{CD8}$ (green) mice at baseline. Tissues were stained for CD45 (left panels) to identify total leukocytes and CD8 (right panels) to identify cytotoxic T cells. Nuclei were counterstained with DAPI (blue). **(B)** Quantitative analysis of infiltrating CD45$^+$ and CD8$^+$ cells per field in the LV corresponding to the images in panel **A**. **(C)** Representative immunofluorescence images of RV sections from PD1$^{f/f}$ wt (blue) and PD1KO$^{CD8}$ (green) mice, stained for CD45 (left panels) and CD8 (right panels). **(D)** Quantitative analysis of infiltrating CD45$^+$ and CD8$^+$ cells per field in the RV corresponding to the images in panel **C**. Scale bar = 20 µm. Data are presented as mean ± SEM. Statistical significance was determined by unpaired two-tailed Student's t-test. ns = not significant. n = 6 mice per group.

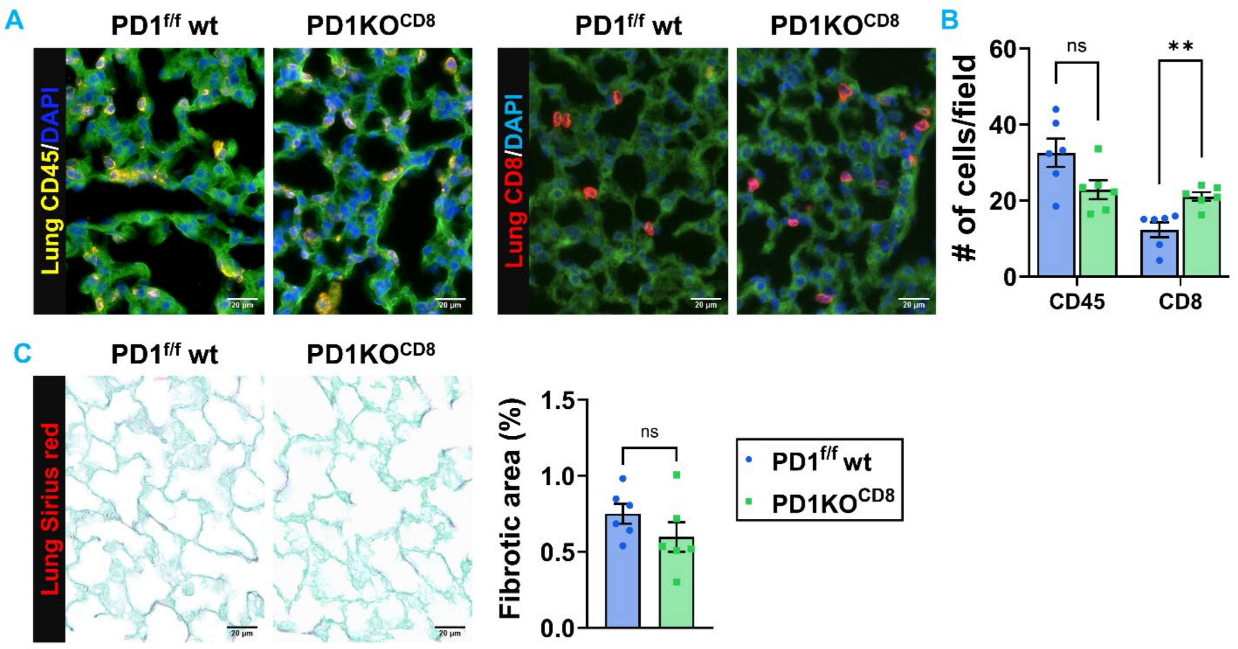


**Supplementary Figure S8. Baseline immune cell infiltration in male lung tissues.**

**(A)** Representative immunofluorescence images of lung sections from male mice at baseline. Tissues were stained for CD45 (left panels, yellow) to identify total leukocytes and CD8 (right panels, red) to identify cytotoxic T cells. Nuclei were counterstained with DAPI (blue). **(B)** Quantitative analysis of infiltrating $CD45^+$ and $CD8^+$ cells per field in the lung corresponding to the images in panel A. **(C)** Representative images of lung sections stained with Sirius red to visualize baseline collagen deposition, alongside the corresponding quantification of the fibrotic area. Scale bar = 20 µm. Data are presented as mean ± SEM. Statistical significance was determined by unpaired two-tailed Student's t-test, except that Mann-Whitney test was used for $CD8^+$ T cells in Panel **(B)**. ns = not significant, $^{**}p < 0.01$. n = 6 mice per group.

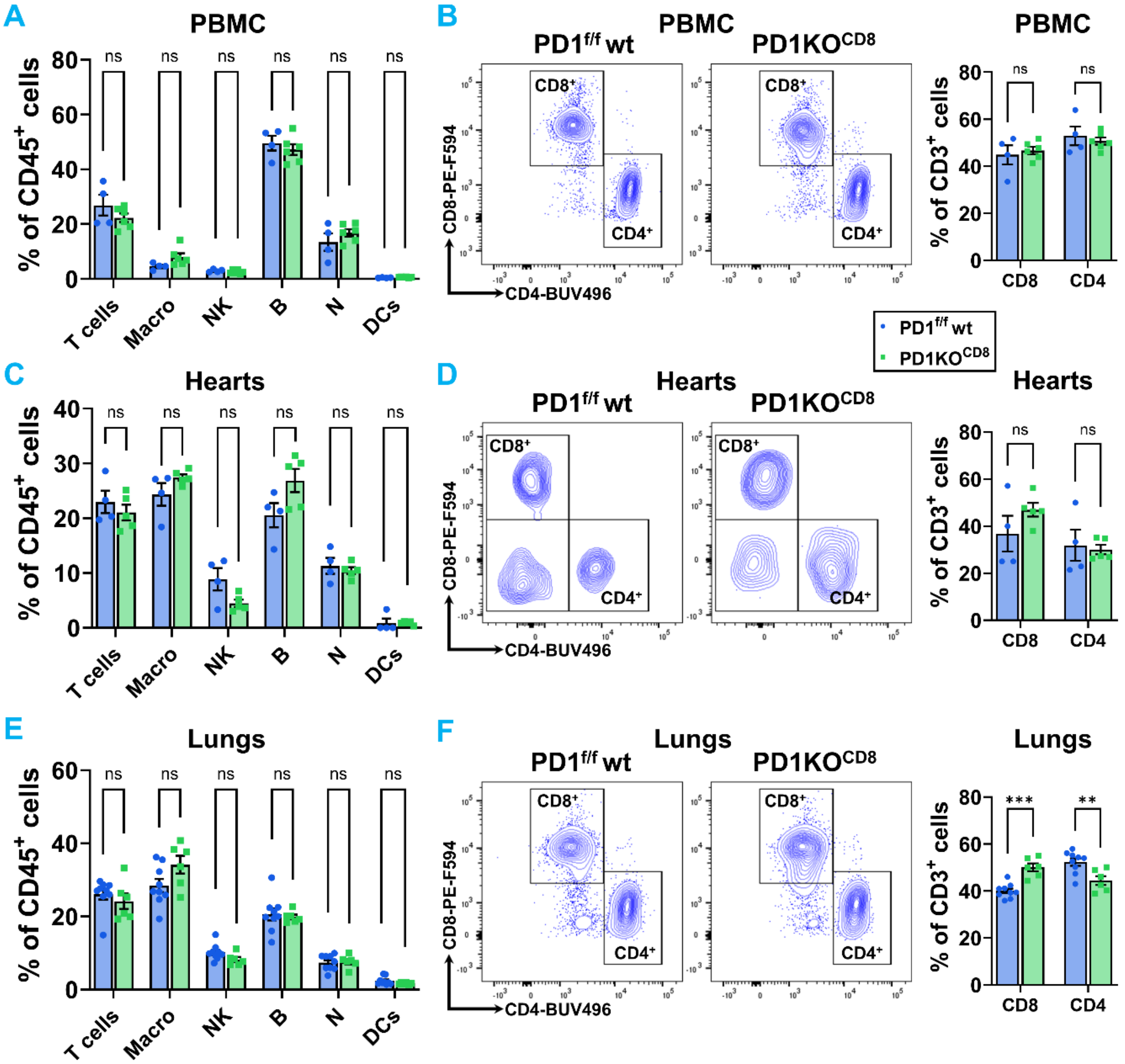


**Supplementary Figure S9. Flow cytometric analysis of immune cell populations in blood, cardiac tissues, and lung tissues of male PD1$^{f/f}$ wt and PD1KO$^{CD8}$ mice.**

**(A, C, E)** Flow cytometric quantification of major immune cell subsets—including T cells, macrophages (Macro), Natural Killer (NK) cells, B cells, neutrophils (N), and dendritic cells (DCs)—expressed as a percentage of total CD45$^{+}$ leukocytes in peripheral blood mononuclear cells (PBMC; **A**), cardiac tissues (Hearts; **C**), and lung tissues (Lungs; **E**) of PD1$^{f/f}$ wt (blue) and PD1KO$^{CD8}$ (green) mice at baseline. **(B, D, F)** Representative flow cytometry contour plots identifying CD8$^{+}$ and CD4$^{+}$ T cell populations within the PBMC **(B)**, Hearts **(D),** and Lungs **(F)** of PD1$^{f/f}$ wt and PD1KO$^{CD8}$ mice, and corresponding quantitative analysis of CD8$^{+}$ and CD4$^{+}$ T cell subsets expressed as a percentage of total CD3$^{+}$ T cells in PBMC, Hearts, and Lungs. Data are presented as mean ± SEM. Statistical significance was determined by unpaired two-tailed Student's t-test, except that Welch's t-test was used for macrophage **(C)** and B cell **(E)**, and that Mann-Whitney test was used for DCs in panels **(C)** and **(E)**. ns = not significant; $^{**}p < 0.01$, $^{***}p < 0.001$. n = 4-9 mice per group.

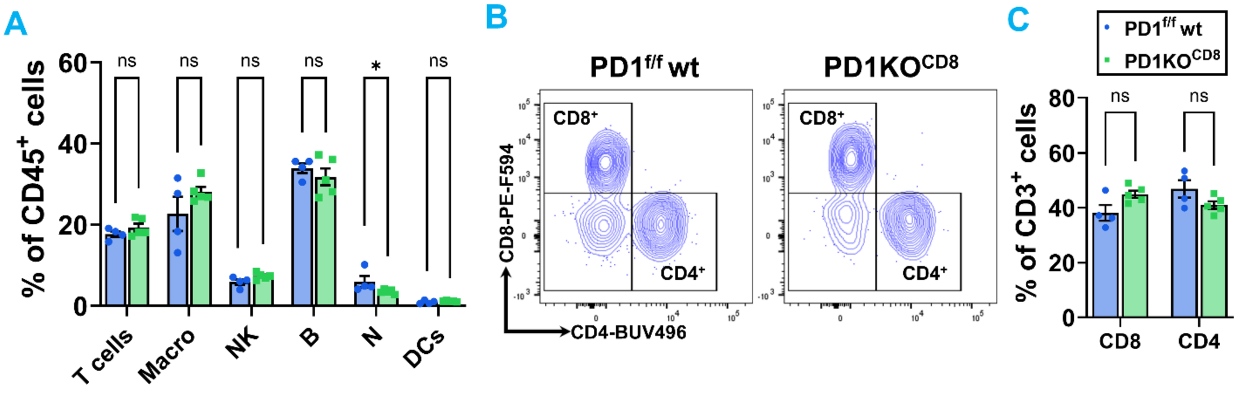


**Supplementary Figure S10. Flow cytometric analysis of LV immune cell populations in female PD1$^{f/f}$ wt and PD1KO$^{CD8}$ mice.**

**(A)** Flow cytometric quantification of major immune cell subsets—including T cells, macrophages (Macro), Natural Killer (NK) cells, B cells, neutrophils (N), and dendritic cells (DCs)—expressed as a percentage of total CD45$^{+}$ leukocytes in LV tissues of female PD1$^{f/f}$ wt (blue) and PD1KO$^{CD8}$ (green) mice. **(B)** Representative flow cytometry contour plots identifying CD8$^{+}$ and CD4$^{+}$ T cell populations within the LV of PD1$^{f/f}$ wt (left) and PD1KO$^{CD8}$ (right) mice. **(C)** Quantitative analysis of CD8$^{+}$ and CD4$^{+}$ T cell subsets expressed as a percentage of total CD3$^{+}$ T cells. Data are presented as mean ± SEM. Statistical significance was determined by unpaired two-tailed Student's t-test, except that Mann-Whitney test was used for neutrophil in panel **(A)**. ns = not significant; *$p < 0.05$. n = 4-5 mice per group.

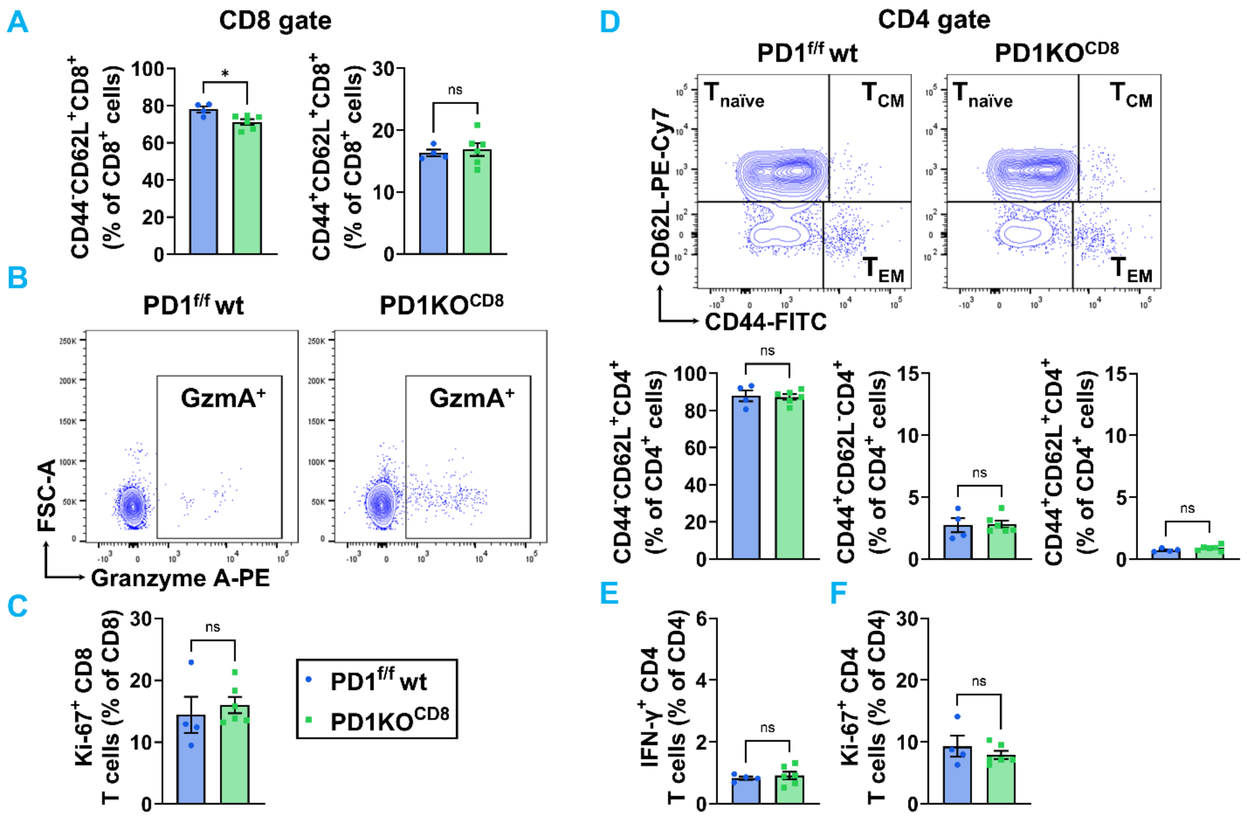


**Supplementary Figure S11. Flow cytometric analysis of activation status and cytokine production in peripheral T cells at baseline.**

**(A)** Summarized data of $CD8^+$ T cell subsets isolated from PBMC of male $PD1^{f/f}$ WT (blue) and $PD1KO^{CD8}$ (green) mice. Subsets are phenotypically defined by surface marker expression as naïve ($T_{naïve}$) and central memory ($T_{CM}$). **(B)** Representative flow cytometry contour plots of Granzyme $A^+CD8^+$ T cell subsets isolated from PBMC of $PD1^{f/f}$ WT (blue) and $PD1KO^{CD8}$ (green) mice. **(C)** Summarized data of proliferation marker, Ki-67, in $CD8^+$ T cell subsets isolated from PBMC of $PD1^{f/f}$ WT (blue) and $PD1KO^{CD8}$ (green) mice. **(D)** Representative flow cytometry contour plots and quantitative analysis of $CD4^+$ T cell subsets isolated from PBMC of $PD1^{f/f}$ WT (blue) and $PD1KO^{CD8}$ (green) mice. Subsets are phenotypically defined by surface marker expression as naïve ($T_{naïve}$), central memory ($T_{CM}$), and effector memory ($T_{EM}$). The data demonstrate no phenotypic changes in $CD4^+$ T cell subsets. **(E–F)** Summarized data depicting the intracellular production of the pro-inflammatory cytokine IFN-γ and proliferation marker, Ki-67 in $CD4^+$ T cells from PBMC. Data are presented as mean ± SEM. Statistical significance was determined by unpaired two-tailed Student's t-test. ns = not significant, $*p < 0.05$, $**p < 0.01$. n = 4-6 mice per group.

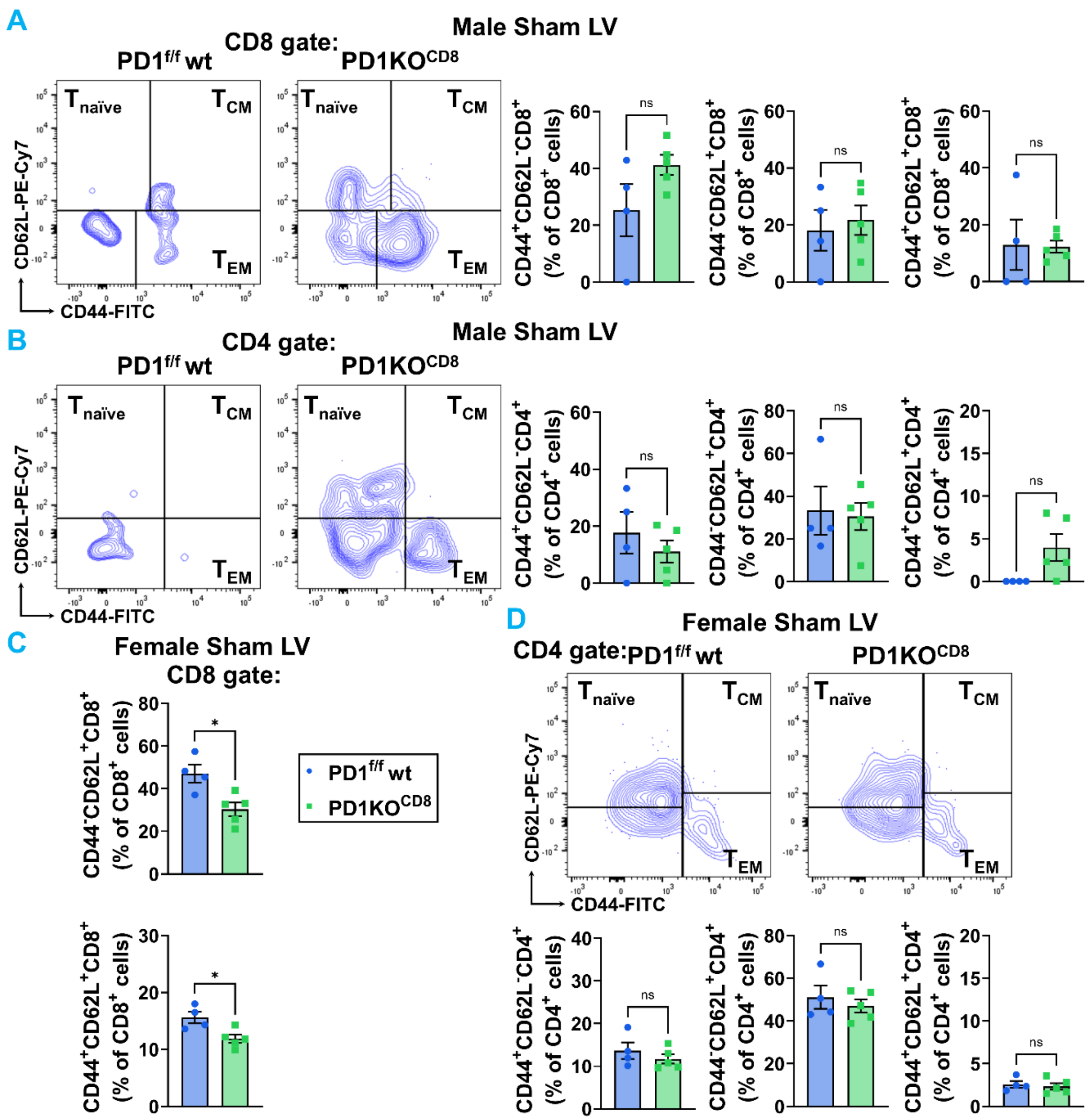


**Supplementary Figure S12. Flow cytometric analysis of activation status in cardiac T cells in $CD8^+$ T cell-specific PD-1-deficient mice at baseline.**

**(A-B)** Representative flow cytometry plots and quantification of naïve ($T_{\text{naïve}}$), effector memory ($T_{EM}$), and central memory ($T_{CM}$) subsets, gated by CD62L and CD44 expression among cardiac $CD8^+$ (A) and $CD4^+$ (B) T cells in the LV of male $PD1^{f/f}$ WT (blue) and $PD1KO^{CD8}$ (green) mice at baseline. **(C-D)** Representative flow cytometry plots and quantification of naïve ($T_{\text{naïve}}$), effector memory ($T_{EM}$), and central memory ($T_{CM}$) subsets, gated by CD62L and CD44 expression among cardiac $CD8^+$ (**C**) and $CD4^+$ (**D**) T cells in the LV of female $PD1^{f/f}$ WT (blue) and $PD1KO^{CD8}$ (green) mice at baseline. Data are presented as mean ± SEM. Statistical significance was determined by unpaired two-tailed Student's t-test, except that Welch's t-test was used for $T_{CM}$ in $CD8^+$ **(A)** and $CD4^+$ **(B)** T cells. ns = not significant, *$p < 0.05$, **$p < 0.01$. n = 4-5 mice per group.

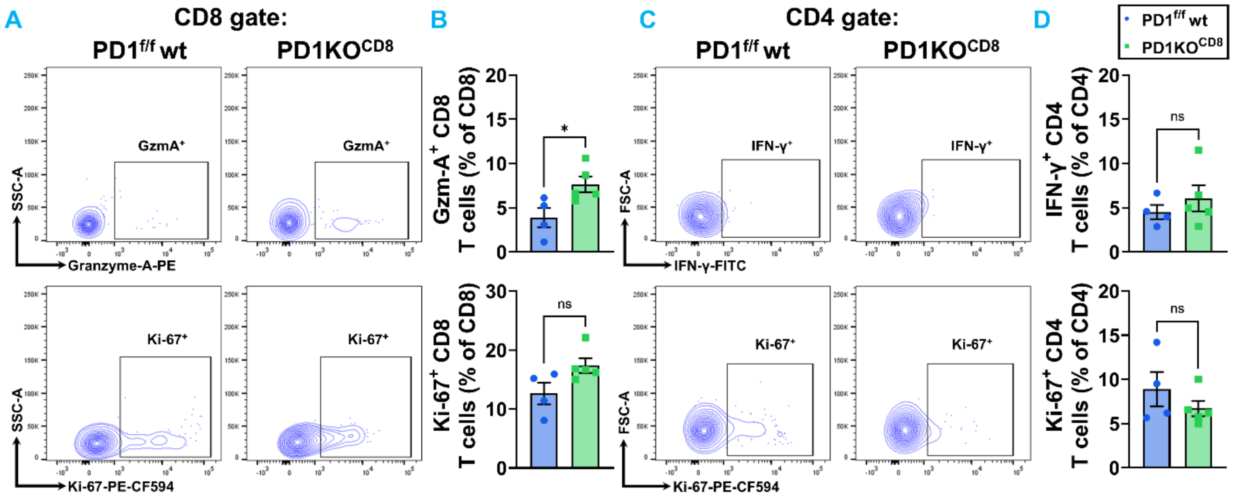


**Supplementary Figure S13. Flow cytometric analysis of cytokine production in cardiac T cells in female CD8$^+$ T cell-specific PD-1-deficient mice at baseline.**

**(A-B)** Representative flow cytometry contour plots of Granzyme A$^+$ and Ki-67$^+$ CD8$^+$ T cell subsets isolated from the LV of female PD1$^{f/f}$ WT (blue) and PD1KO$^{CD8}$ (green) mice. **(C-D)** Representative flow cytometry contour plots of IFN-γ$^+$ and Ki-67$^+$ CD4$^+$ T cell subsets isolated from the LV of female PD1$^{f/f}$ WT (blue) and PD1KO$^{CD8}$ (green) mice. Data are presented as mean ± SEM. Statistical significance was determined by unpaired two-tailed Student's t-test. ns = not significant, *$p < 0.05$. n = 4-5 mice per group.

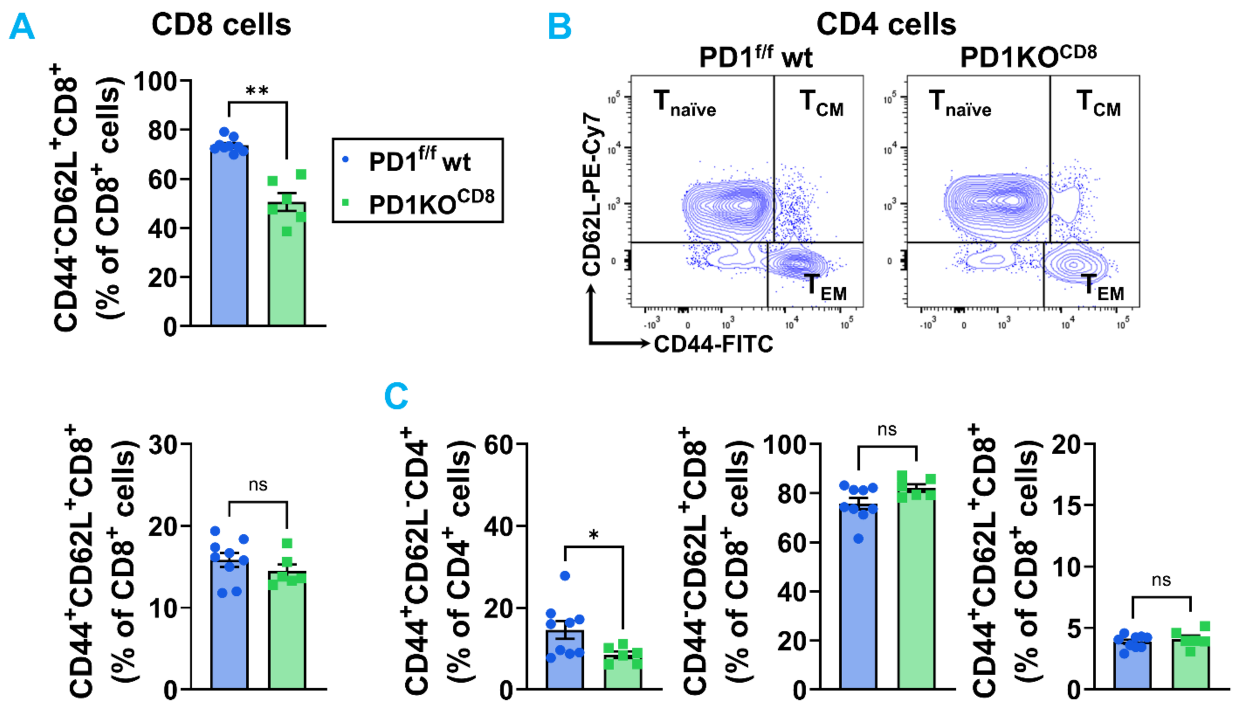


**Supplementary Figure S14. Flow cytometric analysis of activation status in pulmonary T cells in CD8⁺ T cell-specific PD-1-deficient mice at baseline.**

**(A)** Quantification of naïve ($T_{naïve}$) and central memory ($T_{CM}$) subsets, gated by CD62L and CD44 expression among pulmonary CD8⁺ T cells in the male PD1$^{f/f}$ wt (blue) and PD1KO$^{CD8}$ (green) mice at baselinee. **(B-C)** Representative flow cytometry plots and quantification of naïve ($T_{naïve}$), effector memory ($T_{EM}$), and central memory ($T_{CM}$) subsets, among pulmonary CD4⁺ T cells in the male PD1$^{f/f}$ wt (blue) and PD1KO$^{CD8}$ (green) mice at baseline. Data are presented as mean ± SEM. Statistical significance was determined by unpaired two-tailed Student's t-test, except that Welch's t-test was used for $T_{EM}$ and $T_{naïve}$ in CD8⁺ **(A)** and $T_{EM}$ in CD4⁺ **(C)** T cells. ns = not significant, *$p < 0.05$, **$p < 0.01$. n = 6-9 mice per group.

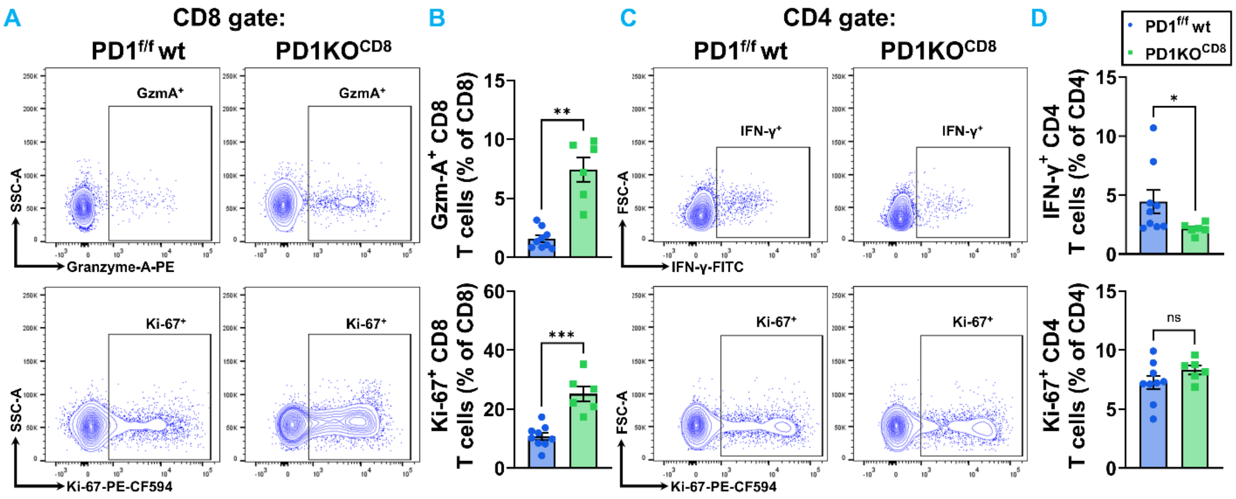


**Supplementary Figure S15. CD8-specific PD-1 deficiency significantly enhances intracellular Granzyme A production and proliferation in pulmonary CD8$^+$ T cells at baseline.**

**(A-B)** Representative flow cytometry contour plots of Granzyme A$^+$ and Ki-67$^+$ CD8$^+$ T cell subsets isolated from the lung of male PD1$^{f/f}$ WT (blue) and PD1KO$^{CD8}$ (green) mice. **(C-D)** Representative flow cytometry contour plots of IFN-γ$^+$ and Ki-67$^+$ CD4$^+$ T cell subsets isolated from the lung of male PD1$^{f/f}$ WT (blue) and PD1KO$^{CD8}$ (green) mice. Data are presented as mean ± SEM. Statistical significance was determined by unpaired two-tailed Student's t-test, except that Welch's t-test was used for Granzyme A$^+$CD8$^+$ T cells in panel **(B)** and Mann Whitney test was used for IFN-γ$^+$CD4$^+$ T cells in Panel **(D)**. ns = not significant, *$p < 0.05$, **$p < 0.01$, ***$p < 0.001$. n = 6-9 mice per group.

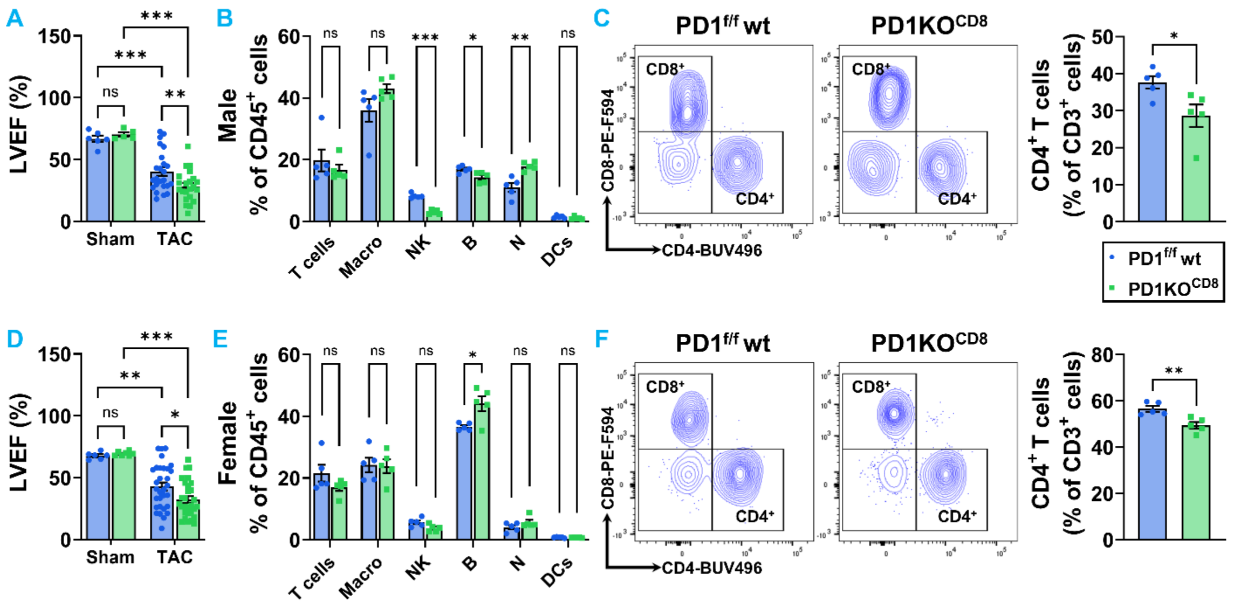


**Supplementary Figure S16. Flow cytometric analysis of LV immune cell populations in male and female PD1$^{f/f}$ wt and PD1KO$^{CD8}$ mice.**

**(A and D)** Quantitative echocardiographic analysis of LVEF% in male (**A**) and female (**D**) wild-type (PD1$^{f/f}$ wt, blue) and CD8$^{+}$ T cell-specific PD-1 knockout (PD1KO$^{CD8}$, green) mice following sham or transverse aortic constriction (TAC) surgery. **(E and F)** Flow cytometric quantification of major immune cell subsets—including T cells, macrophages (Macro), Natural Killer (NK) cells, B cells, neutrophils (N), and dendritic cells (DCs)—expressed as a percentage of total CD45$^{+}$ leukocytes in LV tissues in male **(A)** and female **(D)** mice after TAC. **(B and E)** Representative flow cytometry contour plots identifying CD8$^{+}$ and CD4$^{+}$ T cell populations within the LV of male **(B)** and female **(E)** PD1$^{f/f}$ wt and PD1KO$^{CD8}$ mice after TAC. **(C and F)** Quantitative analysis of CD4$^{+}$ T cell subsets expressed as a percentage of total CD3$^{+}$ T cells in the LV of male **(C)** and female **(F)** PD1$^{f/f}$ wt and PD1KO$^{CD8}$ mice after TAC. Data are presented as mean ± SEM. Statistical significance was determined by unpaired two-tailed Student's t-test, except that two-way ANOVA with Tukey's multiple comparisons test was used for panels **A** and **D**, Welch's t-test was used for B cells in panel **(B),** and Mann Whitney test was used for T cells, macrophages, Panel **(C)**. ns = not significant, *p < 0.05, **p < 0.01, ***p < 0.001.  n = 5-31 mice per group.

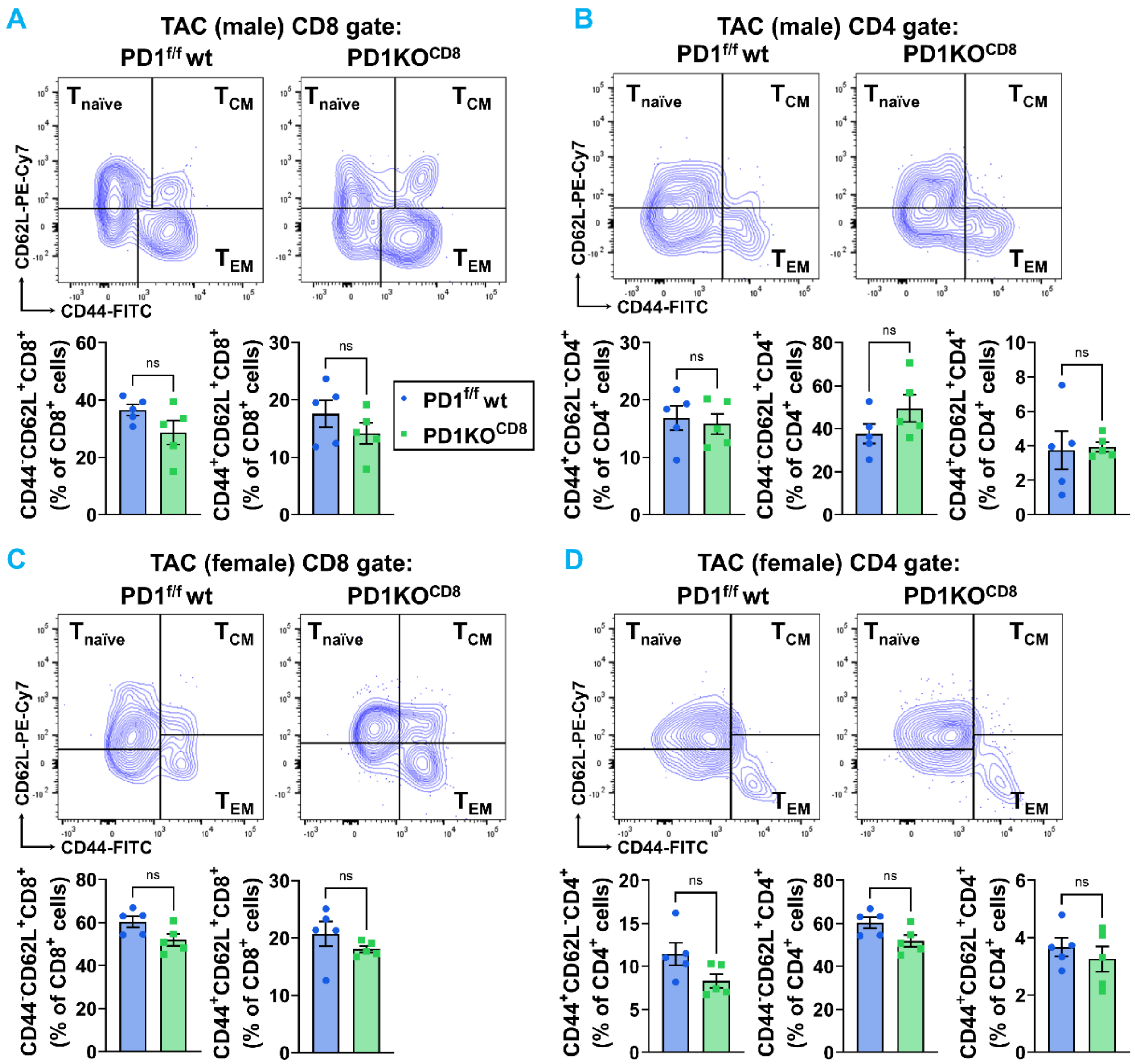


**Supplementary Figure S17. Flow cytometric analysis of activation status in cardiac T cells in CD8⁺ T cell-specific PD-1-deficient mice after TAC.**

**(A-B)** Representative flow cytometry plots and quantification of naïve ($T_{naïve}$), effector memory ($T_{EM}$), and central memory ($T_{CM}$) subsets, gated by CD62L and CD44 expression among cardiac $CD8^+$ **(A)** and $CD4^+$ **(B)** T cells in the LV of male $PD1^{f/f}$ WT (blue) and $PD1KO^{CD8}$ (green) mice after TAC. **(C-D)** Representative flow cytometry plots and quantification of naïve ($T_{naïve}$), effector memory ($T_{EM}$), and central memory ($T_{CM}$) subsets, gated by CD62L and CD44 expression among cardiac $CD8^+$ **(C)** and $CD4^+$ **(D)** T cells in the LV of female $PD1^{f/f}$ WT (blue) and $PD1KO^{CD8}$ (green) mice after TAC. Data are presented as mean ± SEM. Statistical significance was determined by unpaired two-tailed Student's t-test, except that Welch's t-test was used for $T_{CM}$ in $CD4^+$ **(B)** and $CD8^+$ **(C)** T cells. ns = not significant. n = 5 mice per group.

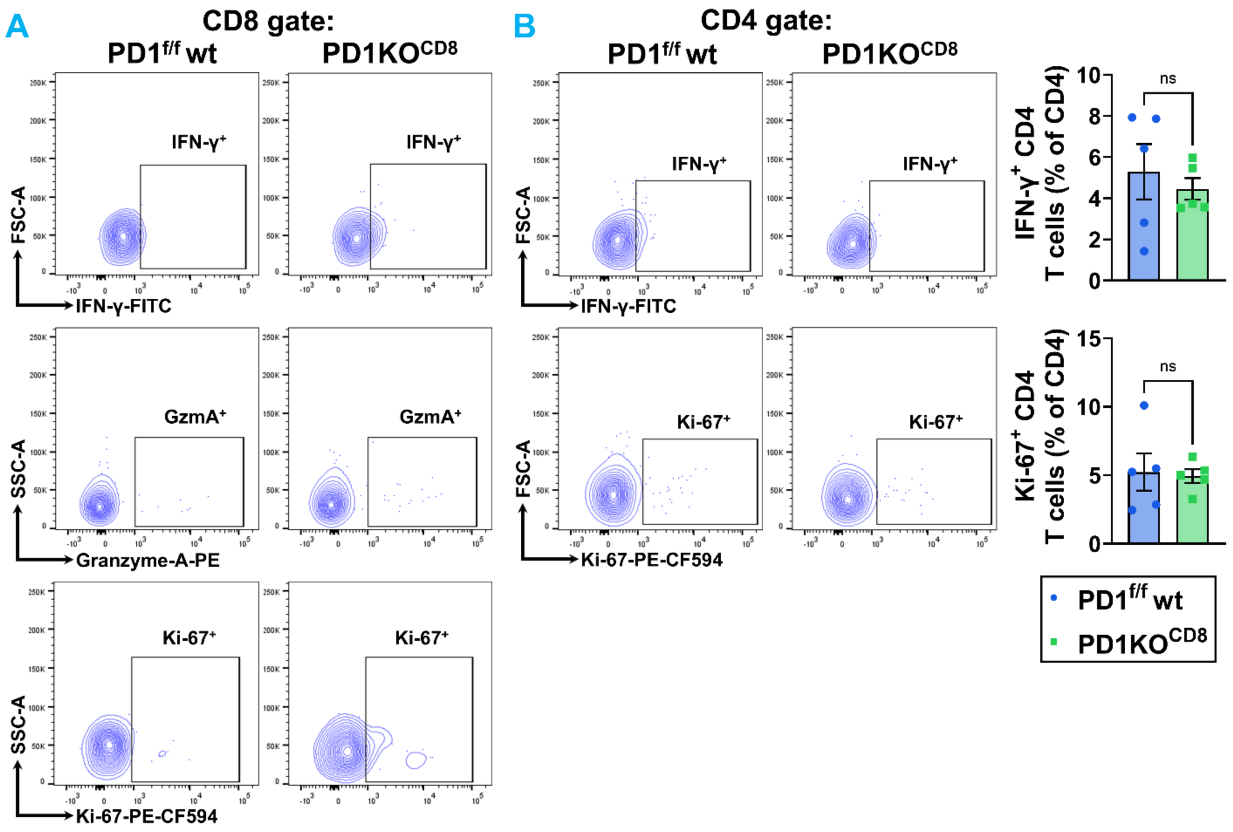


**Supplementary Figure S18. Flow cytometric analysis of effector function in cardiac T cells in male $CD8^+$ T cell-specific PD-1-deficient mice after TAC.**

**(A)** Representative flow cytometry contour plots of IFN-$\gamma^+$, Granzyme $A^+$, and Ki-$67^+$ $CD8^+$ T cell subsets isolated from the LV of male $PD1^{f/f}$ WT (blue) and $PD1KO^{CD8}$ (green) mice. **(B)** Representative flow cytometry contour plots and analysis of IFN-$\gamma^+$ and Ki-$67^+$ $CD4^+$ T cell subsets isolated from the LV of male $PD1^{f/f}$ WT (blue) and $PD1KO^{CD8}$ (green) mice. Data are presented as mean ± SEM. Statistical significance was determined by unpaired two-tailed Student's t-test. ns = not significant. n = 5 mice per group.

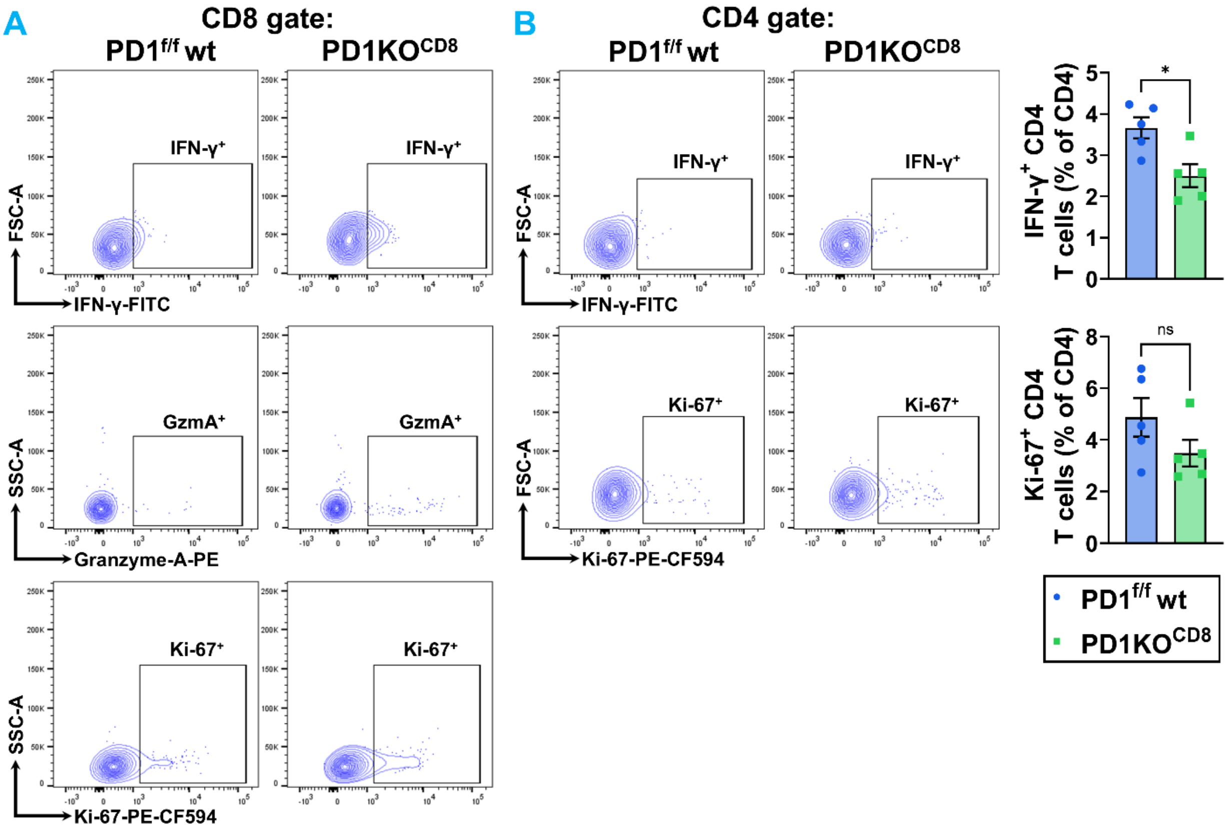


**Supplementary Figure S19. Flow cytometric analysis of effector function in cardiac T cells in female CD8$^{+}$ T cell-specific PD-1-deficient mice after TAC.**

**(A)** Representative flow cytometry contour plots of IFN-γ$^{+}$, Granzyme A$^{+}$, and Ki-67$^{+}$ CD8$^{+}$ T cell subsets isolated from the LV of female PD1$^{f/f}$ WT (blue) and PD1KO$^{CD8}$ (green) mice. **(B)** Representative flow cytometry contour plots and analysis of IFN-γ$^{+}$ and Ki-67$^{+}$ CD4$^{+}$ T cell subsets isolated from the LV of female PD1$^{f/f}$ WT (blue) and PD1KO$^{CD8}$ (green) mice. Data are presented as mean ± SEM. Statistical significance was determined by unpaired two-tailed Student's t-test. ns = not significant, *p < 0.05. n = 5 mice per group.

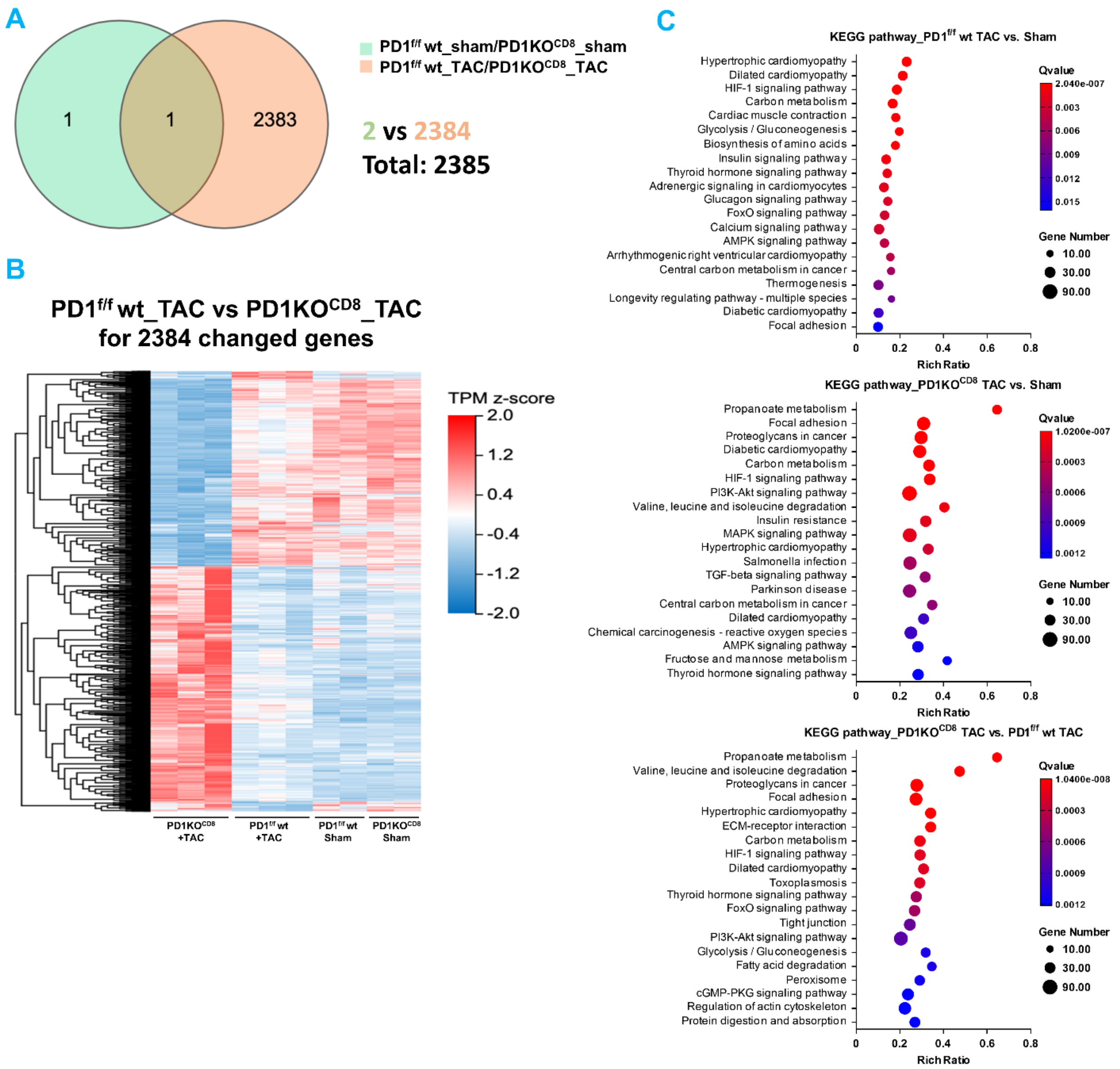


**Supplementary Figure S20. Bulk RNA-sequencing reveals distinct transcriptional signatures in PD1$^{f/f}$ wt and PD1KO$^{CD8}$ hearts following TAC.**

**(A)** Venn diagram depicting the number of significantly changed genes in the PD1$^{f/f}$ wt_sham vs. PD1KO$^{CD8}$ sham comparison (green) and the PD1$^{f/f}$ wt TAC vs. PD1KO$^{CD8}$ TAC comparison (orange), with overlapping genes shown in the intersection. **(B)** Heatmap of the 2,384 differentially expressed genes between PD1$^{f/f}$ wt TAC and PD1KO$^{CD8}$ TAC hearts across the four groups. **(C)** KEGG pathway enrichment analysis (rich ratio) of differentially expressed genes for the comparisons of PD1$^{f/f}$ wt TAC vs. sham (top), PD1KO$^{CD8}$ TAC vs. sham (middle), and PD1KO$^{CD8}$ TAC vs. PD1$^{f/f}$ wt TAC (bottom). RNA-sequencing was performed on cardiac tissue from PD1f/f wt and PD1KOCD8 mice under sham or TAC conditions. n=2-3 mice per group.

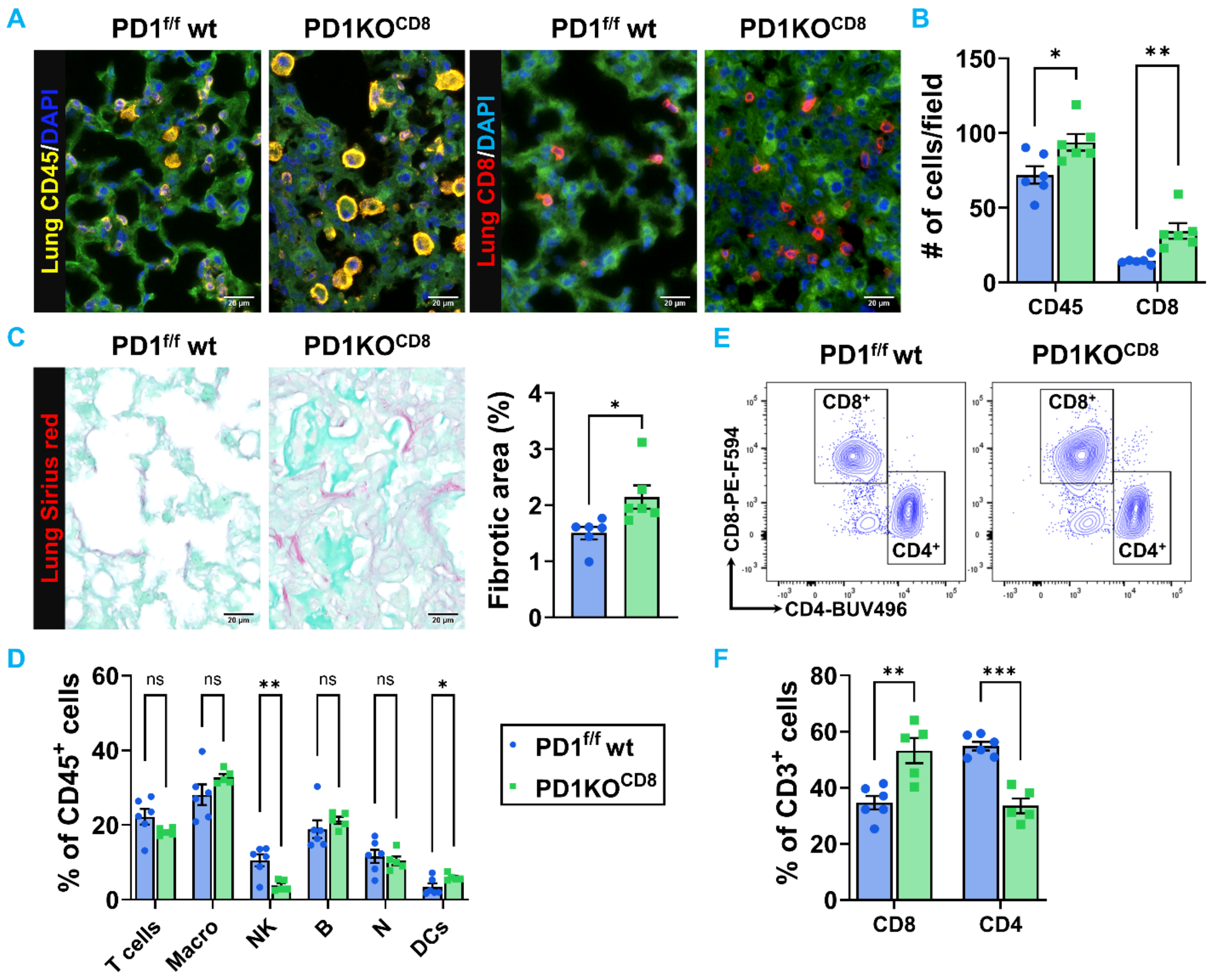


**Supplementary Figure S21. Immune cell infiltration in male lung tissues after TAC.**

**(A)** Representative immunofluorescence images of lung sections from male mice after TAC. Tissues were stained for CD45 (left panels, yellow) to identify total leukocytes and CD8 (right panels, red) to identify cytotoxic T cells. Nuclei were counterstained with DAPI (blue). **(B)** Quantitative analysis of infiltrating $CD45^+$ and $CD8^+$ cells per field in the lung corresponding to the images in panel **A**. **(C)** Representative images of lung sections stained with Sirius red to visualize baseline collagen deposition, alongside the corresponding quantification of the fibrotic area. **(D)** Flow cytometric quantification of major immune cell subsets—including T cells, macrophages (Macro), Natural Killer (NK) cells, B cells, neutrophils (N), and dendritic cells (DCs)—expressed as a percentage of total $CD45^+$ leukocytes isolated from lung tissue of male TAC-operated $PD1^{f/f}$ wt (blue) and $PD1KO^{CD8}$ (green) mice. **(E-F)** Representative flow cytometry contour plots identifying $CD8^+$ and $CD4^+$ T cell populations, with quantification expressed as the percentage of total $CD3^+$ T cells in lung tissue from male TAC-operated $PD1^{f/f}$ wt (blue) and $PD1KO^{CD8}$ (green) mice. Scale bar = 20 μm. Data are presented as mean ± SEM. Statistical significance was determined by unpaired two-tailed Student's t-test, except that Mann-Whitney test was used for $CD8^+$ T cells in Panel **(A)** and Welch's t-test was used for T cell, macrophages, and B cell in Panel **(D)**. ns = not significant, $^*p < 0.05$, $^{**}p < 0.01$, $^{***}p < 0.001$. n = 5-6 mice per group.

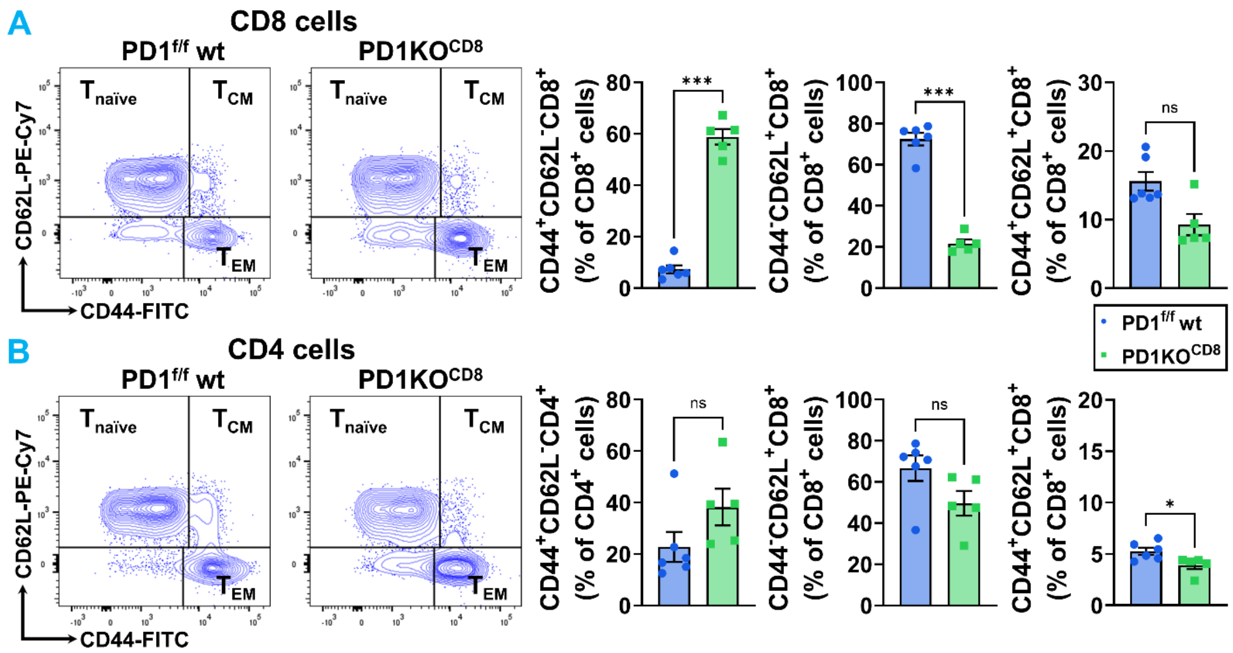


**Supplementary Figure S22. CD8-specific PD-1 deficiency significantly increases pulmonary $CD8^{+}$ T cell activation.**

**(A)** Representative flow cytometry plots and quantification of naïve ($T_{naïve}$), effector memory ($T_{EM}$), and central memory ($T_{CM}$) subsets, gated by CD62L and CD44 expression among pulmonary $CD8^{+}$ T cells in the male TAC-operated $PD1^{f/f}$ wt (blue) and $PD1KO^{CD8}$ (green) mice. **(B)** Representative flow cytometry plots and quantification of naïve ($T_{naïve}$), effector memory ($T_{EM}$), and central memory ($T_{CM}$) subsets, among pulmonary $CD4^{+}$ T cells in the male TAC-operated $PD1^{f/f}$ wt (blue) and $PD1KO^{CD8}$ (green) mice. Data are presented as mean ± SEM. Statistical significance was determined by unpaired two-tailed Student's t-test, except that Mann-Whitney test was used for $T_{CM}$ in $CD8^{+}$ **(A)** and $T_{EM}$ and $T_{naïve}$ in $CD4^{+}$ **(B)** T cells. ns = not significant, $*p < 0.05$, $***p < 0.001$. n = 5-6 mice per group.

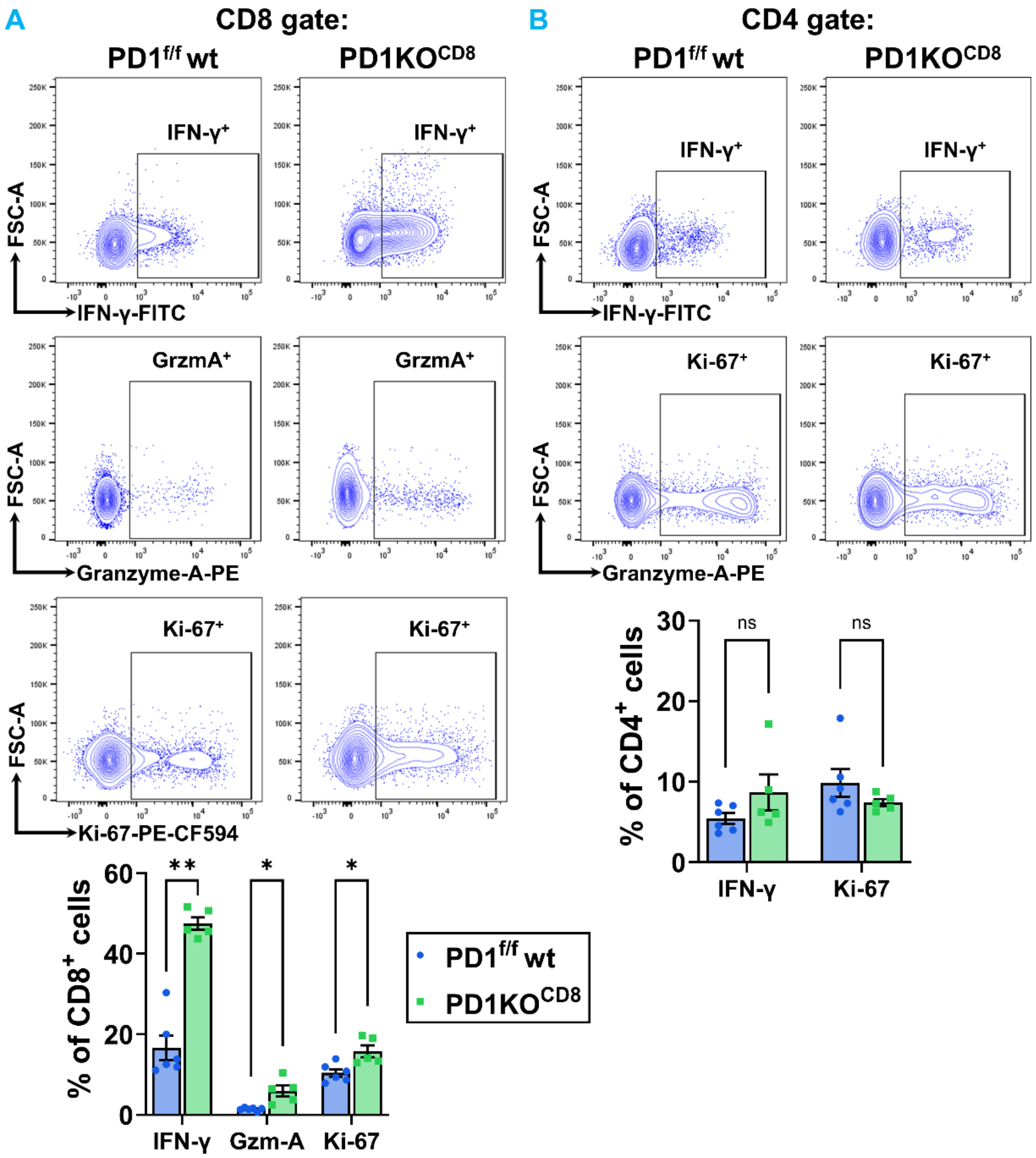


**Supplementary Figure S23. CD8-specific PD-1 deficiency significantly enhances intracellular cytokine production and proliferation in pulmonary CD8⁺ T cells after TAC.**

**(A)** Representative flow cytometry contour plots and analysis of IFN-$\gamma^+$, Granzyme $A^+$, and Ki-$67^+$ $CD8^+$ T cell subsets isolated from the lung of male $PD1^{f/f}$ wt (blue) and $PD1KO^{CD8}$ (green) mice. **B)** Representative flow cytometry contour plots of IFN-$\gamma^+$ and Ki-$67^+$ $CD4^+$ T cell subsets isolated from the lung of male $PD1^{f/f}$ wt (blue) and $PD1KO^{CD8}$ (green) mice. Data are presented as mean ± SEM. Statistical significance was determined by unpaired two-tailed Student's t-test, except that Welch's t-test was used for Granzyme $A^+CD8^+$ T cells in panel **(A)** and Mann Whitney test was used for IFN-$\gamma^+CD8^+$ T cells in panel **(A)**, and IFN-$\gamma^+CD4^+$ and Ki-$67^+CD4^+$ T cells in panel **(B)**. ns = not significant, *$p < 0.05$, **$p < 0.01$. n = 5-6 mice per group.

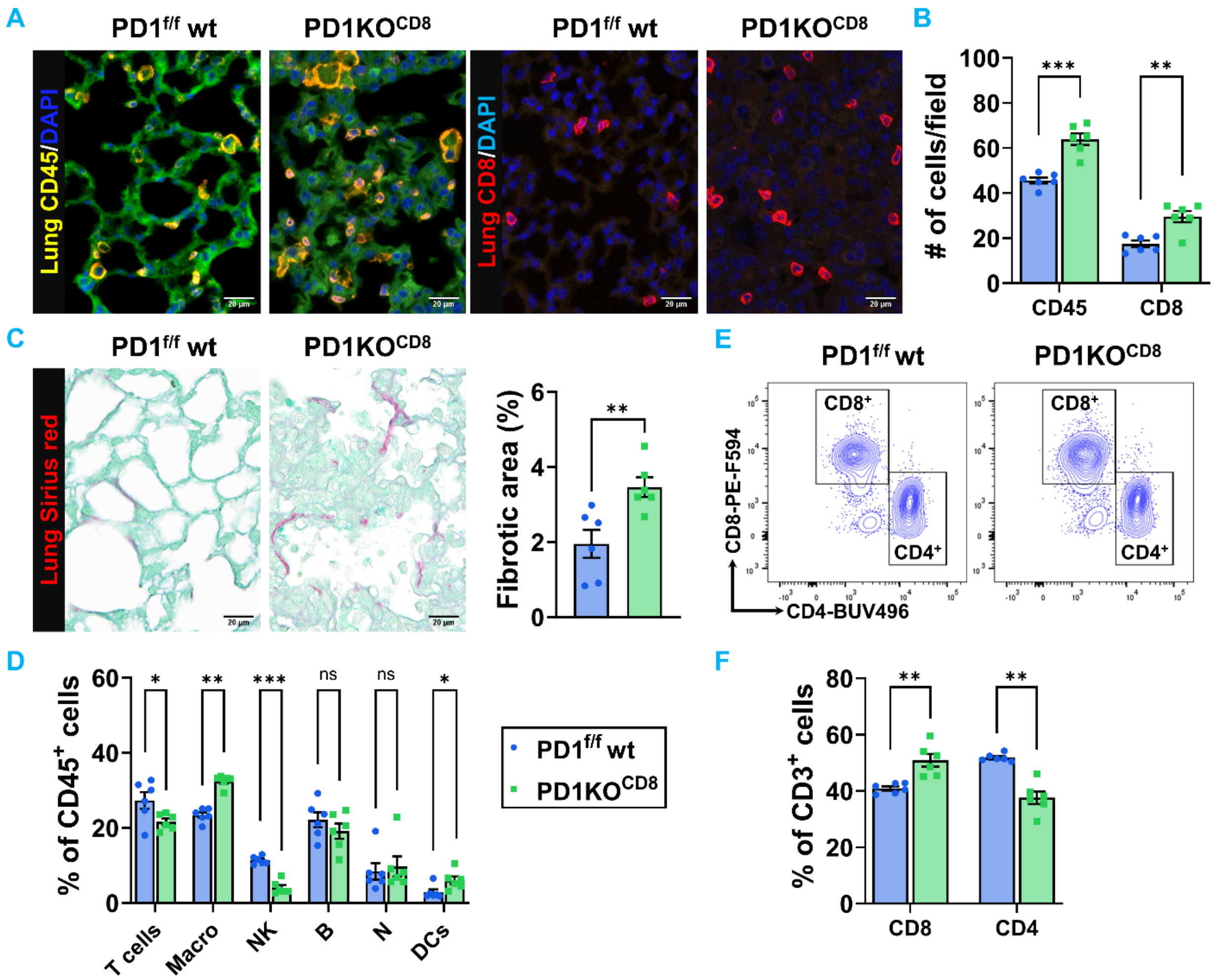


**Supplementary Figure S24. Immune cell infiltration in female lung tissues after TAC.**

**(A)** Representative immunofluorescence images of lung sections from female mice after TAC. Tissues were stained for CD45 (left panels, yellow) to identify total leukocytes and CD8 (right panels, red) to identify cytotoxic T cells. Nuclei were counterstained with DAPI (blue). **(B)** Quantitative analysis of infiltrating $CD45^+$ and $CD8^+$ cells per field in the lung corresponding to the images in panel **A**. **(C)** Representative images of lung sections stained with Sirius red to visualize baseline collagen deposition, alongside the corresponding quantification of the fibrotic area. **(D)** Flow cytometric quantification of major immune cell subsets—including T cells, macrophages (Macro), Natural Killer (NK) cells, B cells, neutrophils (N), and dendritic cells (DCs)—expressed as a percentage of total $CD45^+$ leukocytes isolated from lung tissue of female TAC-operated $PD1^{f/f}$ wt (blue) and $PD1KO^{CD8}$ (green) mice. **(E-F)** Representative flow cytometry contour plots identifying $CD8^+$ and $CD4^+$ T cell populations, with quantification expressed as the percentage of total $CD3^+$ T cells in lung tissue from female TAC-operated $PD1^{f/f}$ wt (blue) and $PD1KO^{CD8}$ (green) mice. Scale bar = 20 μm. Data are presented as mean ± SEM. Statistical significance was determined by unpaired two-tailed Student's t-test, except that Mann-Whitney test was used for macrophage, neutrophil, and DCs in Panel **(D)** and Welch's t-test was used for $CD8^+$ and $CD4^+$ T cells in Panel **(F)**. ns = not significant, *$p < 0.05$, **$p < 0.01$, ***$p < 0.001$. n = 6 mice per group.

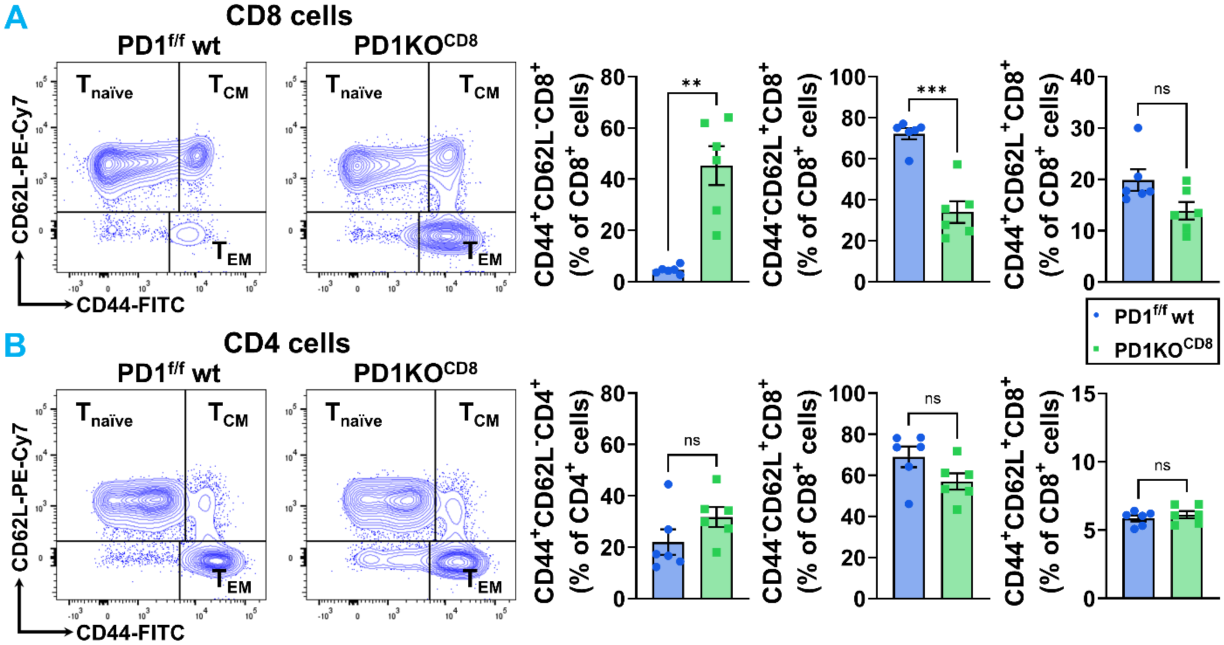


**Supplementary Figure S25. CD8-specific PD-1 deficiency significantly increases female pulmonary CD8$^+$ T cell activation.**

**(A)** Representative flow cytometry plots and quantification of naïve ($T_{naïve}$), effector memory ($T_{EM}$), and central memory ($T_{CM}$) subsets, gated by CD62L and CD44 expression among pulmonary CD8$^+$ T cells in the female TAC-operated PD1$^{f/f}$ wt (blue) and PD1KO$^{CD8}$ (green) mice. **(B)** Representative flow cytometry plots and quantification of naïve ($T_{naïve}$), effector memory ($T_{EM}$), and central memory ($T_{CM}$) subsets, among pulmonary CD4$^+$ T cells in the female TAC-operated PD1$^{f/f}$ wt (blue) and PD1KO$^{CD8}$ (green) mice. Data are presented as mean ± SEM. Statistical significance was determined by unpaired two-tailed Student's t-test, except that Mann-Whitney test was used for $T_{CM}$ in CD8$^+$ **(A)** and $T_{EM}$ and $T_{naïve}$ in CD4$^+$ **(B)** T cells. ns = not significant, $**p < 0.01$, $***p < 0.001$. n = 6 mice per group.

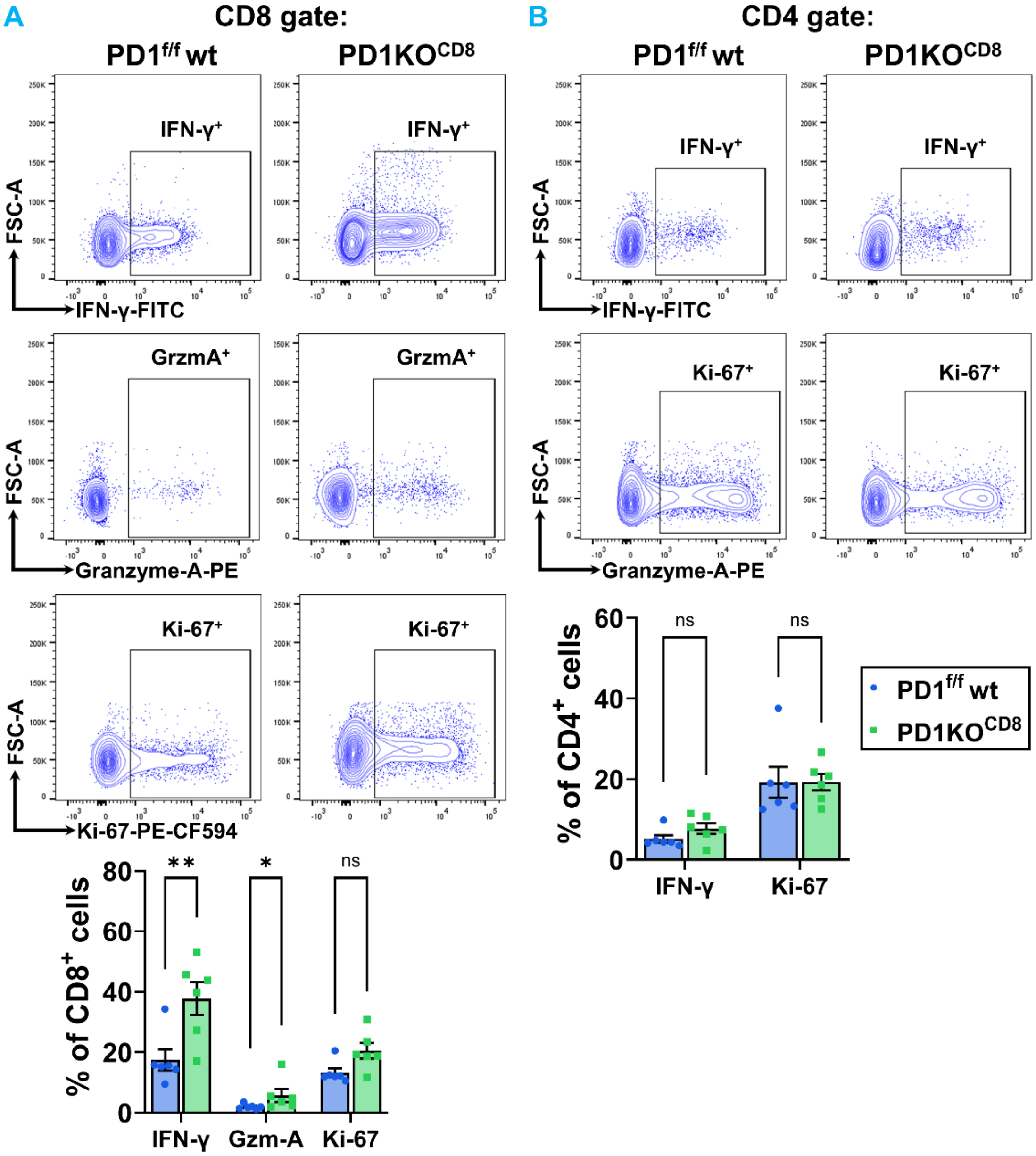


**Supplementary Figure S26. CD8-specific PD-1 deficiency significantly enhances intracellular cytokine production and proliferation in pulmonary CD8⁺ T cells after TAC.**

**(A)** Representative flow cytometry contour plots and analysis of IFN-$\gamma^+$, Granzyme $A^+$, and Ki-$67^+$ $CD8^+$ T cell subsets isolated from the lung of female $PD1^{f/f}$ wt (blue) and $PD1KO^{CD8}$ (green) mice. **(B)** Representative flow cytometry contour plots of IFN-$\gamma^+$ and Ki-$67^+$ $CD4^+$ T cell subsets isolated from the lung of female $PD1^{f/f}$ wt (blue) and $PD1KO^{CD8}$ (green) mice. Data are presented as mean ± SEM. Statistical significance was determined by unpaired two-tailed Student's t-test, except that Mann Whitney test was used for Granzyme $A^+CD8^+$ and Ki-$67^+CD8^+$ in panel **(A),** and Ki-$67^+CD4^+$ T cells in panel **(B)**. ns = not significant, *$p < 0.05$, **$p < 0.01$. n = 6 mice per group.

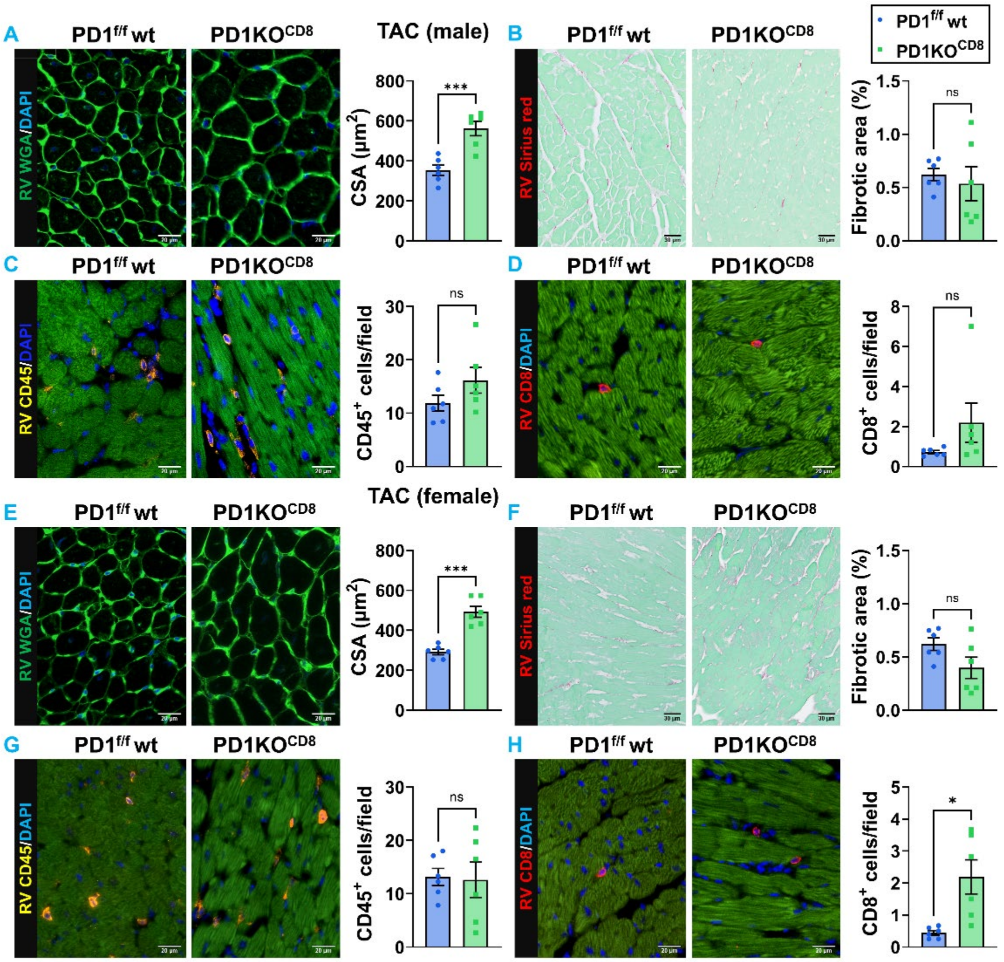


**Supplementary Figure S27. Histological analysis of right ventricular tissues in mice after TAC.**

**(A)** Representative images of right ventricular (RV) sections from male TAC-operated mice. Tissues were stained with WGA (green) to delineate cardiomyocyte membranes and assess cell size. Nuclei were counterstained with DAPI (blue). The accompanying bar graph quantifies the RV cardiomyocyte cross-sectional area (CSA). Scale bar = 20 μm. **(B)** Representative images of RV sections stained with Sirius red to visualize baseline collagen deposition, and the quantified fibrotic area. Scale bar = 30 μm. **(C-D)** Representative immunofluorescence images of RV sections from male TAC-operated mice and quantitative analysis of infiltrating $CD45^+$ and $CD8^+$ cells per mm² in the RV. Nuclei were counterstained with DAPI (blue). Scale bar = 20 μm. **(E)** Representative images of RV sections from female TAC-operated mice. Tissues were stained with WGA (green) to assess cell size. Nuclei were counterstained with DAPI (blue). The accompanying bar graph quantifies the RV cardiomyocyte CSA. Scale bar = 20 μm. **(F)** Representative images of RV sections stained with Sirius red to visualize baseline collagen deposition, alongside the corresponding quantification of the fibrotic area. Scale bar = 30 μm. **(G-H)** Representative images of RV sections from male TAC-operated mice and quantified $CD45^+$ and $CD8^+$ cells in the RV tissues. Nuclei were counterstained with DAPI (blue). Scale bar = 20 μm. Data represent TAC cohorts: $PD1^{f/f}$ wt TAC (blue bars) and $PD1KO^{CD8}$ TAC (green bars). Data are presented as mean ± SEM. Statistical significance was determined by unpaired two-tailed Student's t-test, except that Welch's t-test was used for panel **(B) and (H),** and Mann Whitney test was used for Panel **(D)**. n = 6 mice per group.

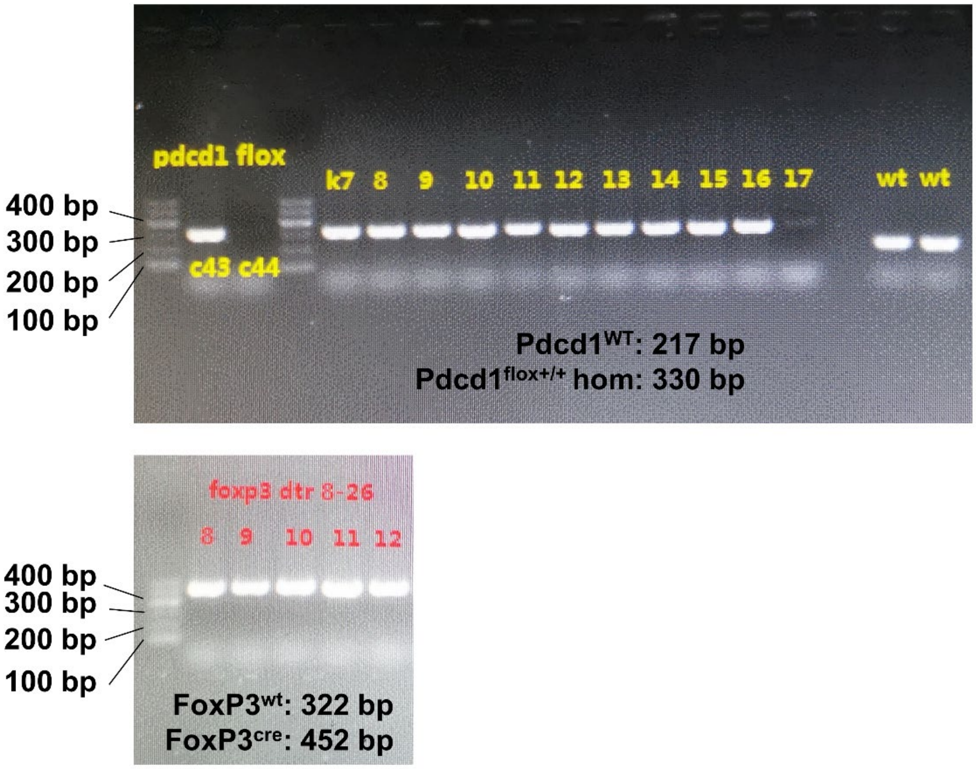


**Supplementary Figure S28. Represented PCR for genotyping of the Treg cell-specific PD1 knockout and PD1$^{f/f}$ wt strains.**

The top gel demonstrates genotyping for the *Pdcd1* allele, distinguishing the wild-type (WT) band at 217 bp from the homozygous floxed band at 330 bp. The bottom gel demonstrates genotyping for the FoxP3-Cre transgene, identifying the hemizygous FoxP3-Cre genotype by the presence of a 452 bp band.

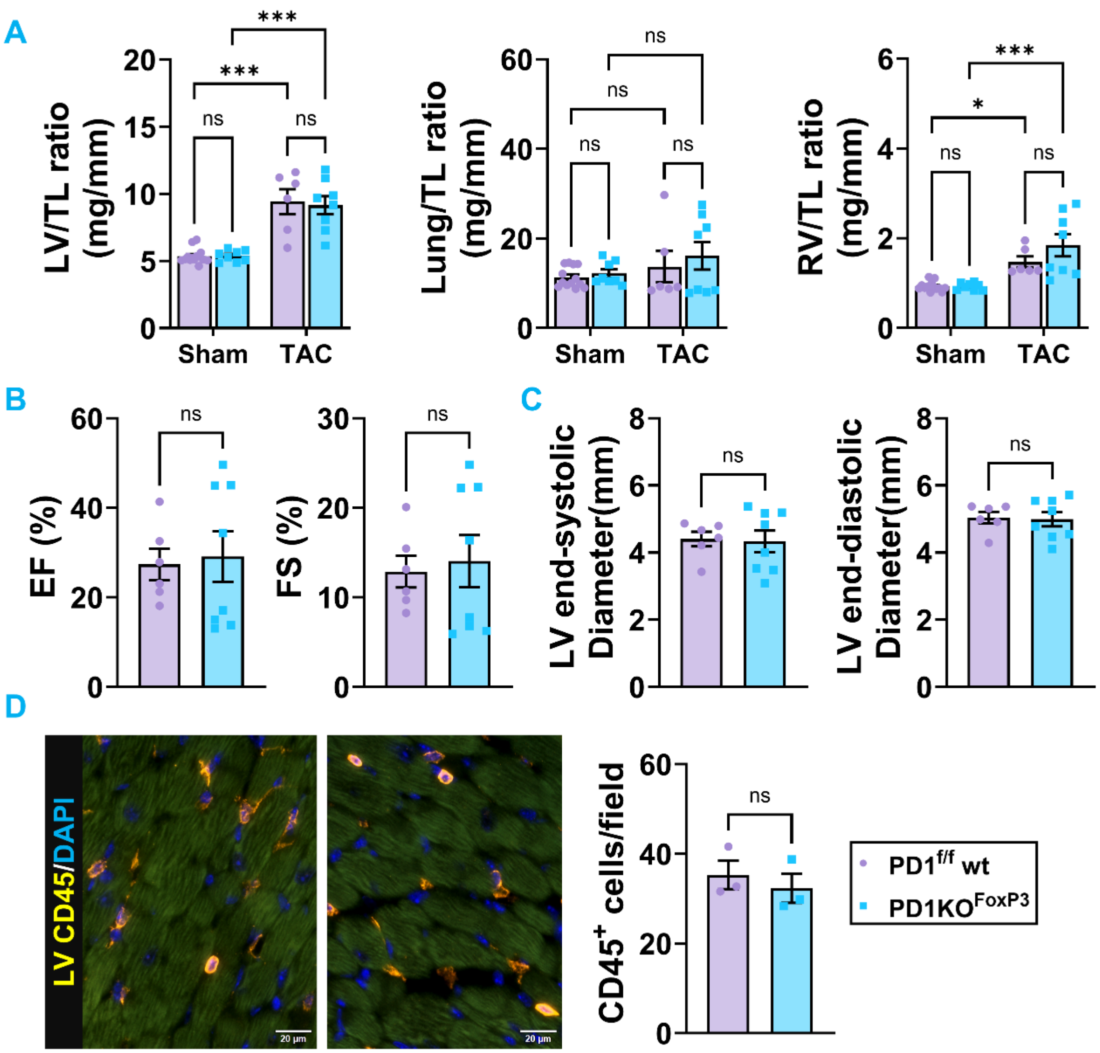


**Supplementary Figure S29. Treg-specific deletion of PD-1 does not rescue cardiac function or inflammation after TAC.**
**(A)** Assessment of cardiac hypertrophy and pulmonary congestion via heart weight-to-tibial length (HW/TL), lung weight-to-tibial length (LungW/TL), and right ventricular weight-to-tibial length (RV/TL) ratios. **(B)** Quantitative analysis of systolic function, including ejection fraction (EF%) and fractional shortening (FS%), in $PD1^{f/f}$ wt (lavender) and $PD1KO^{FoxP3}$ (cyan) mice, evaluated at 6 weeks post-TAC. **(C)** Quantitative analysis by echocardiography of LV dimensions at end-systole and end-diastole. **(D)** Representative immunofluorescence images of LV sections stained for total leukocytes (CD45, orange; left panels). Nuclei were counterstained with DAPI (blue). The accompanying bar graph quantifies the infiltrating $CD45^+$ cells per field in the myocardium. Scale bar = 20 µm. Data are presented as mean ± SEM. Statistical significance was determined by unpaired two-tailed Student's t-test for panels (A) and (B), except that Mann-Whitney test was used for panel **(D)**. Two-way ANOVA with Tukey's multiple comparisons test was used for panel **(C)**. ns = not significant, *p < 0.05, ***p < 0.001.  n = 3-13 mice per group.

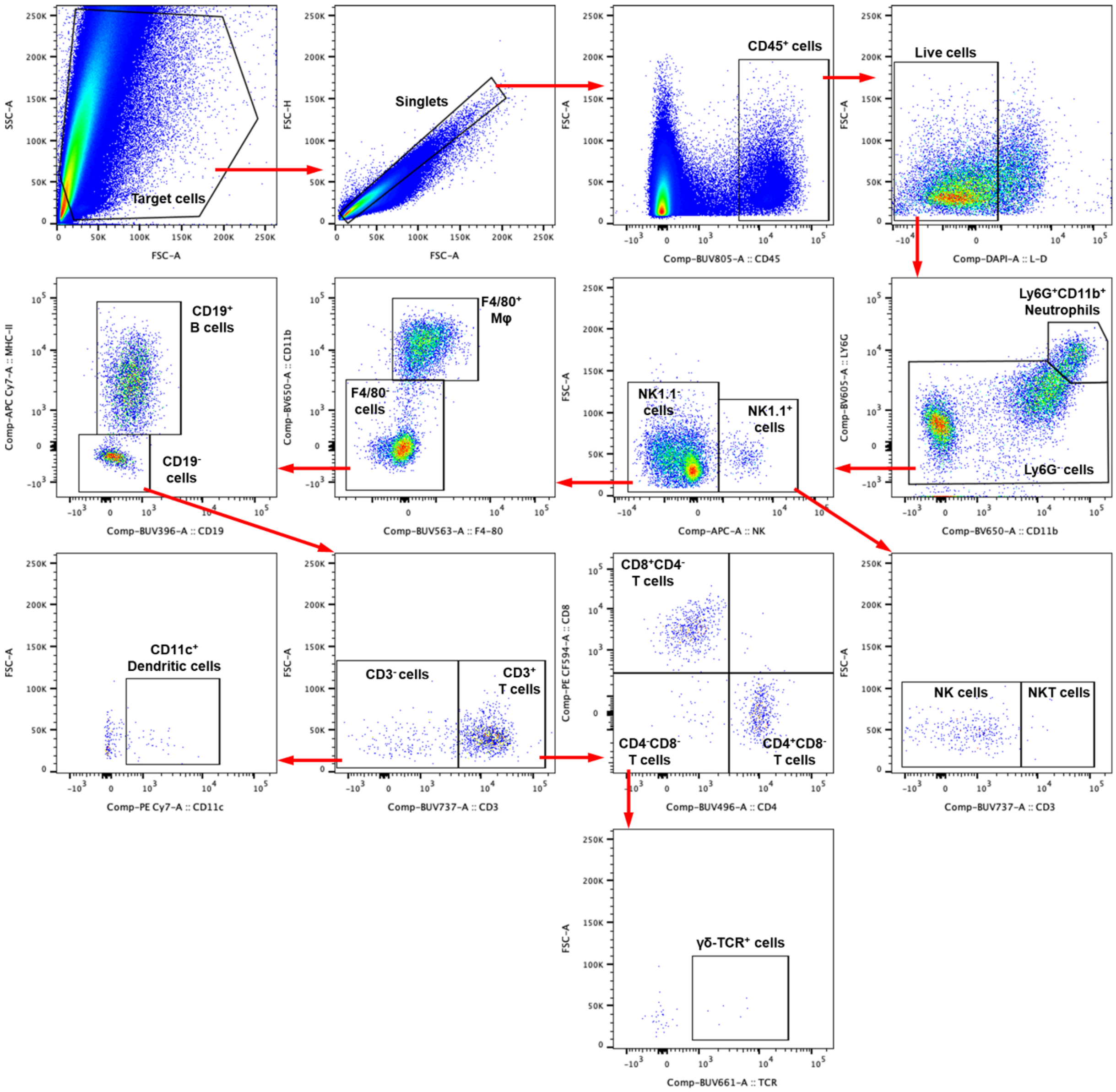


**Supplementary Figure S30. Gating strategy for cardiac and lymph node immune cell profile in global PD1KO mice subjected to TAC for 7 days and treated with IgG isotype control or anti-CD8α antibodies.**

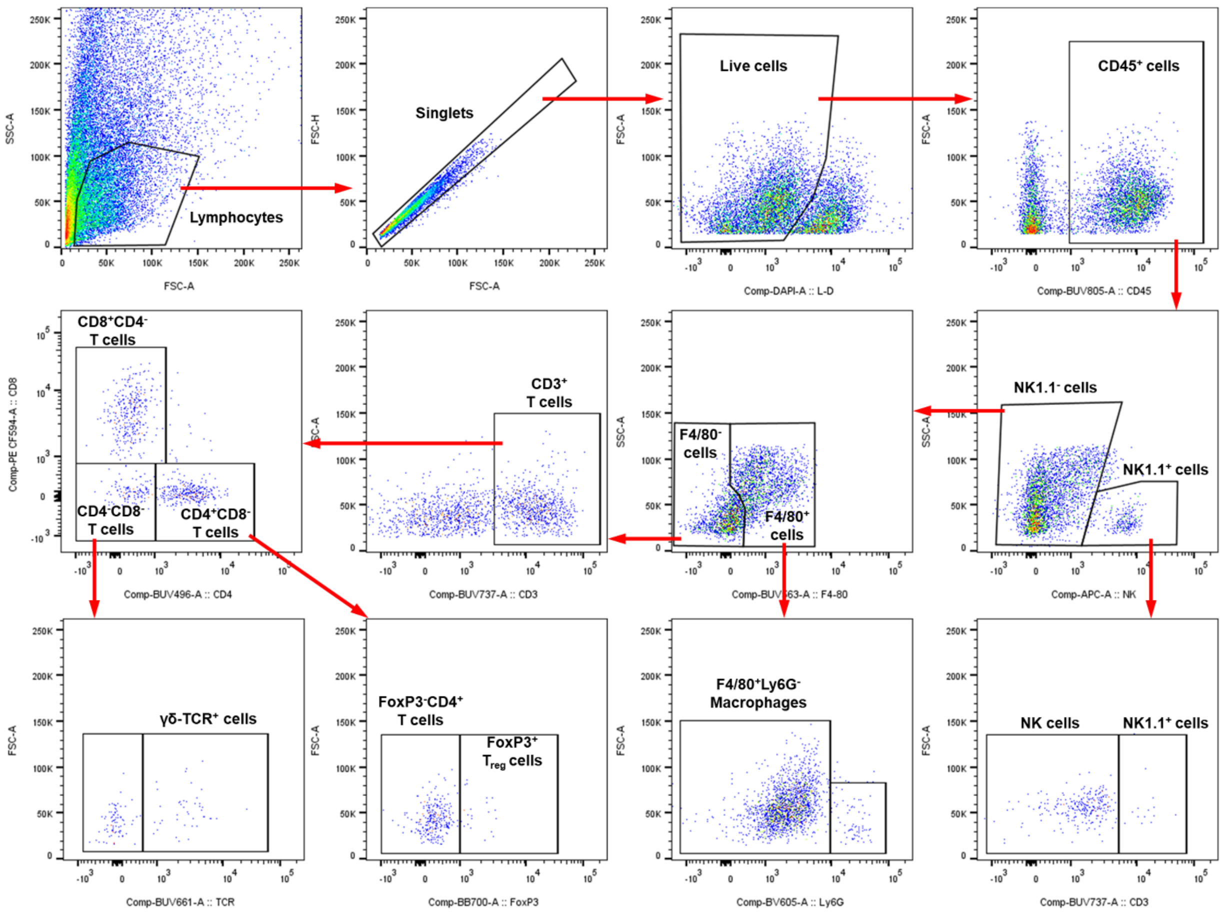


**Supplementary Figure S31. Gating strategy for cytokine production in $CD8^+$ and $CD4^+$ T cells of cardiac tissue or PBMC from $PD1^{f/f}$ wt and $PD1KO^{CD8}$ mice under sham or TAC conditions.**

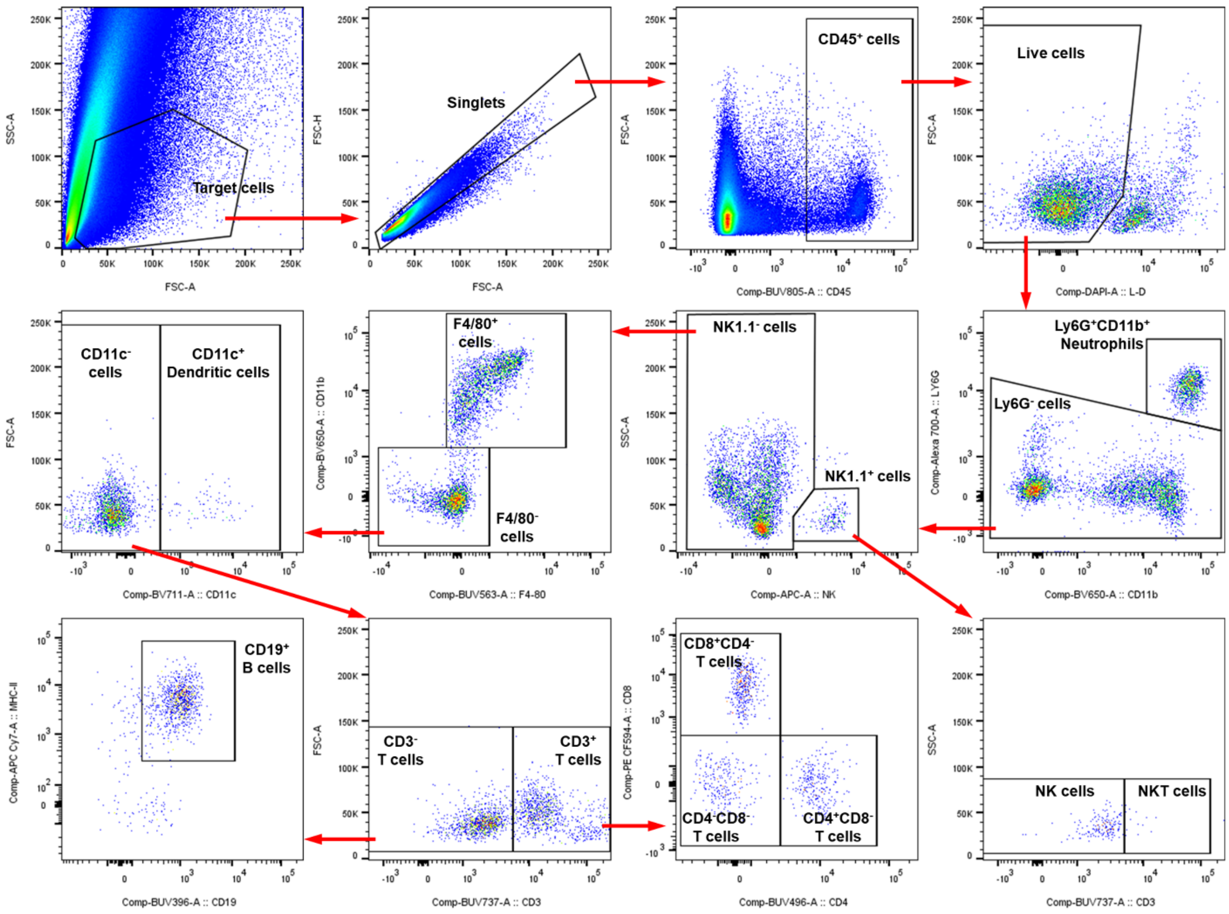


**Supplementary Figure S32. Gating strategy for immune cell profile of cardiac tissue or PBMC from PD1$^{f/f}$ wt and PD1KO$^{CD8}$ mice under sham or TAC conditions.**

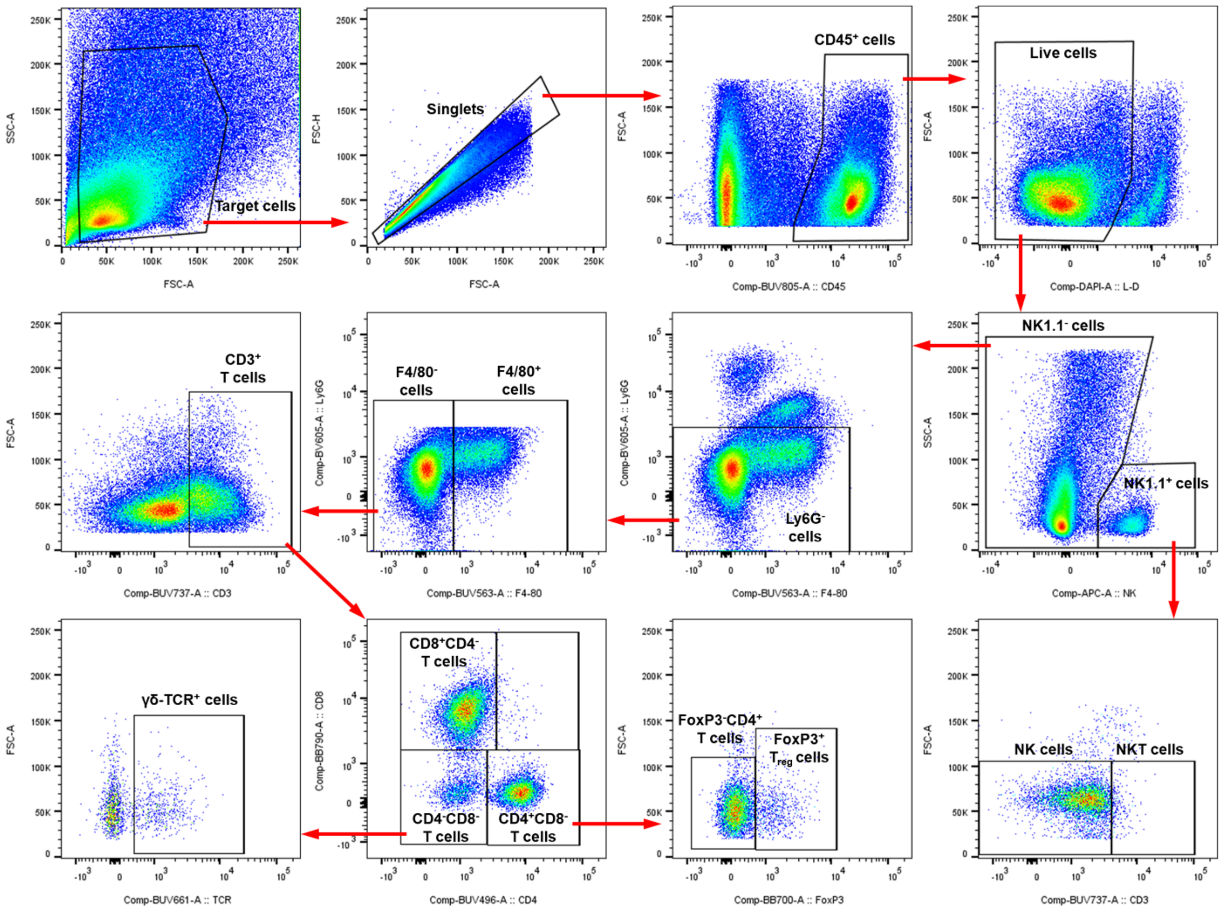


**Supplementary Figure S33. Gating strategy for cytokine production in CD8$^{+}$ and CD4$^{+}$ T cells of lung tissue from PD1$^{f/f}$ wt and PD1KO$^{CD8}$ mice under sham or TAC conditions.**

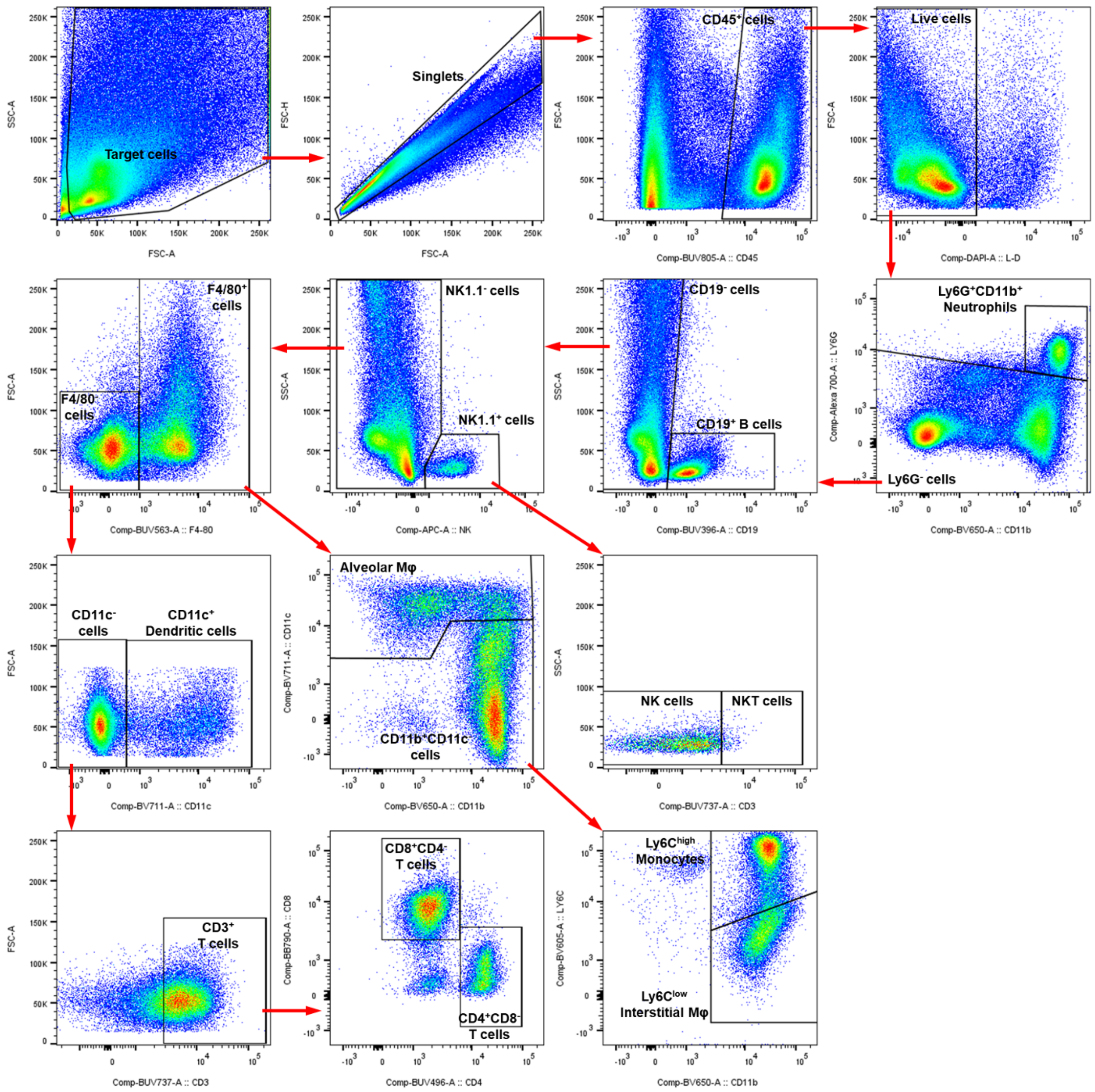


**Supplementary Figure S34. Gating strategy for immune cell profile of lung tissue from PD1$^{f/f}$ wt and PD1KO$^{CD8}$ mice under sham or TAC conditions.**

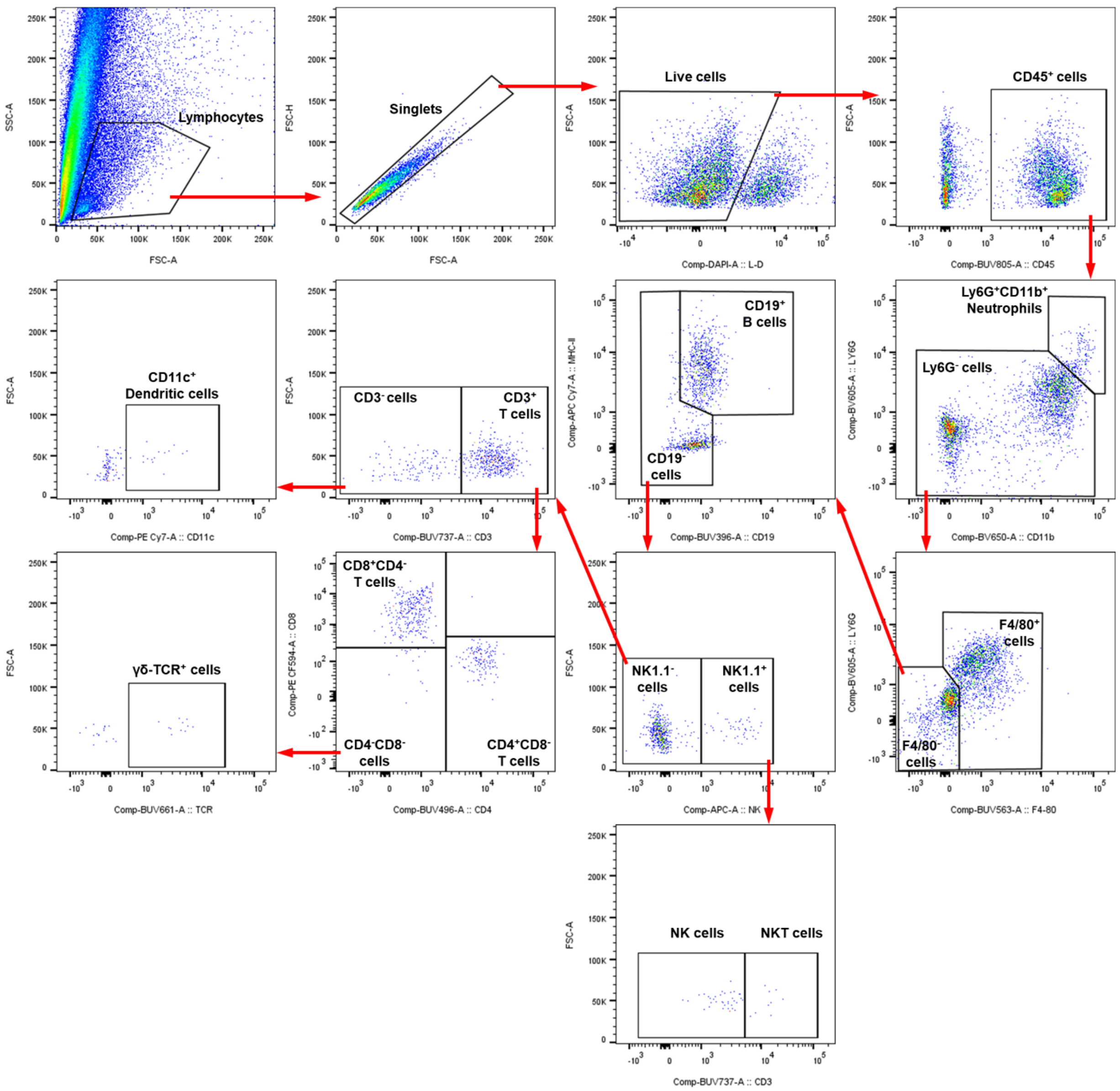


1

2 **Supplementary Figure S35. Gating strategy for immune cell profile of cardiac tissue from**
3 **PD1$^{f/f}$ wt and PD1KO$^{CD8}$ mice subjected to TAC for 7 days and treated with IgG isotype**
4 **control or anti-CXCR3 antibodies.**

**Supplementary Table S1. Echocardiographic data in experimental groups.**

| Parameters | ♂ PD1KO + TAC 1W + IgG Ab (n=12) | ♂ PD1KO + TAC 1W + anti-CD8α Ab (n=9) |
|---|---|---|
| **HR (bpm)** | 561±12 | 567±7 $^{\text{ns}}$ |
| **LVESD (mm)** | 2.86±0.11 | 2.36±0.09 ** |
| **LVEDD (mm)** | 4.00±0.10 | 3.62±0.09 * |
| **LVESV (µL)** | 32.02±2.97 | 19.79±1.88 ** |
| **LVEDV (µL)** | 70.63±4.12 | 55.74±3.26 * |
| **SV (µL)** | 38.61±1.96 | 35.95±1.46 $^{\text{ns}}$ |
| **CO (mL/min)** | 21.80±1.44 | 20.32±0.68 $^{\text{ns}}$ |
| **LVAW;s (mm)** | 1.55±0.05 | 1.50±0.03 $^{\text{ns}}$ |
| **LVAW;d (mm)** | 1.09±0.04 | 1.00±0.02 * |
| **LVPW;s (mm)** | 1.32±0.05 | 1.21±0.03 $^{\text{ns}}$ |
| **LVPW;d (mm)** | 0.90±0.04 | 0.81±0.02 $^{\text{ns}}$ |

Data are shown as mean ± SEM. HR, heart rate; LVESD, left ventricular end-systolic diameter; LVEDD, left ventricular end-diastolic diameter; LVESV, left ventricular end-systolic volume; LVEDV, left ventricular end-diastolic volume; SV, stroke volume; CO, cardiac output; LVAW, left ventricular anterior wall; LVPW, left ventricular posterior wall. Data are presented as mean ± SEM. Statistical significance was determined by unpaired two-tailed Student's t-test, Welch's t-test, or Mann-Whitney test. n = 9-12 mice per group. *p<0.05 vs. WT+TAC, **p<0.01 vs. WT+TAC, ns, not significant.

**Supplementary Table S2. Tissue weight and ratios of mice treated with anti-CD8α Ab.**

| Parameters | ♂ PD1KO + TAC 1W + IgG isotype Ab (n=12) | ♂ PD1KO + TAC 1W + anti-CD8α Ab (n=9) |
|---|---|---|
| BW (g) | 27.5±1.8 | 26.0±1.5 ns |
| LV (mg) | 139.9±6.1 | 97.3±4.7 *** |
| RV (mg) | 21.9±1.1 | 19.5±0.8 ns |
| LA (mg) | 6.1±0.4 | 3.6±0.1 *** |
| RA (mg) | 4.5±0.4 | 3.1±0.1 ** |
| LV/RV | 6.5±0.3 | 5.0±0.1 *** |
| LA/RA | 1.4±0.1 | 1.2±0.1 * |
| Ventricles (mg) | 161.7±6.8 | 116.8±5.4 *** |
| Heart (mg) | 172.3±7.4 | 123.5±5.5 *** |
| Lung (mg) | 233.1±20.6 | 143.9±4.9 ** |
| Spleen (mg) | 96.6±5.8 | 104.0±5.2 ns |
| TL (mm) | 17.18±0.22 | 16.76±0.22 ns |
| LV/BW | 5.20±0.23 | 3.76±0.10 *** |
| RV/BW | 0.81±0.04 | 0.76±0.03 ns |
| LA/BW | 0.23±0.02 | 0.14±0.01 *** |
| Ventricles/BW | 6.01±0.26 | 4.52±0.13 *** |
| Heart/BW | 6.40±0.28 | 4.78±0.14 *** |
| Lung/BW | 8.66±0.73 | 5.66±0.37 ** |
| Spleen/BW | 3.56±0.18 | 4.12±0.34 ns |
| LV/TL | 8.12±0.29 | 5.79±0.22 *** |
| LA/TL | 0.36±0.02 | 0.22±0.01 *** |
| RV/TL | 1.27±0.05 | 1.16±0.04 ns |
| Ventricles/TL | 9.39±0.32 | 6.95±0.25 *** |
| Heart/TL | 10.00±0.34 | 7.35±0.26 *** |
| Lung/TL | 13.52±1.13 | 8.59±0.31 ** |
| Spleen/TL | 5.60±0.30 | 6.22±0.34 ns |

Data are shown as mean ± SEM. BW, body weight; LV, left ventricle; RV, right ventricle; LA, left atrium; RA, right atrium; TL, tibial length. Data are presented as mean ± SEM. Statistical significance was determined by unpaired two-tailed Student's t-test, Welch's t-test, or Mann-Whitney test. n = 9-12 mice per group. * p<0.05, ** p<0.01, ***p<0.001. ns, not significant.

**Supplementary Table S3. Echocardiographic data in male experimental groups.**

| Parameters | ♂ PD1$^{f/f}$ wt sham (n=6) | ♂ PD1KO$^{CD8}$ sham (n=5) | ♂ PD1$^{f/f}$ wt TAC (n=24) | ♂ PD1KO$^{CD8}$ TAC (n=24) |
|---|---|---|---|---|
| **HR (bpm)** | 598±14 | 620±14 | 577±10 | 569±8 $^{ns}$ |
| **LVESD (mm)** | 2.59±0.18 | 2.26±0.09 | 3.78±0.20 * | 4.37±0.16 *** |
| **LVEDD (mm)** | 4.07±0.17 | 3.70±0.08 | 4.66±0.15 | 5.02±0.12 *** |
| **LVESV (µL)** | 25.37±4.74 | 17.56±1.79 | 66.98±7.31 * | 89.72±7.19 *** |
| **LVEDV (µL)** | 74.13±7.88 | 58.51±3.03 | 103.81±7.30 | 121.59±6.49 *** |
| **SV (µL)** | 48.77±3.61 | 40.95±1.38 | 36.83±1.25 ** | 31.87±2.08 |
| **CO (mL/min)** | 29.06±2.01 | 25.44±1.26 | 21.29±0.87 ** | 18.28±1.28 * |
| **LVAW;s (mm)** | 1.11±0.05 | 1.15±0.02 | 1.16±0.04 | 1.14±0.05 $^{ns}$ |
| **LVAW;d (mm)** | 0.68±0.05 | 0.70±0.03 | 0.88±0.03 * | 0.90±0.03 * |
| **LVPW;s (mm)** | 0.90±0.05 | 0.97±0.03 | 0.91±0.03 | 0.89±0.03 $^{ns}$ |
| **LVPW;d (mm)** | 0.57±0.03 | 0.60±0.04 | 0.74±0.02 ** | 0.79±0.02 ** |

Data are shown as mean ± SEM. HR, heart rate; LVESD, left ventricular end-systolic diameter; LVEDD, left ventricular end-diastolic diameter; LVESV, left ventricular end-systolic volume; LVEDV, left ventricular end-diastolic volume; SV, stroke volume; CO, cardiac output; LVAW, left ventricular anterior wall; LVPW, left ventricular posterior wall. Data are presented as mean ± SEM. Statistical significance was determined by *two-way ANOVA with Tukey's multiple comparisons test.* n = 5-24 mice per group. */† p<0.05, **/†† p<0.01, ***/††† p<0.001. * vs. corresponding sham, † vs. PD1$^{f/f}$ wt+TAC. ns, not significant.

**Supplementary Table S4. Tissue weight and ratios of male PD1$^{f/f}$ wt and PD1KO$^{CD8}$ mice.**

| Parameters | ♂ PD1$^{f/f}$ wt sham (n=6) | ♂ PD1KO$^{CD8}$ sham (n=5) | ♂ PD1$^{f/f}$ wt TAC (n=23) | ♂ PD1KO$^{CD8}$ TAC (n=24) |
|---|---|---|---|---|
| **BW (g)** | 27.5±1.1 | 26.8±0.7 | 26.0±0.8 | 25.9±0.8 $^{ns}$ |
| **LV (mg)** | 91.2±2.9 | 81.9±2.5 | 150.3±6.8 *** | 177.1±5.1 *** |
| **RV (mg)** | 19.2±1.4 | 19.4±2.0 | 25.5±1.8 | 31.4±1.9 * |
| **LA (mg)** | 2.3±0.2 | 2.8±0.4 | 9.4±1.9 | 18.1±2.4 * † |
| **RA (mg)** | 2.6±0.4 | 2.5±0.3 | 3.0±0.2 | 4.1±0.3 * †† |
| **LV/RV** | 4.9±0.4 | 4.4±0.5 | 6.2±0.3 | 6.0±0.3 $^{ns}$ |
| **LA/RA** | 0.9±0.1 | 1.11±0.2 | 3.1±0.7 | 4.6±0.7 $^{ns}$ |
| **Ventricles (mg)** | 110.4±3.7 | 101.3±2.4 | 175.8±8.3 *** | 208.5±6.6 *** †† |
| **Heart (mg)** | 115.3±4.0 | 106.5±2.7 | 188.1±10.0 ** | 230.8±8.4 *** †† |
| **Lung (mg)** | 134.7±4.8 | 128.9±5.1 | 267.8±33.1 | 388.9±31.3 ** † |
| **Spleen (mg)** | 76.3±9.6 | 67.4±3.5 | 85.9±3.7 | 90.9±3.6 $^{*}$ |
| **TL (mm)** | 17.44±0.11 | 17.34±0.04 | 17.29±0.09 | 17.41±0.07 $^{ns}$ |
| **LV/BW** | 3.34±0.16 | 3.06±0.11 | 5.93±0.37 ** | 7.04±0.33 *** |
| **RV/BW** | 0.34±0.17 | 0.10±0.02 | 0.88±0.13 | 1.07±0.14 * |
| **LA/BW** | 0.45±0.11 | 0.73±0.09 | 0.54±0.08 | 0.97±0.10 †† |
| **Ventricles/BW** | 4.05±0.21 | 3.79±0.15 | 6.95±0.46 * | 8.31±0.42 *** |
| **Heart/BW** | 4.23±0.23 | 3.99±0.17 | 7.46±0.54 * | 9.24±0.53 *** |
| **Lung/BW** | 4.94±0.28 | 4.83±0.25 | 11.06±1.65 | 15.83±1.52 * |
| **Spleen/BW** | 2.84±0.45 | 2.52±015 | 3.37±0.19 | 3.59±0.17 $^{ns}$ |
| **LV/TL** | 5.23±0.17 | 4.72±0.14 | 8.69±0.39 *** | 10.18±0.29 *** †† |
| **LA/TL** | 0.13±0.01 | 0.16±0.02 | 0.55±0.11 | 1.05±0.14 * † |
| **RV/TL** | 1.10±0.08 | 1.12±0.11 | 1.48±0.11 | 1.81±0.11 * |
| **Ventricles/TL** | 6.33±0.23 | 5.84±0.13 | 10.17±0.49 *** | 11.99±0.38 *** † |
| **Heart/TL** | 6.61±0.25 | 6.14±0.15 | 10.89±0.58 ** | 13.27±0.49 *** †† |
| **Lung/TL** | 7.73±0.30 | 7.43±0.29 | 15.56±1.96 | 22.40±1.83 ** † |
| **Spleen/TL** | 4.39±0.58 | 3.88±0.19 | 4.97±0.22 | 5.22±0.21 $^{ns}$ |

Data are shown as mean ± SEM. BW, body weight; LV, left ventricle; RV, right ventricle; LA, left atrium; RA, right atrium; TL, tibial length. Data are presented as mean ± SEM. Statistical significance was determined by *two-way ANOVA with Tukey's multiple comparisons test.* n = 5-24 mice per group. */† p<0.05, **/†† p<0.01, ***/††† p<0.001. * vs. corresponding sham, † vs. PD1$^{f/f}$ wt+TAC. ns, not significant.

**Supplementary Table S5. Echocardiographic data in female experimental groups.**

| Parameters | ♀ PD1$^{f/f}$ wt sham (n=6) | ♀ PD1KO$^{CD8}$ sham (n=7) | ♀ PD1$^{f/f}$ wt TAC (n=31) | ♀ PD1KO$^{CD8}$ TAC (n=29) |
|---|---|---|---|---|
| **HR (bpm)** | 636±13 | 604±9 | 576±9 * | 568±8 $^{ns}$ |
| **LVESD (mm)** | 2.35±0.05 | 2.30±0.05 | 3.48±0.16 ** | 3.87±0.13 *** |
| **LVEDD (mm)** | 3.73±0.06 | 3.73±0.07 | 4.38±0.11 * | 4.56±0.09 ** |
| **LVESV (µL)** | 19.18±1.07 | 18.12±0.95 | 55.28±5.81 * | 67.95±4.95 *** |
| **LVEDV (µL)** | 59.54±2.34 | 59.36±2.82 | 88.97±5.13 * | 96.86±4.13 ** |
| **SV (µL)** | 40.36±1.73 | 41.23±1.99 | 33.69±1.34 | 28.91±1.58 ** |
| **CO (mL/min)** | 25.68±1.22 | 24.94±1.38 | 19.40±0.82 * | 16.56±1.00 *** |
| **LVAW;s (mm)** | 1.14±0.04 | 1.11±0.05 | 1.13±0.04 | 1.12±0.04 $^{ns}$ |
| **LVAW;d (mm)** | 0.70±0.02 | 0.66±0.02 | 0.84±0.02 * | 0.86±0.02 *** |
| **LVPW;s (mm)** | 0.84±0.03 | 0.89±0.03 | 0.89±0.02 | 0.87±0.02 $^{ns}$ |
| **LVPW;d (mm)** | 0.55±0.01 | 0.59±0.02 | 0.72±0.02 *** | 0.76±0.02 *** |

Data are shown as mean ± SEM. HR, heart rate; LVESD, left ventricular end-systolic diameter; LVEDD, left ventricular end-diastolic diameter; LVESV, left ventricular end-systolic volume; LVEDV, left ventricular end-diastolic volume; SV, stroke volume; CO, cardiac output; LVAW, left ventricular anterior wall; LVPW, left ventricular posterior wall. Data are presented as mean ± SEM. Statistical significance was determined by two-way ANOVA with Tukey's multiple comparisons test. n = 6-31 mice per group. */† p<0.05, **/†† p<0.01, ***/††† p<0.001. * vs. corresponding sham, † vs. PD1$^{f/f}$ wt+TAC. ns, not significant.

**Supplementary Table S6. Tissue weight and ratios of female PD1$^{f/f}$ wt and PD1KO$^{CD8}$ mice.**

| Parameters | ♂ PD1$^{f/f}$ wt sham (n=6) | ♂ PD1KO$^{CD8}$ sham (n=7) | ♂ PD1$^{f/f}$ wt TAC (n=22) | ♂ PD1KO$^{CD8}$ TAC (n=18) |
|---|---|---|---|---|
| **BW (g)** | 21.7±1.0 | 23.3±0.8 | 22.5±0.4 | 20.9±0.5 * |
| **LV (mg)** | 75.6±1.9 | 76.2±3.0 | 124.3±5.4 *** | 149.2±3.4 *** ††† |
| **RV (mg)** | 16.3±0.4 | 15.1±0.5 | 19.3±1.1 | 26.0±1.8 *** †† |
| **LA (mg)** | 1.9±0.1 | 1.8±0.1 | 4.7±0.7 | 13.3±2.3 *** ††† |
| **RA (mg)** | 2.4±0.3 | 2.1±0.2 | 2.8±0.2 | 3.1±0.2 * |
| **LV/RV** | 4.6±0.2 | 5.0±0.1 | 6.6±0.2 ** | 6.1±0.3 |
| **LA/RA** | 0.9±0.1 | 0.9±0.1 | 1.7±0.2 | 4.1±0.6 *** ††† |
| **Ventricles (mg)** | 91.9±1.9 | 91.3±3.4 | 143.5±6.3 *** | 175.3±5.0 *** ††† |
| **Heart (mg)** | 96.1±2.2 | 95.2±3.4 | 151.0±6.9 *** | 191.7±6.8 *** ††† |
| **Lung (mg)** | 168.8±29.1 | 140.0±5.3 | 196.7±21.9 | 339.1±37.3 ** †† |
| **Spleen (mg)** | 83.7±4.4 | 88.0±2.0 | 98.0±2.8 | 111.1±7.5 $^{ns}$ |
| **TL (mm)** | 16.72±0.19 | 17.14±0.13 | 17.06±0.09 | 17.00±0.10 $^{ns}$ |
| **LV/BW** | 3.50±0.10 | 3.27±0.05 | 5.56±0.28 *** | 7.19±0.22 *** ††† |
| **RV/BW** | 0.76±0.04 | 0.65±0.02 | 0.87±0.06 | 1.27±0.10 *** †† |
| **LA/BW** | 0.09±0.00 | 0.08±0.00 | 0.21±0.04 | 0.66±0.13 ** ††† |
| **Ventricles/BW** | 4.26±0.14 | 3.92±0.06 | 6.43±0.33 ** | 8.46±0.32 *** ††† |
| **Heart/BW** | 4.45±0.13 | 4.08±0.06 | 6.77±0.36 * | 9.27±0.44 *** ††† |
| **Lung/BW** | 7.67±1.14 | 6.00±0.09 | 8.95±1.12 | 16.48±1.93 ** †† |
| **Spleen/BW** | 3.86±0.15 | 3.17±0.54 | 4.37±0.12 | 5.34±0.35 ** † |
| **LV/TL** | 4.52±0.08 | 4.44±0.14 | 7.29±0.32 *** | 8.78±0.20 *** ††† |
| **LA/TL** | 0.11±0.00 | 0.11±0.01 | 0.27±0.04 | 0.78±0.14 *** ††† |
| **RV/TL** | 0.98±0.03 | 0.88±0.02 | 1.13±0.07 | 1.53±0.11 *** †† |
| **Ventricles/TL** | 5.50±0.08 | 5.32±0.16 | 8.42±0.37 *** | 10.32±0.30 *** ††† |
| **Heart/TL** | 5.75±0.10 | 5.55±0.16 | 8.86±0.41 *** | 11.28±0.41 *** ††† |
| **Lung/TL** | 10.06±1.70 | 8.16±0.25 | 11.55±1.29 | 19.97±2.22 **†† |
| **Spleen/TL** | 4.99±0.22 | 4.37±0.74 | 5.74±0.16 | 6.54±0.45 $^{ns}$ |

Data are shown as mean ± SEM. BW, body weight; LV, left ventricle; RV, right ventricle; LA, left atrium; RA, right atrium; TL, tibial length. Data are presented as mean ± SEM. Statistical significance was determined by two-way ANOVA with Tukey's multiple comparisons test. n = 6-22 mice per group. */† p<0.05, **/†† p<0.01, ***/††† p<0.001. * vs. corresponding sham, † vs. PD1$^{f/f}$ wt+TAC. ns, not significant.

**Supplementary Table S7. Echocardiographic data in experimental groups.**

| Parameters | ♂ $PD1^{f/f}$ wt + TAC 1W + IgG isotype Ab (n=8) | ♂ $PD1KO^{CD8}$ + TAC 1W + IgG isotype Ab (n=10) | ♂ $PD1KO^{CD8}$ + TAC 1W + anti-CXCR3 Ab (n=10) |
|---|---|---|---|
| HR (bpm) | 582±13 | 562±12 | 560±10 $^{ns}$ |
| LVESD (mm) | 2.51±0.13 | 3.44±0.10 ** | 2.75±0.18 †† |
| LVEDD (mm) | 3.72±0.11 | 4.36±0.08 *** | 3.98±0.10 † |
| LVESV (µL) | 23.34±2.99 | 49.56±3.44 ** | 30.00±5.60 †† |
| LVEDV (µL) | 59.66±4.15 | 86.27±3.57 *** | 69.73±4.63 † |
| SV (µL) | 36.32±2.50 | 36.71±1.89 | 39.73±1.44 $^{ns}$ |
| CO (mL/min) | 21.09±1.44 | 20.58±1.01 | 22.23±0.87 $^{ns}$ |
| LVAW;s (mm) | 84.30±3.95 | 1.41±0.05 | 1.21±0.03 †† |
| LVAW;d (mm) | 1.34±0.03 | 1.01±0.05 | 0.85±0.02 † |
| LVPW;s (mm) | 0.94±0.03 | 0.89±0.02 | 0.91±0.04 $^{ns}$ |
| LVPW;d (mm) | 0.97±0.03 | 0.78±0.03 * | 0.61±0.02 ††† |

Data are shown as mean ± SEM. HR, heart rate; LVESD, left ventricular end-systolic diameter; LVEDD, left ventricular end-diastolic diameter; LVESV, left ventricular end-systolic volume; LVEDV, left ventricular end-diastolic volume; SV, stroke volume; CO, cardiac output; LVAW, left ventricular anterior wall; LVPW, left ventricular posterior wall. Data are presented as mean ± SEM. Statistical significance was determined by one-way ANOVA with Tukey's multiple comparisons test or Kruskal-Wallis test with Dunn's multiple comparisons test. n = 8-10 mice per group. */† $p<0.05$, **/†† $p<0.01$, ***/††† $p<0.001$. * vs. $PD1^{f/f}$ wt+TAC+IgG isotype, † vs. $PD1KO^{CD8}$+TAC+IgG isotype. ns, not significant.

**Supplementary Table S8. Tissue weight and ratios of mice treated with anti-CXCR3 Ab.**

| Parameters | ♂ PD1$^{f/f}$ wt + TAC 1W + IgG isotype Ab (n=8) | ♂ PD1KO$^{CD8}$ + TAC 1W + IgG isotype Ab (n=10) | ♂ PD1KO$^{CD8}$ + TAC 1W + anti-CXCR3 Ab (n=10) |
|---|---|---|---|
| **BW (g)** | 28.8±0.9 | 27.7±1.1 | 27.6±1.2 $^{ns}$ |
| **LV (mg)** | 123.8±4.9 | 151.9±7.0 ** | 109.3±3.9 ††† |
| **RV (mg)** | 18.8±0.7 | 25.7±1.7 ** | 20.5±1.2 † |
| **LA (mg)** | 3.6±0.4 | 7.4±1.4 | 5.7±2.6 $^{ns}$ |
| **RA (mg)** | 3.4±0.2 | 3.9±0.3 | 3.4±0.2 $^{ns}$ |
| **LV/RV** | 6.6±0.4 | 6.1±0.4 | 5.4±0.2 $^{ns}$ |
| **LA/RA** | 1.1±0.1 | 2.0±0.5 | 1.6±0.6 $^{ns}$ |
| **Ventricles (mg)** | 142.7±4.9 | 177.5±7.3 ** | 129.8±5.0 ††† |
| **Heart (mg)** | 149.7±5.3 | 188.8±8.1 ** | 138.9±7.1 ††† |
| **Lung (mg)** | 182.0±21.5 | 247.1±25.7 | 206.0±30.8 $^{ns}$ |
| **Spleen (mg)** | 76.0±3.0 | 98.6±5.7 * | 102.5±6.6 ** |
| **TL (mm)** | 17.54±0.06 | 17.57±0.15 | 17.51±0.15 $^{ns}$ |
| **LV/BW** | 4.29±0.10 | 5.56±0.33 | 4.05±0.30 †† |
| **RV/BW** | 0.35±0.11 | 0.49±0.16 | 0.41±0.14 $^{ns}$ |
| **LA/BW** | 0.43±0.10 | 0.75±0.09 | 0.60±0.13 $^{ns}$ |
| **Ventricles/BW** | 4.95±0.10 | 6.51±0.38 * | 4.81±0.38 †† |
| **Heart/BW** | 5.19±0.11 | 6.94±0.46 * | 5.18±0.51 †† |
| **Lung/BW** | 6.37±0.80 | 9.21±1.18 | 7.97±1.74 $^{ns}$ |
| **Spleen/BW** | 2.66±0.15 | 3.57±0.17 ** | 3.83±0.40 * |
| **LV/TL** | 7.05±0.26 | 8.64±0.38 ** | 6.25±0.24 ††† |
| **LA/TL** | 0.21±0.02 | 0.43±0.09 | 0.33±0.16 $^{ns}$ |
| **RV/TL** | 1.07±0.04 | 1.46±0.10 ** | 1.17±0.07 † |
| **Ventricles/TL** | 8.13±0.26 | 10.10±0.39 ** | 7.42±0.31 ††† |
| **Heart/TL** | 8.53±0.28 | 10.74±0.45 ** | 7.95±0.46 ††† |
| **Lung/TL** | 10.37±1.21 | 14.09±1.49 | 11.87±1.92 $^{ns}$ |
| **Spleen/TL** | 4.33±0.17 | 5.60±0.30 ** | 5.87±0.40 * |

Data are shown as mean ± SEM. BW, body weight; LV, left ventricle; RV, right ventricle; LA, left atrium; RA, right atrium; TL, tibial length. Data are presented as mean ± SEM. Statistical significance was determined by one-way ANOVA with Tukey's multiple comparisons test or Kruskal-Wallis test with Dunn’s multiple comparisons test. n = 8-10 mice per group. */† $p<0.05$, **/†† $p<0.01$, ***/††† $p<0.001$. * vs. PD1$^{f/f}$ wt+IgG, † vs. PD1KO$^{CD8}$+IgG. ns, not significant.

**Supplementary Table S9. Echocardiographic data in experimental groups.**

| Parameters | ♂ $PD1^{f/f}$ wt + TAC 6W (n=6) | ♂ $PD1KO^{FoxP3}$ + TAC 6W (n=8) |
|---|---|---|
| **HR (bpm)** | 510±14 | 510±7 $^{ns}$ |
| **LVESD (mm)** | 4.41±0.22 | 4.34±0.32 $^{ns}$ |
| **LVEDD (mm)** | 5.04±0.17 | 4.99±0.21 $^{ns}$ |
| **LVESV (μL)** | 89.53±9.53 | 89.54±14.67 $^{ns}$ |
| **LVEDV (μL)** | 121.63±9.08 | 120.08±11.49 $^{ns}$ |
| **SV (μL)** | 32.10±2.64 | 30.54±3.40 $^{ns}$ |
| **CO (mL/min)** | 16.41±1.51 | 15.51±1.63 $^{ns}$ |
| **LVAW;s (mm)** | 139.82±15.40 | 1.13±0.04 $^{ns}$ |
| **LVAW;d (mm)** | 1.09±0.05 | 0.91±0.04 $^{ns}$ |
| **LVPW;s (mm)** | 0.86±0.06 | 0.93±0.02 $^{***}$ |
| **LVPW;d (mm)** | 0.79±0.02 | 0.77±0.02 $^{ns}$ |

Data are shown as mean ± SEM. HR, heart rate; LVESD, left ventricular end-systolic diameter; LVEDD, left ventricular end-diastolic diameter; LVESV, left ventricular end-systolic volume; LVEDV, left ventricular end-diastolic volume; SV, stroke volume; CO, cardiac output; LVAW, left ventricular anterior wall; LVPW, left ventricular posterior wall. Data are presented as mean ± SEM. Statistical significance was determined by unpaired two-tailed Student's t-test, Welch's t-test, or Mann-Whitney test. n = 6-8 mice per group. ***p<0.01 vs. $PD1^{f/f}FoxP3^{wt}$ +TAC, ns, not significant.

**Supplementary Table S10. Tissue weight and ratios of male $PD1^{f/f}$ wt and $PD1KO^{FoxP3}$ mice.**

| Parameters | ♂ $PD1^{f/f}$ wt + TAC 6W (n=6) | ♂ $PD1KO^{FoxP3}$ + TAC 6W (n=8) |
|---|---|---|
| BW (g) | 30.4±1.4 | 27.3±1.6 $^{ns}$ |
| LV (mg) | 164.4±17.2 | 161.7±11.6 $^{ns}$ |
| RV (mg) | 25.7±2.4 | 32.6±4.3 $^{ns}$ |
| LA (mg) | 6.0±2.0 | 16.2±4.7 $^{ns}$ |
| RA (mg) | 3.2±0.2 | 3.3±0.3 $^{ns}$ |
| LV/RV | 6.5±0.6 | 5.3±0.4 $^{ns}$ |
| LA/RA | 1.8±0.5 | 4.7±1.4 $^{ns}$ |
| Ventricles (mg) | 190.1±18.9 | 194.3±15.6 $^{ns}$ |
| Heart (mg) | 199.3±20.6 | 213.7±20.0 $^{ns}$ |
| Lung (mg) | 242.0±64.4 | 285.0±54.0 $^{ns}$ |
| Spleen (mg) | 86.1±6.2 | 94.0±5.9 $^{ns}$ |
| TL (mm) | 17.40±0.18 | 17.68±0.10 $^{ns}$ |
| LV/BW | 5.44±0.56 | 6.15±0.68 $^{ns}$ |
| RV/BW | 0.85±0.08 | 1.26±0.21 $^{ns}$ |
| LA/BW | 0.20±0.07 | 0.66±0.20 $^{ns}$ |
| Ventricles/BW | 6.29±0.62 | 7.41±0.88 $^{ns}$ |
| Heart/BW | 6.60±0.68 | 8.19±1.09 $^{ns}$ |
| Lung/BW | 7.99±2.15 | 11.36±2.64 $^{ns}$ |
| Spleen/BW | 2.82±0.14 | 3.47±0.19 * |
| LV/TL | 9.42±0.93 | 9.16±0.68 $^{ns}$ |
| LA/TL | 0.34±0.11 | 0.92±0.27 $^{ns}$ |
| RV/TL | 1.47±0.12 | 1.85±0.25 $^{ns}$ |
| Ventricles/TL | 10.90±1.01 | 11.00±0.91 $^{ns}$ |
| Heart/TL | 11.42±1.10 | 12.10±1.16 $^{ns}$ |
| Lung/TL | 13.74±3.45 | 16.14±3.07 $^{ns}$ |
| Spleen/TL | 4.94±0.33 | 5.32±0.34 $^{ns}$ |

Data are shown as mean ± SEM. BW, body weight; LV, left ventricle; RV, right ventricle; LA, left atrium; RA, right atrium; TL, tibial length. Data are presented as mean ± SEM. Statistical significance was determined by unpaired two-tailed Student’s t-test, Welch’s t-test, or Mann-Whitney test. n = 6-8 mice per group. * $p<0.05$, ns, not significant.

**Supplementary Table S11. Primers for genotyping**

| Targeted alleles | Sequence (5'-3') |
|---|---|
| **PD1$^{flox/flox}$ Forward** | TCACGACTTGTTTTCTTCTGTGC |
| **PD1$^{flox/flox}$ Reverse** | GCAAATTCACCTCAGTCATGGAAG |
| **CD8-Cre transgene-Forward** | CAATGGAAGGAAGTCGTGGT |
| **CD8-Cre transgene-Common** | TGGGATTTACAGGGCATACTG |
| **CD8-Cre wild type-Forward** | CACACATGCAAGTCTAAATCAGG |
| **FoxP3-Cre transgene-Forward** | CAGAAGAACGGCATCAAGGTGAAC |
| **FoxP3-Cre transgene-Reverse** | TTGTTCAGCTTGCACCAGGCA |
| **FoxP3-Cre wild type-Forward** | CCTAGCCCCTAGTTCCAACC |
| **FoxP3-Cre wild type-Reverse** | AAGGTTCCAGTGCTGTTGCT |

**Supplementary Table S12. Primary antibodies used for flow cytometry analysis**

| Primary Antibody | Conjugate | Clone | Vender | Catalog # |
|---|---|---|---|---|
| **CD3e** | BUV737 | 145-2C11 | BD Biosciences | 612771 |
| **CD4** | BUV496 | GK1.5 | BD Biosciences | 612952 |
| **CD8α** | PE-CF594 | 53-6.7 | BD Biosciences | 624296* |
| **CD11b** | BV650 | M1/70 | Biolegend | 101259 |
| **CD11c** | BV711 | N418 | Biolegend | 117349 |
| **CD11c** | PE-Cy7 | N418 | Biolegend | 117317 |
| **CD16/32** | - | 93 | Biolegend | 101302 |
| **CD19** | BUV395 | 1D3 | BD Biosciences | 563557 |
| **CD44** | FITC | IM7 | BD Biosciences | 553133 |
| **CD45** | BUV805 | 30-F11 | BD Biosciences | 748370 |
| **CD62L** | PE-Cy7 | MEL-14 | BD Biosciences | 560516 |
| **CD69** | AF700 | H1.2F3 | Biolegend | 104539 |
| **CD279 (PD1)** | R718 | 29F.1A12 | BD Biosciences | 568618 |
| **CD279 (PD1)** | BV480 | 29F.1A12 | BD Biosciences | 568612 |
| **CXCR3** | BV786 | CXCR3-173 | BD Biosciences | 741032 |
| **F4/80** | BUV563 | T45-2342 | BD Biosciences | 749284 |
| **FoxP3** | PerCP-Cy5.5 | FJK-16s | eBioscience | 45-5773-82 |
| **γδTCR** | BUV661 | GL3 | BD Biosciences | 750410 |
| **Granzyme A** | PE | 3G8.5 | Biolegend | 149704 |
| **I-A/I-E (MHC-II)** | APC-Cy7 | M5/114.15.2 | Biolegend | 107628 |
| **IFN-γ** | FITC | XMG1.2 | Biolegend | 505806 |
| **Ki-67** | RB780 | B56 | BD Biosciences | 568761 |
| **Ly6C** | BV605 | AL-21 | BD Biosciences | 563011 |
| **Ly6G** | AF700 | 1A8 | Biolegend | 12-9668-82 |
| **Ly6G** | BV605 | 1A8 | BD Biosciences | 563005 |
| **NK1.1** | APC | PK136 | eBioscience | 17-5941-82 |

**Supplementary Table S13. Primary antibodies used for Western blots and IF staining**

| Primary Antibody | Vender | Catalog # |
|---|---|---|
| **ANP** | ThermoFisher Scientific | PA5-29559 |
| **CD8α** | Cell Signaling | #98941 |
| **CD45** | R&D System | AF114 |
| **Collagen III** | Proteintech | 22734-1-AP |
| **CXCL10** | Invitrogen | 701225 |
| **Vimentin** | Santa Cruz Biotechnology | sc-7557 |
